# Predicting dynamic expression patterns in budding yeast with a fungal DNA language model

**DOI:** 10.1101/2025.09.19.677475

**Authors:** Kuan-Hao Chao, Majed Mohamed Magzoub, Emily H. Stoops, Sean Hackett, Johannes Linder, David R. Kelley

## Abstract

Predicting gene expression from DNA sequence remains challenging due to complex regulatory codes. We introduce a masked DNA language model pretrained on 165 fungal genomes related to budding yeast that learns conserved regulatory sequence motifs. Fine-tuning the LM on yeast RNA-seq data—including high-resolution transcriptional regulator induction time courses generated in this study—yielded Shorkie, a model that substantially improves gene expression prediction compared to baselines trained without self-supervision. Shorkie identified canonical transcription factor (TF) binding motifs and captured changes in their regulatory contributions across induction time courses. Furthermore, Shorkie accurately predicted variant effects, outperforming leading sequence-to-expression models in *cis*-eQTL classification and achieving high concordance with massively parallel reporter assays. Interpretability analyses revealed Shorkie’s ability to resolve promoter dynamics, splicing signals, and temporal changes in regulatory motif usage. Together, these results support combining pretraining on related fungal genomes with RNA-seq fine-tuning to improve sequence-based prediction of gene expression in budding yeast and provide insights into noncoding variants.

## Introduction

Predicting gene expression levels from DNA sequence is a fundamental challenge in genomics with broad implications for understanding gene regulation and disease. *Saccharomyces cerevisiae* (budding yeast) has served as the premier model for eukaryotic gene regulation, with ∼7,000 genes controlled by hundreds of transcription factors (TFs). Despite decades of work mapping *cis*-regulatory motifs and their regulators^1–13^, quantitative prediction of gene expression from regulatory sequences remains challenging, reflecting the complexity of promoter grammar and its interaction with chromatin and TF activity.

Supervised deep learning can in principle learn regulatory features directly from DNA sequence^14,15^, but faces a fundamental limitation in yeast: the compact 12 Mb genome provides too few independent training examples to support large supervised models without overfitting. Self-supervised DNA language models (LMs) can mitigate this limitation by learning sequence representations from many unlabeled genomes, which can then be transferred to supervised tasks with limited labeled data. Models such as DNABERT^16,17^, GPN^18^, and others^19–27^ demonstrate that masked-token prediction captures conserved regulatory syntax and the locations of genes. Parallel advances in protein LMs^28–32^ further validate self-supervised pretraining for extracting functional patterns from sequence.

Fungal genomics is particularly well suited to this paradigm because many high-quality genomes are available^33^, while matched expression datasets exist for only a handful of species. We therefore asked whether multi-species masked language modeling can provide a useful initialization for yeast regulatory prediction by seeding the model weights with learned conserved features that correlate with gene expression. We also compared transfer to *S. cerevisiae* from four pretraining corpora spanning different taxonomic scopes, reasoning that a trade-off might exist between learning from diverse genomes and from closely related ones.

Here, we systematically explore these questions to improve yeast expression modeling from sequence. We pretrained a BERT-style bidirectional masked LM on curated sets of fungal genomes with varying divergence, from *S. cerevisiae* strains to order- and kingdom-level fungal collections. We found that an LM trained on 165 “moderately diverse” genomes from the Saccharomycetales order performed better at reconstructing *S. cerevisiae* regulatory sequences than LMs trained with either narrower or broader pretraining corpora. We then fine-tuned this model on high-resolution RNA-seq time courses from transcriptional regulator induction experiments in *S. cerevisiae*^34^, plus publicly available epigenomic and transcriptomic data, creating a high-quality expression predictor called **Shorkie**.

By combining evolutionary pretraining with transfer learning, Shorkie outperforms models trained without self-supervision in predicting expression levels and temporal dynamics of held-out genes, and delivers robust variant effect predictions in both *cis*-eQTL classification and massively parallel reporter assays. These findings demonstrate that masked language modeling across a carefully curated collection of fungal genomes, coupled with transfer learning, provides a practical framework for quantitative gene regulation modeling and noncoding variant interpretation in yeast.

## Results

### Yeast language model design and training across evolutionary divergences

We trained a family of masked DNA language models (LMs) using the same core transformer-based architecture while varying only the evolutionary scope of the pretraining corpus (Figure 1). We compared four corpora spanning increasing divergence from the *S. cerevisiae* reference genome (R64): (i) R64 (species-level), (ii) 80 S. cerevisiae strains (strain-level), (iii) 165 Saccharomycetales genomes (order-level), and (iv) 1,341 fungal genomes (kingdom-level). The kingdom-scale corpus (1,341 genomes from Ensembl Fungi) matches the genome collection used by the Species-aware LM of Karollus et al. (2024)^21^, but here we additionally trained the three narrower corpora to enable a controlled evolutionary-scope comparison.

**Figure 1.**
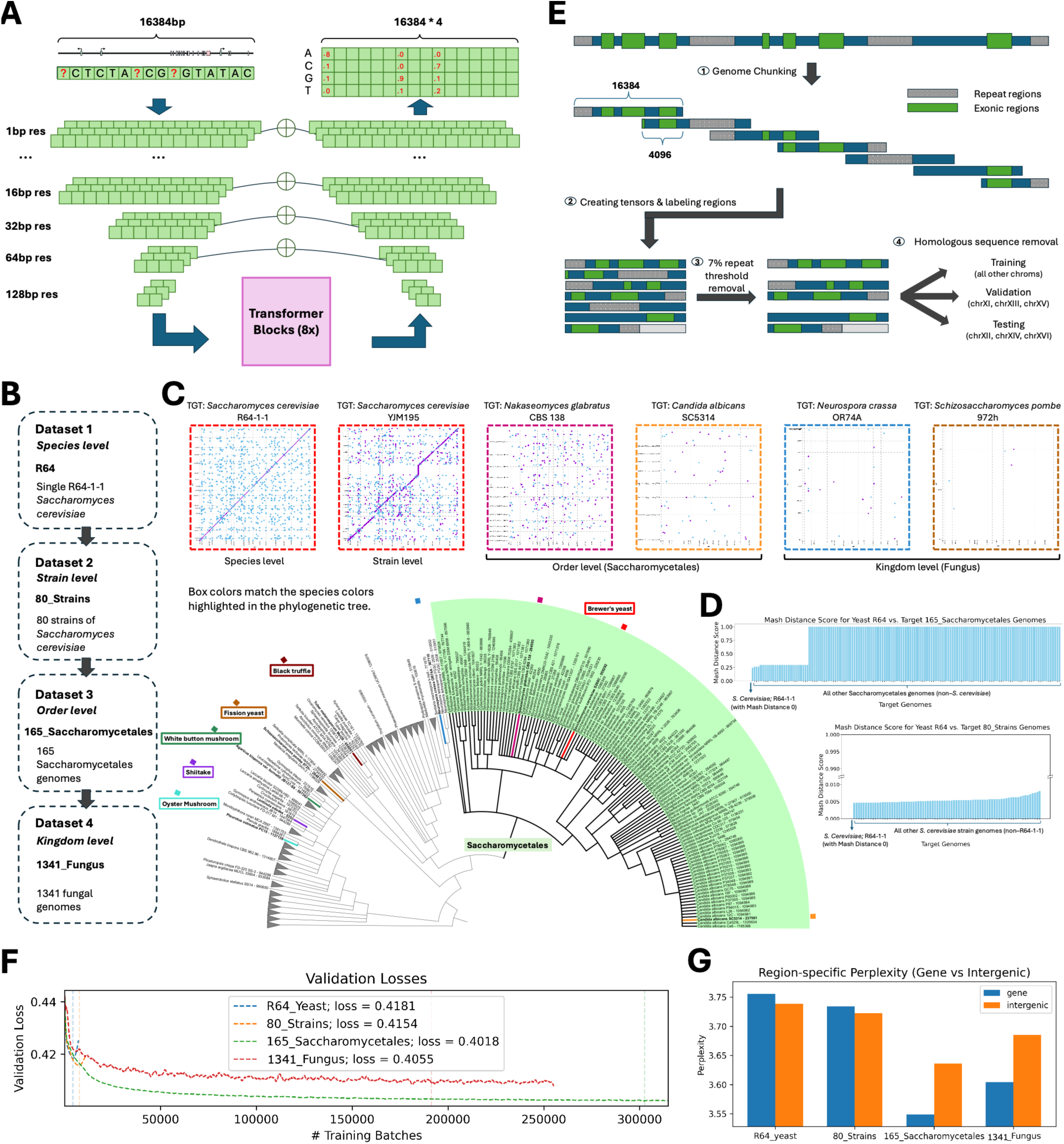
Overview of datasets, preprocessing pipeline, model architecture, and performance metrics for the fungal language model (Shorkie LM). **(A)** Schematic of the Shorkie LM architecture. **(B)** Four datasets employed: single *S. cerevisiae* genome (R64_yeast, species-level), 80 *S. cerevisiae* strains (80_strains, strain-level), 165 Saccharomycetales genomes (165_Saccharomycetales, order-level) with Saccharomycetales highlighted in light green, and 1,341 fungal genomes spanning the kingdom (1341_Fungal, kingdom-level), including common mushrooms such as oyster mushroom (*Pleurotus ostreatus*), shiitake (*Lentinula edodes*), white button mushroom (*Agaricus bisporus*), and black truffle (*Tuber melanosporum*), as well as fission yeast (*Schizosaccharomyces pombe*), and brewer’s yeast (*Saccharomyces cerevisiae*). **(C)** Representative genome distance dot plots for selected genomes from each dataset, with the x-axis representing the R64 *S. cerevisiae* genome and the y-axis representing the comparison genome. **(D)** Mash distance between R64 *S. cerevisiae* genome and genomes in 80_strains and 165_Saccharomycetales. **(E)** Data preprocessing pipeline converting raw genomic sequences into one-hot tensors and region-specific loss weights for Shorkie LM training, validation, and testing. **(F)** Validation loss progression across training steps. **(G)** Comparison of test set perplexity in genic and intergenic regions across four model variants.

Our LM model architecture integrates elements from Enformer^35^ and Borzoi^36^, employing a convolutional tower with subsampling followed by eight self-attention blocks operating at 128 bp resolution. We repeated Borzoi’s U-Net upsampling block^37^ seven times to progressively restore single-nucleotide resolution for masked token prediction (Figure 1A; detailed model configurations in Figure 1-figure supplement 1; Methods).

To characterize the evolutionary scope of our four corpora, we constructed a taxonomic topology with ETE3 using NCBI Taxonomy^38^ and visualized it in iTOL^39,40^ (Figure 1B; see Figure 1-figure supplement 2 for full tree; Methods). We quantified divergence from *S. cerevisiae* R64 using MUMmer dot plots^41^ (Figure 1C) and Mash distances^42^ (Figure 1D). These metrics highlight the spectrum of genomic preservation: closely related strains (e.g., YJM195 at Mash ≈ 0.01) exhibited near-continuous synteny, taxa such as *C. albicans* and *N. glabratus* showed structural fragmentation (Mash 0.25–1), and distant outgroups (*S. pombe*, *N. crassa*) yielded negligible alignments (Figure 1C).

We applied a consistent preprocessing and split strategy across all evolutionary scopes: we prepared genomes for LM training by masking repetitive elements^43–48^ (Figure 1-figure supplement 3A–B; Methods), segmenting into overlapping 16,384 bp windows with 4,096 bp stride, and excluding windows with >7% repetitive content. We assessed gene count per window (Figure 1-figure supplement 4A–B), coding-to-noncoding ratios per window (Figure 1-figure supplement 4C–D), repetitive region distribution (Figure 1-figure supplement 4E–F), and gene annotation completeness (Figure 1-figure supplement 4G–I; Methods). To focus learning on regulatory sequences, we down-weighted the masked-token loss function by 0.1 at annotated exonic (72% of *S. cerevisiae* R64) and repeat-masked (7.39%) positions, multiplying the two factors where both apply (Figure 1-figure supplement 3C; Methods). Training, validation, and test sets were split by *S. cerevisiae* chromosomes. To prevent sequence leakage, the training set included all other genomes only after removing any sequences homologous to the validation or test sets (using minimap2 at a 20% divergence cutoff^49,50^) (Figure 1E; Figure 1-figure supplement 3D–E; Methods).

Among the four pretraining corpora, the 165_Saccharomycetales model achieved the lowest validation loss (Figure 1F) and test perplexity on held-out *S. cerevisiae* chromosomes (Figure 1G). This ranking was also observed across the alternative architectures tested: two residual CNN baselines and a larger U-Net-transformer (Figure 1-figure supplement 5B–D, Figure 1-figure supplement 6). These results identify the Saccharomycetales corpus as the best-performing corpus among the training configurations tested. Therefore, we refer to the selected 165_Saccharomycetales-trained model as “Shorkie LM”.

### Shorkie LM captures regulatory conservation and generalizes across diverse fungi

Transcription factor (TF) binding motifs are fundamental regulatory units^51^, and previous studies demonstrate that masked DNA LMs learn subtle motif co-occurrence patterns^21^. To visualize sequence patterns reconstructed by Shorkie LM, we first computed per-position nucleotide probability distributions from a single unmasked forward pass and compared these probabilities to those produced by the Species-aware LM (SpeciesLM) from Karollus et al.^21^ (Figure 2A; Methods). Visualized as sequence logos, the two LMs yield broadly similar base-probability patterns at the SMT3 promoter, including Poly(dA:dT), Cbf1, Tye7, and Reb1.

**Figure 2.**
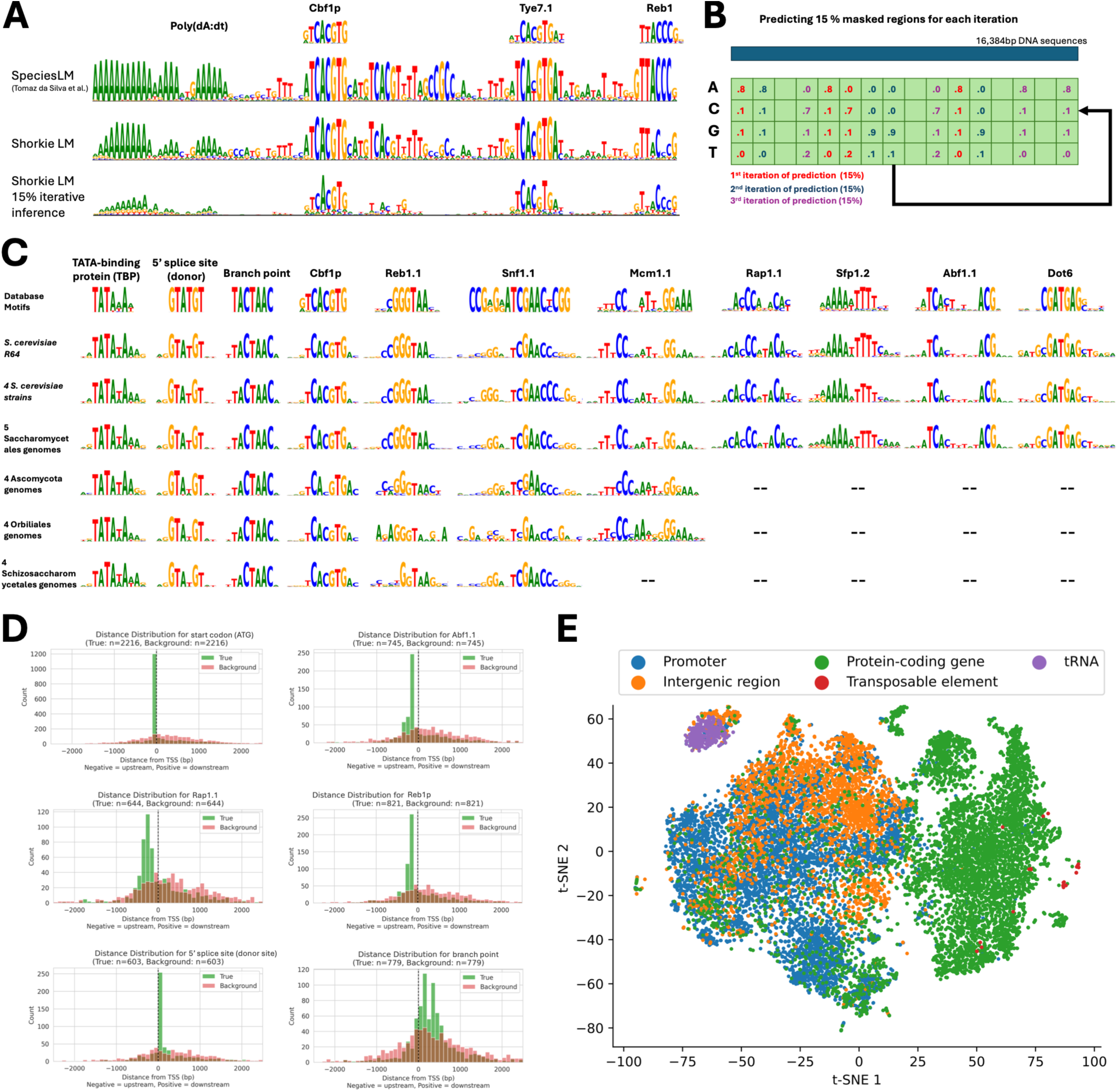
Shorkie LM identifies conserved transcription factor binding motifs across fungal genomes. **(A)** Comparative analysis of SMT3 promoter predictions (chrIV:1,469,090-1,469,198) by Shorkie LM and the Species-aware DNA LM^21,64^, highlighting key motifs including poly(dA:dT), Cbf1, Tye7, and Reb1. **(B)** Position probability matrix (PPM) reconstruction from DNA sequences using the fungal language model Shorkie LM. **(C)** Summary of known motifs detected by Shorkie LM across six datasets: (1) reference *S. cerevisiae* genome; (2) four randomly selected *S. cerevisiae* strains; (3) five genomes from the Saccharomycetales order; (4) four genomes from the Ascomycota phylum; (5) four genomes from the Orbiliales order; and (6) four genomes from the Schizosaccharomycetales order. TF-MoDISco-identified motifs include TATA-binding protein, 5′ splice site (donor), branch point, Cbf1p, Reb1.1, Snf1.1, Mcm1.1, Rap1.1, Sfp1.2, Abf1.1 and Dot6. **(D)** Histograms depicting enrichment of TF-MoDISco-identified motifs upstream of transcription start sites (TSS) relative to background distributions in *S. cerevisiae*, and enrichment of 5’ splice sites (donors) and branch points within genic regions. **(E)** t-SNE embeddings of genomic elements from the first transformer block of Shorkie LM, using 500 bp of native sequence per element and occupancy-weighted pooling.

Because Shorkie LM is trained with random 15% masking, we additionally report a training-consistent Monte Carlo reconstruction to reduce visualization noise: we randomly mask 15% of the positions, predict the masked bases, and repeatedly mask and predict a new set of positions until all positions are covered (Figure 2B; Methods). Averaging over many random masking patterns, the resulting sequence logos lose many spurious low-entropy positions while retaining the core motifs (Figure 2A). We assessed Shorkie LM across six fungal datasets spanning different evolutionary distances (see Methods). Following prediction, we employed TF-MoDISco-lite^52^ for *de novo* motif clustering and matched clusters to yeast motif databases^53–58^ (Methods). Motif conservation varied across evolutionary distance. Shorkie recovered core regulatory motifs and features, including TATA-binding protein (TBP)/TATA elements, start codons, splice sites, and TF binding motifs such as Cbf1, Reb1, and Snf1. Mcm1.1-like motif was not recovered in the Schizosac-charomycetales evaluation set, as this order lacks a direct homolog; this role is instead served by the functionally analogous MADS-box transcription factor Map1^59–62^. Motif conservation declined beyond the Saccharomycetales order, consistent with model training metrics (Figure 1G; Figure 2C). See Figure 2-figure supplement 1 and Figure 2-figure supplement 2 for comprehensive motif discovery results.

To validate biological relevance, we mapped TF-MoDISco-derived motifs onto the *S. cerevisiae* genome, assigning motifs to their nearest genes and computing transcription start site (TSS) distances. Compared with random controls, TBP, Cbf1p, Reb1.1, Mcm1.1, and Snf1.1 showed promoter enrichment (Figure 2D; Figure 2-figure supplement 1). The 5’ splice donor site localized downstream of TSSs, while branch points distributed broadly within genes. Five randomly selected Saccharomycetales genomes produced similar enrichments (Figure 2-figure supplement 2). At the representation level, t-SNE projections of Shorkie LM representations from the first attention layer revealed structure associated with genomic feature annotations using fixed 500-bp native contexts (Figure 2E; Figure 2-figure supplement 3). Furthermore, nucleotide dependency analysis^63^ detected interactions between regulatory elements at selected loci that were absent in an untrained, architecture-matched control and were replaced by diffuse, non-localized dependencies after dinucleotide shuffling (Figure 2-figure supplement 4).

These results show that Shorkie LM recovers conserved regulatory motifs in promoter and splice-site contexts and across distantly related fungi.

### Shorkie: LM transfer learning enables improved gene expression prediction

Building on the strong self-supervised sequence foundation, we developed Shorkie, a supervised model predicting RNA-seq and ChIP-exo/MNase aligned coverage tracks at 16 bp resolution from DNA sequences. Starting with the LM architecture, we removed its final four upsampling layers and added task-specific output heads (Figure 3A; complete configurations in Figure 3-figure supplement 1; Methods). We curated 2,162 experimental tracks for training: 1,128 ChIP-exo^65^, 20 ChIP-MNase^65^, and 1,014 RNA-seq datasets from various yeast isolates^66^. We trained these assays jointly to provide complementary supervision — TF occupancy and nucleosome organization alongside transcript output — reasoning that multi-assay training can improve shared regulatory representations and regularize RNA-seq learning, as demonstrated by ablations in Borzoi^36^.

**Figure 3.**
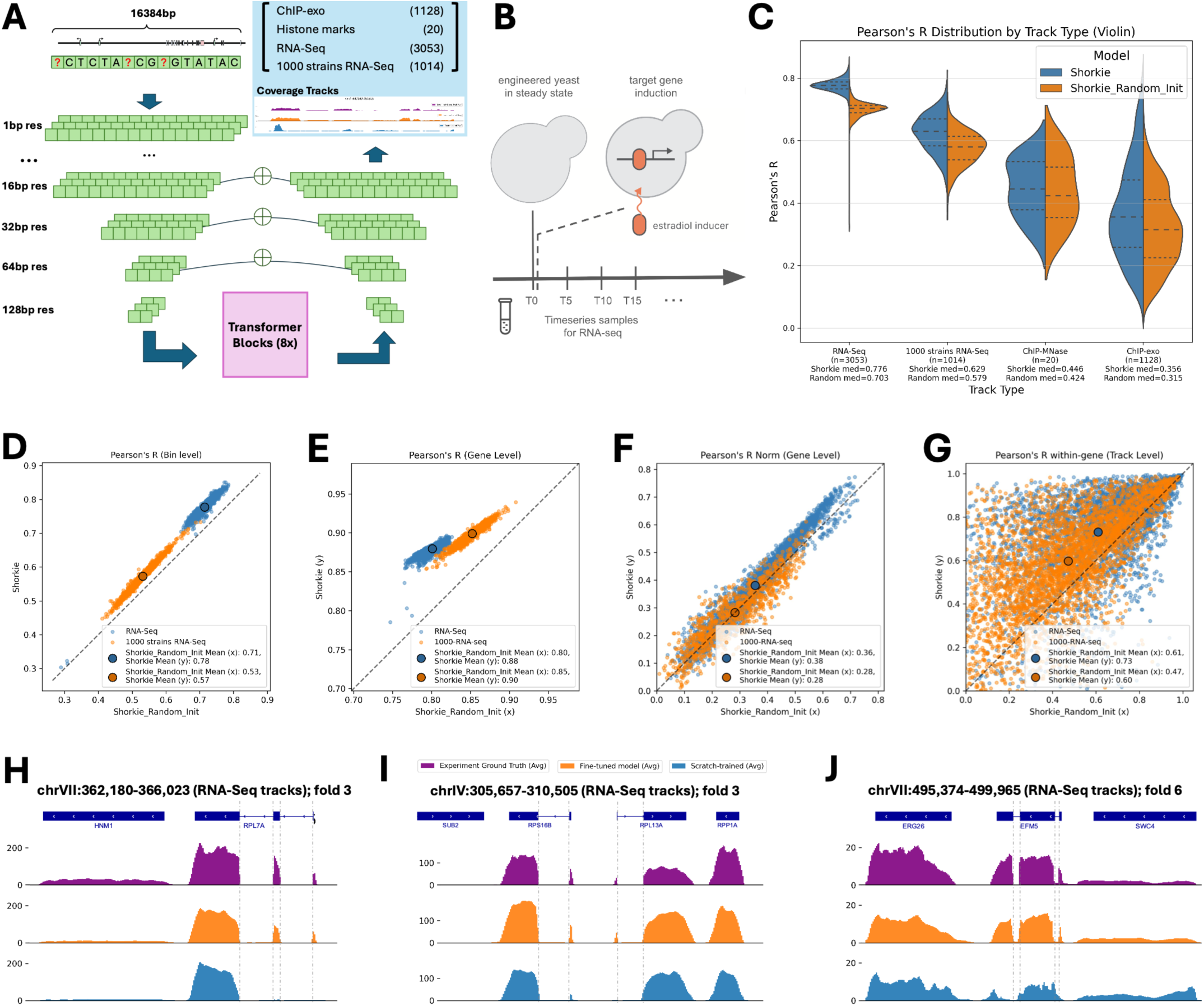
Shorkie architecture and RNA-seq prediction performance across multiple scales. **(A)** Shorkie architecture: U-Net model with eight transformer blocks. All layers inherit pretrained Shorkie LM weights; task-specific output heads (blue) predict perturbation time point RNA-seq (n = 3,053), 1000-strain RNA-seq (n = 1,014), ChIP-exo (n = 1,128) and ChIP-MNase histone marks (n = 20). **(B)** Yeast cells were grown to steady state, as determined by culture density, prior to addition of β-estradiol to the culture and subsequent sampling. **(C)** Distribution of bin-level Pearson’s R on held-out test data for each track type, comparing Shorkie and Shorkie_Random_Init. **(D-G)** Scatterplots comparing Shorkie and Shorkie_Random_Init for RNA-seq tracks at **(D)** bin-level Pearson’s R; **(E)** gene-level Pearson’s R; **(F)** quantile-normalized and mean-centered gene-level Pearson’s R; **(G)** gene-by-gene, track-level Pearson’s R. **(H-J)** RNA-seq coverage snapshots of *S. cerevisiae* test set gene loci: **(H)** chrVII:362,180–366,023 (RPL7A); **(I)** chrIV:305,657*–*310,505 (RPS16B and RPL13A); and **(J)** chrVII:495,374–499,965 (EFM5).

In addition, we generated 3,053 new high-resolution induction RNA-seq time points using a protocol adapted from Hackett et al.^34^ (Figure 3B), bringing the total to 5,215 experimental tracks (Figure 3-figure supplement 2). Key protocols and descriptions of the chemostats (ministat array), culture conditions, library preparation, data pre-processing and quality controls are detailed in Methods (Figure 3-figure supplement 3 and Figure 3-figure supplement 4).

To quantify the benefit of pretraining, we initialized the supervised Shorkie architecture either from random weights (Shorkie_Random_Init) or from masked LMs pretrained at increasing evolutionary scope (R64-only; 80 S. cerevisiae strains; 165 Saccharomycetales genomes, Shorkie_LM; 1,341 fungi), then fine-tuned all variants. Across eight-fold cross-validation, LM initialization improved validation performance and accelerated convergence relative to random initialization. Among candidate LMs, the 165 Saccharomycetales-trained model achieved the highest validation Pearson’s r and was therefore used as the default initialization (Figure 3-figure supplement 5A). Across LM-pretrained initializations, lower held-out S. cerevisiae masked-LM perplexity was associated with better downstream fine-tuning performance (best validation Pearson’s R; Figure 3-figure supplement 5B). To ensure a robust from-scratch baseline (Shorkie_Random_Init), we optimized both its learning rate and model architecture; however, neither approach closed the performance gap with the pretrained model (Methods; Figure 3-figure supplement 6, Figure 3-figure supplement 7, and Figure 3-figure supplement 8).

On the transcriptional regulator induction RNA-seq test data, Shorkie achieved a median bin-level Pearson’s R of 0.78 versus 0.70 for the tuned Shorkie_Random_Init baseline (paired difference across the eight held-out folds +0.063, 95% CI +0.028 to +0.098; Figure 3C; Figure 3-figure supplement 9A) and higher per-track correlations (Figure 3D). The paired difference represents the mean of the per-fold differences in median R, whereas 0.78 and 0.70 are the medians across all test tracks. Induction time-course RNA-seq tracks were predicted more accurately than the 1,000-strain compendium tracks, consistent with greater cross-study technical variation in the compendium (e.g., differences in sequencing depth, read configuration, and batch effects) and the added difficulty of predicting expression across diverse genetic backgrounds. Gene-level aggregation across exon-overlapping bins (Methods) yielded mean Pearson’s R of 0.88 for Shorkie versus 0.80 for the tuned Shorkie_Random_Init (Figure 3E). Normalized gene-level correlations (quantile-normalized per experiment and mean-centered per gene) showed a modest gain that was not statistically significant across folds (Figure 3F; Figure 3-figure supplement 9B). After averaging track-specific performance across tracks for each gene, Shorkie exceeded Shorkie_Random_Init for 75.3% of genes in the induction data and 79.2% in the 1,000-strain data, particularly at higher expression levels (Figure 3G; Methods).

While our genome-wide performance metrics establish Shorkie’s overall predictive advantage, we also examined locus-level predictions and attention patterns at representative genes for interpretability. At held-out loci, Shorkie more faithfully reproduces exon-intron structure and coverage profiles than Shorkie_Random_Init (Figure 3H–J). Furthermore, example self-attention maps around *EFM5* and *RPL7A* illustrate that Shorkie and Shorkie LM place more concentrated attention on discrete intergenic regulatory regions (putative promoters), whereas Shorkie_Random_Init exhibits more diffuse attention (Figure 3-figure supplement 10; Methods).

Together, these genome-wide metrics and locus-level examples show that initializing from the fungal language model substantially improves expression prediction.

### Shorkie transfer learning identifies regulatory motifs in expression predictions

To interpret sequence patterns learned during pretraining versus those used for downstream expression prediction, we used two complementary analyses on promoters. First, Shorkie LM information content compares the masked-token predictive distribution against a genomic background, highlighting enriched/conserved sequence regularities within the multi-species pretraining data. Second, *in silico* saturation mutagenesis (ISM) on the fine-tuned Shorkie model measures the change in predicted RNA-seq coverage following each nucleotide substitution under the supervised objective^67^. Because these metrics quantify different properties, they need not agree. A motif that is highly conserved across species may contribute little to baseline expression in a particular condition, while a sequence feature with strong effects on predicted expression may not be among the most conserved patterns.

We performed ISM on 500-nt promoter windows (−450 to +50 relative to TSS) for three gene cohorts: 137 ribosomal protein (RP), 64 ribosome and rRNA biosynthesis (RRB), and 3,258 additional protein-coding genes. ISM maps for Shorkie and Shorkie_Random_Init were averaged across eight cross-validation folds using baseline *T*_0_ RNA-seq tracks for predictions, and compared to LM-derived information content logos to contrast learned regularities with features affecting model-predicted expression.

In RP promoters like RPL26A (Figure 4A, Figure 4-figure supplement 1), both Shorkie LM and Shorkie recovered the forkhead-binding IFHL motif, typically located ∼50–80 bp upstream of the TSS. This motif is recognized by the winged-helix domain of Fhl1, which upon phosphorylation recruits the coactivator Ifh1, linking TOR/PKA signaling to transcriptional activation^68–70^. Both models also identified the UASrpg element (Upstream Activation Sequence, ribosomal protein genes), bound by the pioneer factor Rap1, which recruits chromatin remodelers, scaffolds Fhl1–Ifh1 complexes, displaces the +1 nucleosome, and establishes nucleosome-depleted regions essential for preinitiation complex assembly^71–74^.

**Figure 4.**
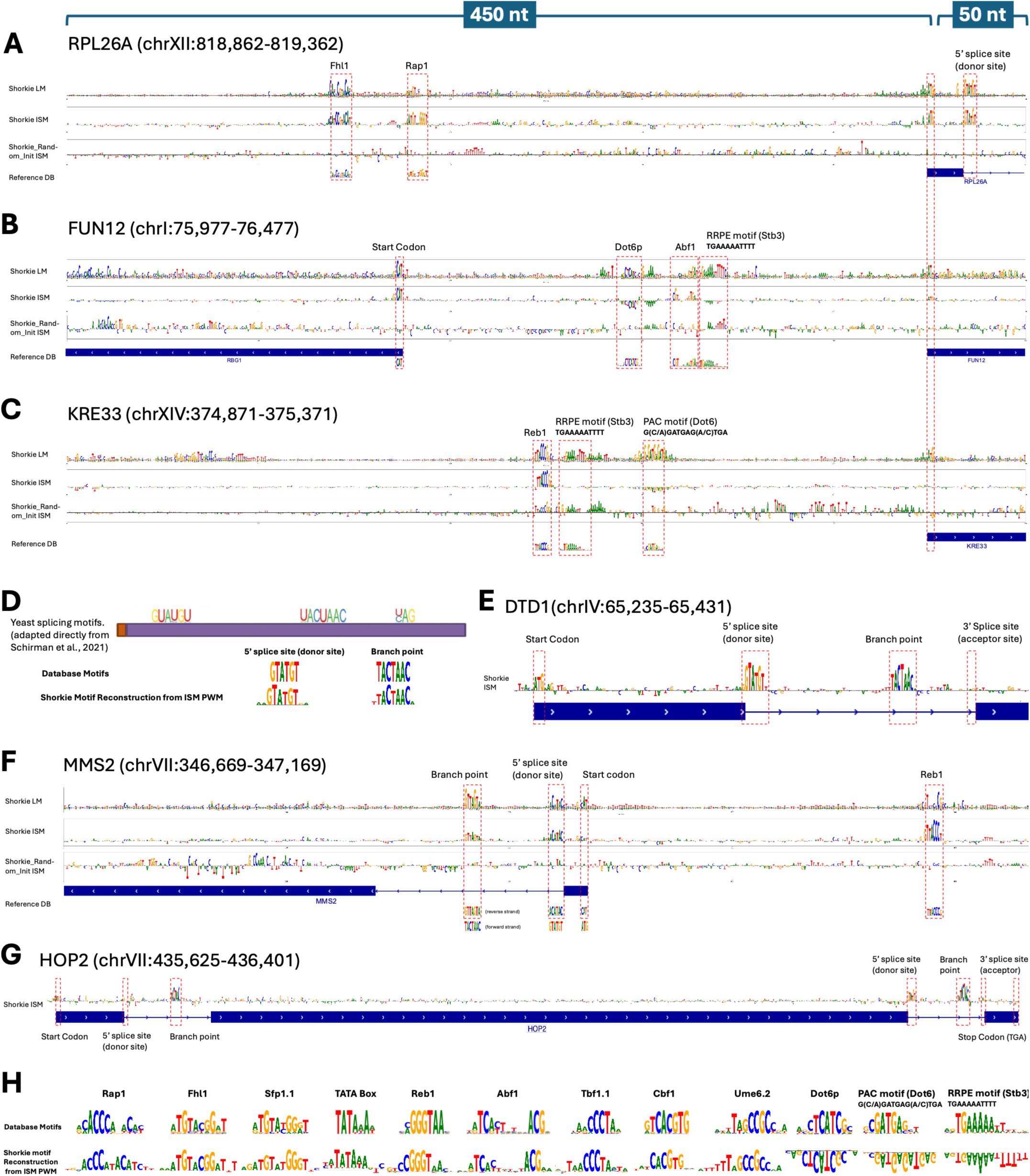
Shorkie uses promoter and splicing motifs learned during pretraining. **(A-C)** Promoter regions (−450 to +50 bp relative to the TSS; 500 bp total) of RPL26A (chrXII:818,862–819,362), FUN12 (chrI:75,977–76,477), and KRE33 (chrXIV:374,871–375,371). Rows 1-3 show DNA logos from Shorkie LM, and ISM maps from Shorkie (fine-tuned) and Shorkie_Random_Init (no self-supervised pretraining). Row 4 shows gene annotations from igv.js^85^. The Shorkie LM PWMs were generated from PPMs (Figure 2A; Methods), whereas the Shorkie and Shorkie_Random_Init ISM maps were produced via an ISM analysis that systematically substituted each nucleotide with the three alternatives. **(D)** Canonical S. cerevisiae splicing motifs—5’ splice site (GUAUGU), branch point (UACUAAC) and 3’ splice site (YAG)—on a schematic synthetic oligonucleotide (purple), adapted directly from Schirman et al. (2021)^81^; below, the corresponding database motifs and the motifs reconstructed from Shorkie ISM maps. **(E-G)** Shorkie ISM maps for splicing motifs in DTD1, MMS2, and the two-intron gene HOP2. **(H)** TF-MoDISco-identified motifs on Shorkie ISM maps: curated yeast database motifs (top) and Shorkie-derived motifs (bottom).

In RRB promoters, such as FUN12 (Figure 4B) and KRE33 (Figure 4C), Shorkie LM captured the RRPE motif (5’-TGAAAAATTTT-3’), bound by the repressor Stb3, and the PAC motif (5’-GCGATGA-GATGAG-3’), recognized by Dot6/Tod6 repressors. These *cis*-elements coordinate rRNA processing and ribosome assembly genes during the cell cycle and stress responses^75–80^. Shorkie recapitulated PAC motifs and additionally detected Abf1 and Reb1 binding motifs. See Figure 4-figure supplement 2 for four additional protein-coding genes.

Within genes, Shorkie LM and Shorkie detected canonical splicing signals (Figure 4D)^81^: in DTD1 (Figure 4E), MMS2 (Figure 4F), and the multi-exon HOP2 (Figure 4G), Shorkie ISM maps delineated the 5’ splice donor sites and branch-point^81–84^. Models were insensitive to acceptor-site mutations—an observation mirrored by Tomaz da Silva et al.^64^. TF-MoDISco analysis recovered additional motifs including TATA, Sfp1, Tbf1, Cbf1, and Ume6 (Figure 4H).

In a paired comparison of ISM maps across 1,685 promoter windows, Shorkie showed significantly higher motif recovery than Shorkie_Random_Init for PAC and RAP1, two of the five candidates with estimated motif-call FDR ≤0.5 (Holm correction across all 144 motifs; Methods; Figure 4-figure supplement 3).

### Shorkie captures dynamic *cis*-regulatory motif changes across time-course TF inductions

Building on Shorkie’s ability to identify static cis-regulatory elements, we investigated temporal motif usage during TF induction. We performed ISM across promoters (−450 to +50 nt relative to TSSs) of Saccharomyces Genome Database (SGD)-curated TF targets^86^ to generate time-resolved maps, computing scores over bins overlapping the target gene’s annotated exons (Methods).

We first examined MSN2, a C2H2 zinc-finger TF that activates ∼200 stress-responsive genes via STRE motifs^87–90^. Shorkie’s ISM maps at the ATG42 promoter revealed progressive STRE sharpening over 0– 90 min (Figure 5A), mirroring RNA-seq fold-changes (Figure 5B; Methods). Euclidean distance heatmaps quantified temporal ISM divergence at this local promoter (Figure 5C). Expanding our evaluation genome-wide across all *S. cerevisiae* genes, we obtained per-track Pearson’s correlations between normalized, mean-centered experimental and predicted RNA-seq expression, with time-point medians ranging from 0.55 to 0.65 during MSN2 induction (Figure 5D). Additionally, genome-wide TF-MoDISco analysis of ΔT ISM maps effectively captured average temporal motif kinetics across these targets (Figure 5E; Figure 5-figure supplement 1A–D show another example at the GLK1 promoter; Methods).

**Figure 5.**
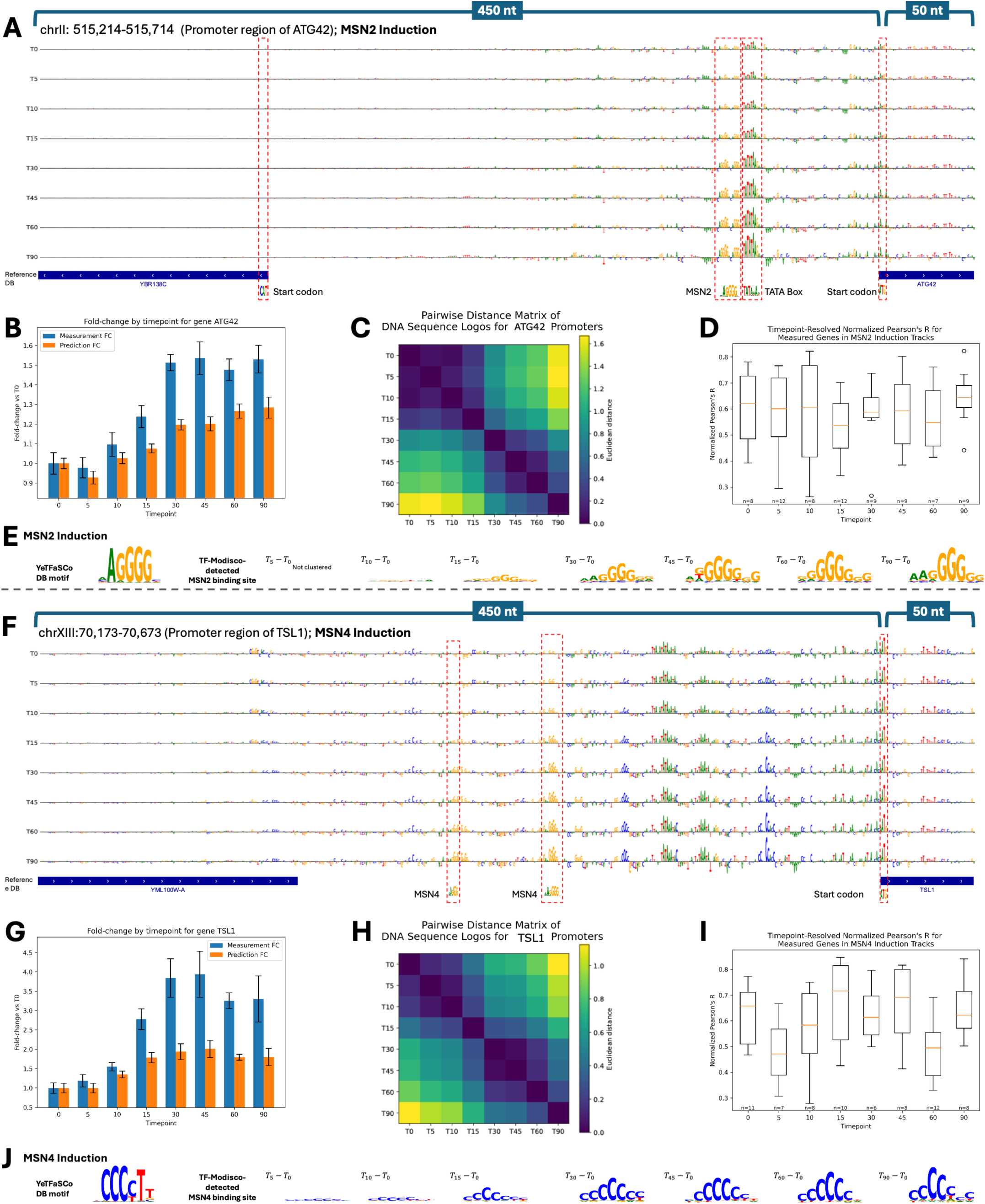
Time-course analysis of stress-responsive transcription factor induction. **(A-E)** MSN2 induction at the ATG42 promoter region (–450 to +50 bp relative to the TSS; chrII:515,214–515,714), sampled at eight time points, including *T*_0_, labeled in minutes. **(A)** Shorkie ISM sequence logos: rows correspond to successive time points (top to bottom), with the bottom row showing the reference. Key TF-binding motifs are annotated. **(B)** Experimental fold-change in reads per million (RPM) (blue) versus Shorkie-predicted signal (orange) across the ATG42 locus at each time point. **(C)** Heatmap of pairwise Euclidean distances between ISM logos, illustrating temporal divergence in motif strength and composition. **(D)** Boxplot of normalized Pearson’s R between experimental and predicted profiles across all *S. cerevisiae* genes for MSN2 induction at each time point. **(E)** TF-MoDISco-identified motifs extracted from ΔT ISM matrices relative to *T*_0_. **(F–J)** MSN4 induction at the TSL1 promoter region (–450 to +50 bp relative to the TSS; chrXIII:70,173–70,673), with panels analogous to **(A–E)**.

We next examined MSN4, an MSN2 paralog rapidly induced upon stress^91^. Shorkie’s ISM analysis at the TSL1 promoter showed similar STRE dynamics, corresponding with RNA-seq fold-changes (Figure 5F–G) and local Euclidean distance heatmaps (Figure 5H). On a genome-wide scale, TF-MoDISco analysis of ΔT ISM maps identified motifs across the analyzed promoter set, while per-track normalized Pearson’s correlations between experimental and predicted RNA-seq expression across all *S. cerevisiae* genes had time-point medians ranging from 0.45 to 0.70 (Figure 5I–J; Figure 5-figure supplement 1E–H, AYR1 promoter).

Finally, we investigated MET4, a bZIP co-activator recruited to E-box motifs (TCACGTG) by cofactors Cbf1 and Met31/Met32^92,93^. Shorkie’s E-box ISM maps inversely correlated with TF induction and were attenuated by cofactor binding, suggesting capture of cofactor-mediated recruitment rather than direct MET4–DNA binding (Figure 5-figure supplement 2; see Discussion).

Beyond representative loci, we quantified genome-wide agreement between predicted and measured RNA-seq log_2_ fold-changes (relative to *T*_0_) across all TF–target gene pairs at each time point, obtaining Pearson’s R = 0.51 – 0.65 from 5–180 min and consistently positive per-TF correlations (Figure 5-figure supplement 3). These results show induction-associated changes in model attribution at regulatory motifs and agreement between predicted and measured expression time courses.

### Shorkie predicts promoter variant effects validated by MPRAs

Massively parallel reporter assays (MPRAs) provide high-throughput measurements of *cis*-regulatory activity, enabling regulatory syntax hypothesis testing and variant interpretation. Because Shorkie was trained on endogenous 16 kb genomic windows rather than short reporter constructs, we do not expect perfect concordance with MPRA measurements. Nevertheless, marginalizing predictions over surrounding context should capture concordant regulatory signal.

We assessed Shorkie using the Random Promoter DREAM Challenge MPRA dataset^94^ containing a held-out set of 71,103 sequences across eight categories: native yeast promoters; random 80-bp oligonucleotides; high-expression sequences; low-expression sequences; sequences challenging prior models; single-nucleotide variant (SNV) perturbations; motif perturbations; and motif-tiling constructs. Each sequence was assayed in ∼100 cells for precise expression estimates^94^. To make marginal predictions with Shorkie, we replaced MPRA constructs upstream of the TSSs of selected “background” genes and averaged predictions across backgrounds.

To characterize positional effects, we selected 10 forward-strand and 12 reverse-strand genes representing low (5–25th percentile), medium (25–75th percentile), and high (75–95th percentile) pre-induction RNA-seq expression quantiles. We systematically inserted MPRA sequences at eleven positions, stepping every 10 bp from 200 to 100 bp upstream of the TSS, and quantified regulatory impact as the log fold-change in downstream gene expression (Figure 6A; Methods).

**Figure 6.**
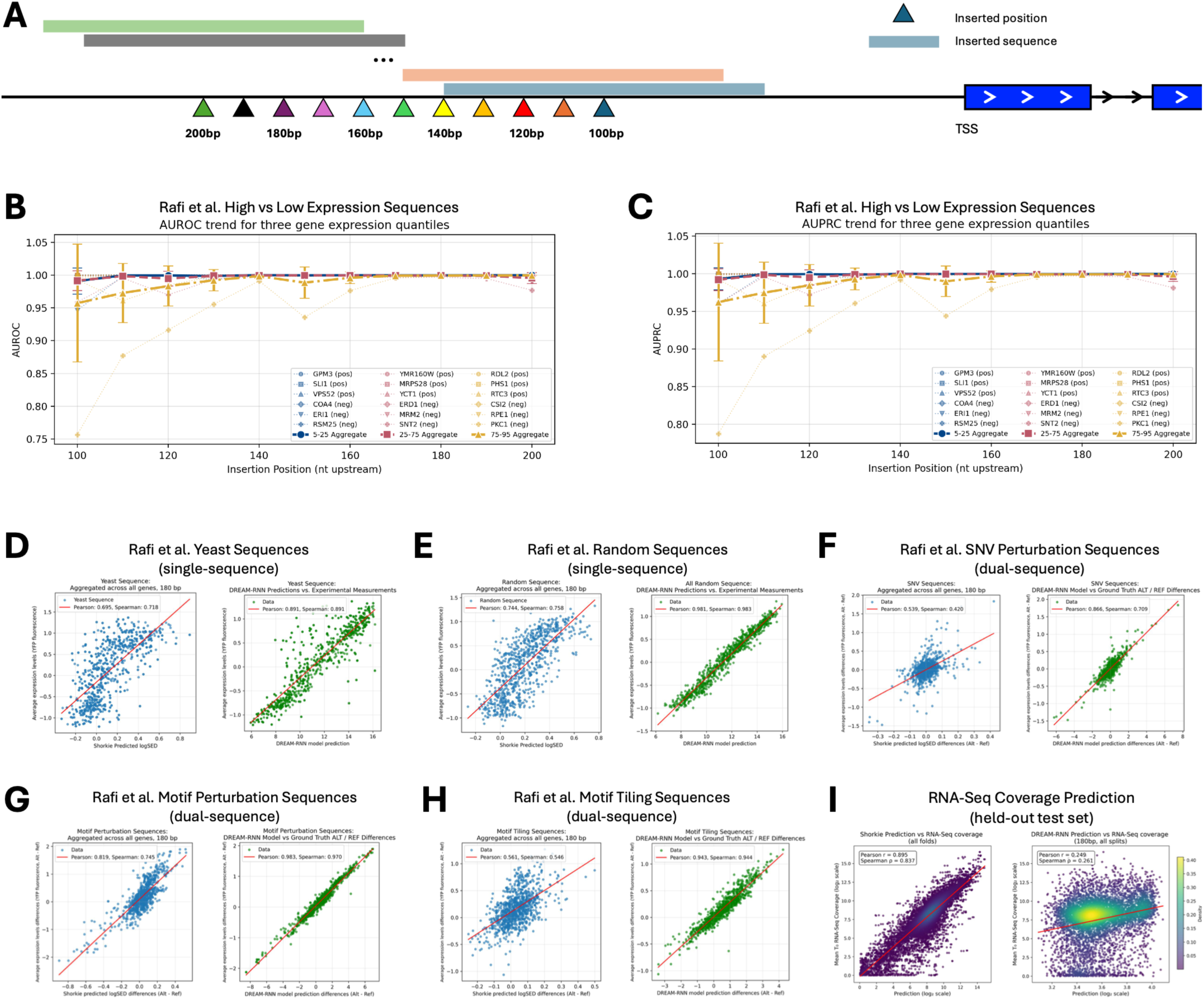
Evaluation of Shorkie’s predictions of promoter variant effects using MPRA data. **(A)** Experimental schematic showing MPRA sequences inserted at positions 100–200 bp upstream of the TSS in 10 bp increments for selected yeast genes. **(B-C)** Classification performance distinguishing high-versus low-expression sequences across upstream insertion sites assessed by **(B)** AUROC and **(C)** AUPRC. Genes were stratified into three RNA-seq expression quantiles (5–25%, 25–75%, and 75–95%); dashed colored lines represent individual genes, and black lines depict mean ± standard error. **(D–E)** Comparison between Shorkie predictions (log fold-change scores) and DREAM-RNN model predictions with experimentally measured expression for **(D)** native yeast sequences and **(E)** challenging sequences. **(F–H)** Model performance evaluated for specific regulatory variant sets: **(F)** single-nucleotide variants (SNV), **(G)** motif perturbations, and **(H)** motif tiling constructs. **(I)** Endogenous RNA-seq coverage prediction (held-out test set) for Shorkie (trained on endogenous genomic tracks) and DREAM-RNN (trained on MPRA), included to illustrate domain specificity and cross-domain generalization.

Across tested genes, high-expression sequences yielded positive log fold-change scores, while low-expression sequences typically produced negative scores (Figure 6-figure supplement 1A–B). Effect magnitudes modestly increased with greater TSS distance. Framing high-versus low-expression sequence prediction as binary classification, Shorkie achieved near-perfect discrimination at each insertion site (AUROC and AUPRC >0.95; Figure 6B–C; Figure 6-figure supplement 1C–D). The position 180 bp upstream was selected for subsequent analyses (Figure 6-figure supplement 2A).

Shorkie’s marginalized predictions strongly correlated with experimental MPRA measurements: Pearson’s R of 0.70 for native yeast promoters, 0.74 for random sequences, and 0.70 for challenging sequences (Figure 6D–E, Figure 6-figure supplement 2B–D). For variant-specific categories, comparing reference-alternate differences, correlations were 0.54 for SNV perturbations, 0.82 for motif perturbations, and 0.56 for motiftiling constructs (Figure 6F–H). As expected, the best-performing MPRA-trained DREAM-RNN outper-formed Shorkie on the DREAM MPRA tasks. Nevertheless, Shorkie—despite not being trained on MPRA data—achieved strong concordance via marginalized-context prediction, demonstrating zero-shot transfer from endogenous genome training to reporter constructs. Furthermore, Shorkie consistently outperformed tuned Shorkie_Random_Init across all categories, with the largest improvements on native promoters and motif perturbations (Figure 6-figure supplement 3). To complement the MPRA benchmarks, we also report performance on endogenous RNA-seq coverage prediction (Figure 6I), highlighting domain specificity: Shorkie generalizes strongly within its training domain (endogenous genomic tracks), whereas DREAM-RNN is optimized for short MPRA sequences. In sum, Shorkie’s endogenous sequence-to-expression predictions are strongly predictive but may also reflect correlative, non-causal associations embedded in genomic sequence.

### Shorkie predicts *cis*-eQTL regulatory impacts

Predicting how regulatory variants alter gene expression is critical for dissecting genetic association mechanisms. We used Shorkie to interpret yeast *cis*-eQTL effects^66,95,96^. For instance, at the OMA1 locus, the eQTL alternate allele reduced Shorkie’s predicted RNA-seq coverage relative to reference (Figure 7A)^66,96^, while at LAP3, the lead eQTL alternate allele increased predicted coverage (Figure 7B). We quantified variant effects via gene expression log fold-change (Figure 7C; Methods).

**Figure 7.**
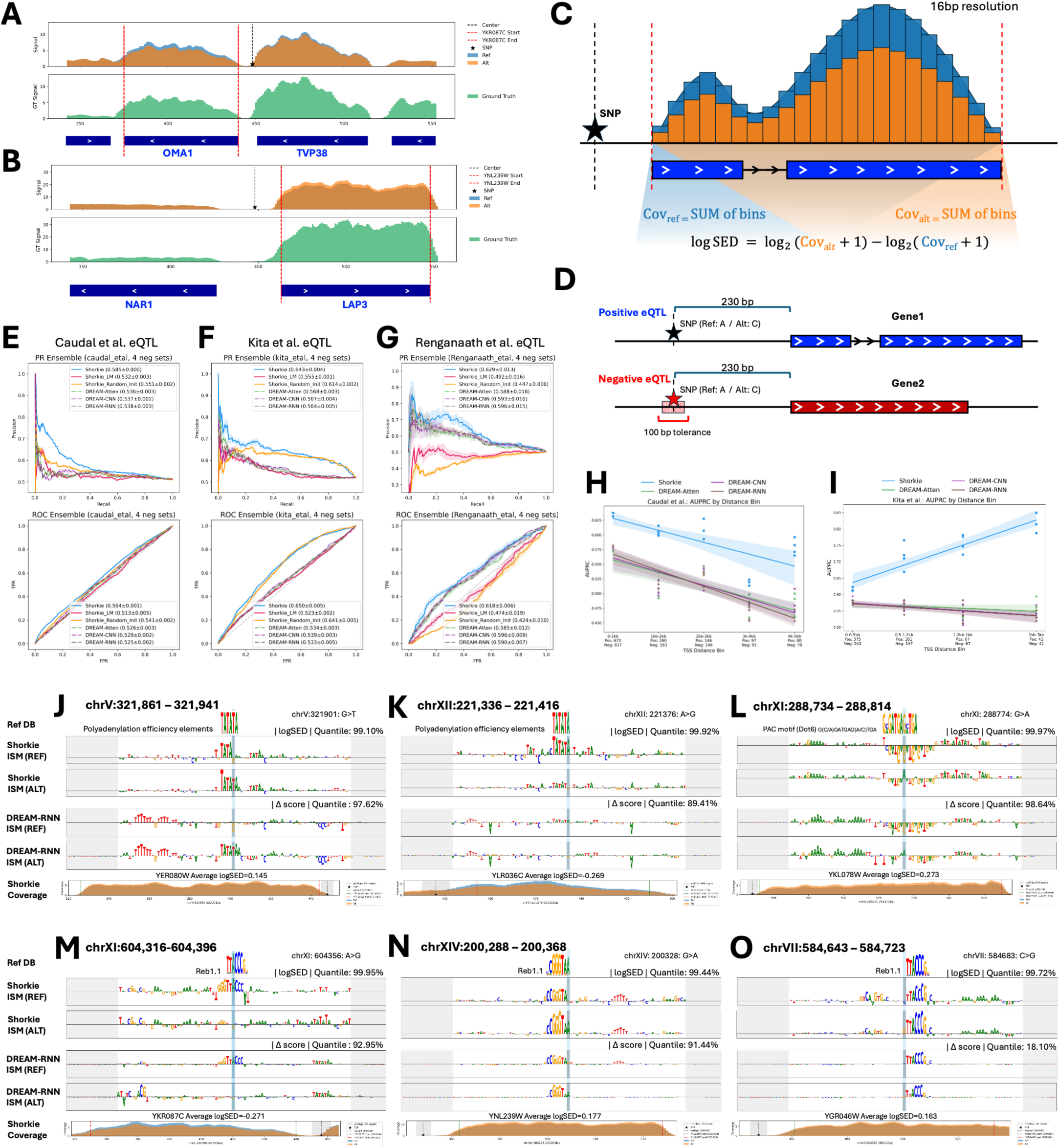
Shorkie accurately predicts cis-eQTL variant effects. **(A)** Positive eQTL example demonstrating reduced expression associated with the alternate allele at OMA1 (chrXI:603,195–604,232). **(B)** Positive eQTL showing increased expression associated with the alternate allele at LAP3 (chrXIV:200,569–201,933). **(C)** Computation of log fold-change expression scores for evaluating variant effects. **(D)** Generation of negative eQTL controls matched by genomic characteristics from ∼1,000 natural yeast isolates. **(E–G)** Precision-recall (PR) and receiver operating characteristic (ROC) curves comparing Shorkie, Shorkie_Random_Init, Shorkie_LM, and DREAM models for Caudal et al. **(E)**, Kita et al. **(F)**, and Renganaath et al. **(G)** datasets. **(H,I)** AUPRC scores by TSS distance bins in Caudal et al. **(H)** and Kita et al. **(I)** datasets. **(J–O)** ISM maps centered on eQTL SNPs, highlighting regulatory motifs identified by Shorkie (SNP in light-blue) and DREAM-RNN, with adaptor sequences in gray.

We benchmarked Shorkie on eQTL datasets from Caudal et al. (1,901 local *cis*-eQTLs from ∼1,000 yeast isolates; Methods)^66,96^ and Kita et al. (683 eQTLs categorized as “Promoter,” “UTR5,” “UTR3,” and “ORF”; Methods)^95^. Additionally, we evaluated Shorkie using a dataset from Renganaath et al. (142 causal cis-regulatory variants within core promoter regions; Methods)^97^. We note that this is not strictly eQTL data, but rather provides MPRA labels based on a prior selection of SNPs with proximity to genes with known allele-specific expression. Negative controls were generated by randomly selecting noncoding SNPs matched by reference/alternate alleles, TSS distance, and minor allele frequency ≥5% (Figure 7D; Figure 7-figure supplement 1; Figure 7-figure supplement 2 A; Methods).

Compared to top DREAM Challenge models (DREAM-CNN, DREAM-RNN, DREAM-Atten)^94^, Shorkie_LM and Shorkie_Random_Init, Shorkie achieved superior ROC and PR metrics for Caudal et al. (Figure 7E; Figure 7-figure supplement 1E), Kita et al. datasets (Figure 7F; Figure 7-figure supplement 2), and Renganaath et al. (Figure 7G) with higher AUROC and AUPRC scores across all TSS distance bins (Figure 7H–I).

ISM analyses of eQTL SNP loci revealed allele-specific remodeling of key *cis*-regulatory elements: creation (Figure 7J) or loss (Figure 7K) of polyadenylation efficiency motifs; disruption of the PAC motif bound by Dot6/Tod6 repressors, correlating with increased expression (Figure 7L); and Reb1 motif alterations either weakening binding (Figure 7M) or strengthening it (Figure 7N–O). See Figure 7-figure supplement 3 for additional examples.

## Discussion

In this study, self-supervised pretraining on Saccharomycetales order genomes performed best of the four evolutionary scopes and the training configurations we tested. Integrating convolutions, transformers, and residual connections within a U-Net architecture, Shorkie LM attains low perplexity on held-out *S. cerevisiae* chromosomes and reconstructs canonical TF binding motifs *de novo* across diverse fungal species. Transfer learning from this foundation into Shorkie substantially improves RNA-seq and ChIP-exo/MNase coverage prediction, raising median bin-level Pearson’s R from 0.70 (tuned random initialization) to 0.78 and gene-level Pearson’s R from 0.80 to 0.88.

Beyond predictive accuracy, TF-MoDISco and ISM analyses show that Shorkie captures both static and dynamic motif signatures. The model recovers static motifs including Poly(dA:dT) tracts, Cbf1, Reb1 and canonical splice-site signals, showing model attribution at known promoter and splice-site motifs. During transcription factor induction time courses, Shorkie dynamically tracks regulatory changes: STRE motif signals sharpen during MSN2 and MSN4 responses, and E-box engagement reflects cofactor-mediated MET4 activation.

However, while ISM and TF-MoDISco correctly identify E-box motifs at MET4 target promoters, they also highlight secondary signals with unclear mechanisms. These features may reflect persistent SCF^Met^^30^-mediated ubiquitination of Met4^98^, transcriptional squelching via Mediator/SAGA cofactor sequestration^99,100^ or feedback from intracellular sulfur metabolites^101^. Disentangling these regulatory layers demands targeted experiments—such as Met4 and cofactor ChIP-seq, ubiquitination-deficient Met4 variants and time-resolved metabolomics—to distinguish direct DNA binding from indirect regulatory cascades.

Despite these biological complexities, Shorkie’s ability to learn regulatory patterns from perturbation experiments opens new avenues for network inference. Genome-wide perturbation screens could systematically map regulator-target relationships through sequence interpretation, while time-series experiments could reveal kinetic parameters governing transcription, mRNA degradation, and splicing. These applications extend Shorkie beyond prediction toward mechanistic discovery.

Realizing these applications requires acknowledging the technical boundaries of the current framework. First, the choice of pretraining corpus matters^102^. The reduced performance of kingdom-level pretraining (1,341 fungi) compared to the order-level corpus (165 Saccharomycetales) may reflect a combination of corpus size, evolutionary divergence, sequence heterogeneity, and increased optimization difficulty. The 165-genome corpus performed best across four tested model capacities (Figure 1-figure supplement 5) and yielded the highest downstream predictive performance (Figure 3-figure supplement 5). We hypothesize that mixing highly divergent species in the kingdom-level corpus may increase gradient variance, destabilizing optimization. Future studies exploring intermediate evolutionary distances (e.g., family-or phylum-level clades) under expanded compute budgets will be critical to cleanly disentangle the effects of phylogenetic divergence from model capacity and training stability.

Second, rigorous dataset quality control remains essential. Although fungi are generally less repeat-rich than mammals, repeat/low-complexity handling remains important because many public fungal assemblies are inconsistently soft-masked, and a small subset of repeat-heavy windows can disproportionately bias the MLM objective. We therefore use a conservative strategy—soft-masking and loss down-weighting at repeat bases, and removing only windows with >7% repeat content. A direct ablation (loss-masking only vs additional window removal) would quantify the benefit in fungi and is left for future work.

Third, assay-specific prediction challenges persist. ChIP-exo was the most challenging assay: predictions track broad trends but systematically underestimate extreme, narrow peaks. This reflects the zero-inflated, heavy-tailed nature of ChIP-exo data and suggests that variance-stabilizing target transforms (e.g., square-root) or distribution-aware losses (e.g., focal loss) could improve performance (Figure 3-figure supplement 11). Similarly, Shorkie RNA-seq time course predictions show compressed dynamic range relative to experimental measurements (Figure 5B,G).

Fourth, although pretraining clearly improves expression prediction, our analyses do not identify which pretrained representations are responsible. The gain could reflect motif detectors, representations of gene structure, higher-order regulatory relationships, or more general inductive biases acquired during pretraining. The motif-specific recovery comparison (Figure 4-figure supplement 3) and nucleotide dependency maps (Figure 2-figure supplement 4) do not establish which representations cause the improvement. Selectively reinitializing the convolutional encoder, transformer blocks or decoder while retaining the remaining language-model weights could help localize the transferred information, although such experiments may be difficult to interpret if this information is distributed across layers.

Ultimately, yeast exemplifies where self-supervised pretraining excels. With a ∼12 Mb compact genome, *S. cerevisiae* cannot support large-scale supervised learning from random initialization alone. MPRAs provide large-scale experimental measurements on synthetic constructs but sacrifice native genome context, creating a tradeoff between scale and biological authenticity. De Boer and Taipale have argued that training models on synthetic sequences rather than endogenous genomes can better isolate the contributions of individual regulatory elements by reducing genomic confounding and spurious correlations^103^. In this light, our evolutionary pretraining strategy is complementary: it leverages abundant related genomes to learn representations grounded in native promoters and genomic context, while we note that endogenous sequence-to-expression prediction is robustly predictive but may reflect correlative, non-causal sequence associations embedded in the genome. Future work may explore hybrid strategies that combine synthetic sequence designs with endogenous, context-rich training to exploit the strengths of both approaches, including evaluating whether self-supervised genomic pretraining can improve reporter-based models like DREAM-RNN.

We propose that genome size and label abundance jointly determine when pretraining yields benefit. Small-genome organisms like yeast — where genome size limits the number of independent training examples and condition-specific genome-wide labels remain scarce — stand to benefit most from phylogenetically in-formed pretraining. Large-genome species (like mammals) with extensive experimental resources may show smaller pretraining gains, though benefits could persist in data-sparse contexts and cross-species transfer. Systematic studies varying taxonomic scope, phylogenetic distance, and experimental data volume will define the boundaries of this pretraining regime.

Our results show that masked nucleotide pretraining on related genomes improves expression prediction in a small-genome, data-limited setting. Shorkie provides a starting point for modeling gene regulation in yeast, and the same approach may extend to other compact-genome organisms.

## Online Methods

### Fungal genome and annotation download, filtering, and biotype partitioning

We downloaded the main species table containing accession IDs, species names, and assembly and database identifiers from Ensembl Fungi release 59 (https://ftp.ebi.ac.uk/ensemblgenomes/pub/release-59/fungi/species_EnsemblFungi.txt) and used it to retrieve all genomic and annotation data. For each taxon, we retrieved the unmasked genome FASTA file and its XML metadata, plus the corresponding GTF annotation file.

We filtered each genome FASTA by parsing its XML to determine assembly level (“chromosome” vs. “scaf-fold”). We retained both chromosome- and scaffold-level assemblies, discarding contigs shorter than 32,768 bp. We renamed remaining contigs according to a unified convention (chrI through chrXVI for yeast, original names otherwise). We produced a cleaned species manifest listing assembly levels, chromosome counts, and total base counts for all retained genomes. We partitioned each annotation file into subsets based on gene biotype (e.g. protein-coding, non-coding, rRNA, etc.).

### Shorkie LM model architecture

Shorkie LM is a U-Net transformer-based model that processes 16,384-bp genomic windows and outputs a probability distribution over the four nucleotides (A, C, G, T) at each position (Figure 1-figure supplement 1). It contains 13,665,828 parameters (13,651,812 trainable; 14,016 non-trainable).

### Encoder (down-sampling) path

The encoder begins with a 1D convolution (kernel size = 11, filters = 96), projecting the input tensor to a feature map of shape (16,384 × 96). Seven successive residual down-sampling stages follow. At stage *i*, the feature map is passed through a residual convolutional block: batch normalization → Gaussian Error Linear Unit (GELU) activation^104^ → Conv1D (kernel size = 5; filters = *C_i_*) → dropout (p = 0.05) → learned residual scaling. The block output is added to its input, and a MaxPooling1D layer (pool size = 2) halves the sequence length. Channel widths increase across stages as *C_i_* ∈ [96, 128, 160, 192, 256, 320, 384], while the sequence length decreases by 2 × each stage from 16,384 (1-bp) to 128 (128-bp).

### Transformer bottleneck

At 128 bp resolution, the (128 × 384) feature map was fed into a stack of eight Transformer blocks to capture long-range dependencies. Each layer uses LayerNorm, then multi-head self-attention (model dim = 384; 4 heads; key dim = 64) with residual dropout (p = 0.2), followed by a two-layer position-wise feed-forward network (Dense → Dropout → ReLU → Dense → Dropout) with a residual connection. This bottleneck integrates context across the full input span.

### Decoder (up-sampling) path and output

The decoder mirrors the encoder in seven up-sampling stages. Each stage applies BatchNorm and GELU, a channel-preserving projection (Dense 384→384), and UpSampling1D (size = 2) to double the sequence length (e.g., 128 → 256 →…→ 16,384). The up-sampled features are merged with the corresponding encoder features via U-Net–style skip connections to restore fine-grained detail, then refined by a depthwise-separable convolution (kernel size = 3; filters = 384), BatchNorm, and GELU. A final 1×1 Conv1D (filters = 4) followed by softmax yields a per-position probability distribution over {A, C, G, T}. (Nearest-neighbor upsampling follows the UpSampling1D definition.)

### Taxonomic tree construction and distance estimation among fungal genomes

#### NCBI taxonomy tree creation

We converted all species names to NCBI Taxonomy identifiers (TaxIDs) with the ETE Toolkit Python API ^38^. To extract a pruned NCBI taxonomy tree connecting our taxa of interest, we ran the ete-ncbiquery command-line module with our TaxID list and requested Newick-formatted output. This step pruned all non-target lineages while preserving branch lengths and hierarchical relationships. We loaded the resulting Newick file into ETE to verify the expected taxonomic groupings and confirm presence of every target species in NCBI. For visualization, we uploaded the pruned tree to the Interactive Tree Of Life (iTOL v5) web server^39,40^ and overlaid custom annotations (e.g., colored clade highlights) using iTOL dataset templates (Figure 1-figure supplement 2).

#### Pairwise genomic distance estimation using alignment- and sketching-based methods

We quantified genomic divergence of the 1,341 fungal assemblies relative to the *S. cerevisiae* R64 reference using complementary alignment-based and sketching-based methods. For the alignment-based approach, we ran MUMmer4’s nucmer^41,105^ with 40 CPU threads to align each cleaned FASTA against the R64 reference, producing a delta file of all maximal unique matches. We then extracted detailed alignment statistics with `show-coords-lcr`, yielding tables of reference and query start-end positions, block lengths, percent identity, and coverage for every alignment block. To visualize large-scale synteny and structural variation, we generated dot plots for each genome pair using MUMmerplot. These plots enable rapid inspection of collinearity breaks, inversions, and translocations across the fungal assemblies.

For rapid, alignment-free genomic distance estimation, we applied two sketching-based tools: Dashing2^106^ and Mash^42^ on the same genome pairs. Dashing 2 leverages the SetSketch data structure^107^ (using a truncated logarithm of hashed k-mers) together with ProbMinHash^108^ for multiplicity-aware sketching. Mash^42^ employs classical MinHash^109^ on each genome’s k-mer set (default k = 21, sketch size s = 1,000) to estimate Jaccard similarity^110^.

### Shorkie LM data preprocessing

#### Repetitive region detection and masking

Most fungal assemblies in the Ensembl database lack adequate soft-masking of repetitive elements. To address this, we retrieved 1,501 genomes from the Ensembl FTP site and implemented a two-tiered repeat-masking pipeline. First, we generated a *de novo* repeat library for each genome using RepeatModeler v2.0^44^, which integrates multiple discovery algorithms, RepeatScout, RECON, and LTR_retriever^44,47,48^, to capture both interspersed and structural elements unique to each assembly, and then merged the resulting consensus sequences with curated entries from Dfam^46,111^. Next, we ran RepeatMasker against this custom library (plus standard repeat databases), employing RMBlast, a RepeatMasker compatible version of the standard NCBI blastn program (https://www.repeatmasker.org/rmblast/), for sensitive and high-throughput alignments. We enabled the “`-xsmall`” option to soft-mask repeats in lowercase thereby preserving original sequence length and coordinates and the “`-gff`” flag to output annotations in GFF3 format. Finally, we applied the DUST algorithm (https://meme-suite.org/meme/doc/dust.html) via the MEME suite^43^ (default cutoff score threshold 20) to soft-mask residual low-complexity regions (Figure 1-figure supplement 3A). Of the 1,501 downloaded assemblies, 160 failed to complete this masking workflow or produced unusable masking out-puts due to assembly/format issues; these were excluded. The remaining 1,341 genomes were successfully masked and used for further preprocessing. We validated the workflow by comparing Ensembl’s original soft-masked regions with our custom masks in six representative assemblies and observed strong concordance (Figure 1-figure supplement 3B). We then partitioned each genome into 16,384-bp windows with a 4,096-bp stride and applied a 7% repeat-content threshold for quality control, excluding ∼20% of windows from the training, validation, and test sets (Figure 1-figure supplement 3C).

#### Homologous and paralogous sequence removal between training, validation and test sets

To guard against data leakage and ensure truly independent evaluation splits, we developed a three-step homology-filtering pipeline that removes any training sequences sharing appreciable similarity with those in our validation or test sets^50^. First, we aligned every training sequence against both the validation and test sets using Minimap2 (v2.28-r1209; assembly-to-assembly mode, “`-x asm`”)^49^ to produce PAF files that report, for each query, the coordinates and match statistics of all detected alignments. From each PAF file, we extracted two metrics for every query-target pair:

• Coverage = matching bases (PAF field 10) / query length (PAF field 2)

• Identity = matching bases (PAF field 10) / alignment block length (PAF field 11)

These ratios quantify how much of a training sequence overlaps with other splits and how similar those overlaps are. Finally, we removed any training sequence for which both coverage ≥ 5 % and identity ≥ 30 % in any alignment, yielding a leakage-free training corpus (Figure 1-figure supplement 3D). The alignment scatter plots for train-test and train-validation splits for each dataset are shown in Figure 1-figure supplement 3E.

#### Evaluating fungal genome annotation completeness using BUSCO

We first retrieved and unpacked the BUSCO v5 fungal lineage dataset (fungi_odb10, 758 orthologs; 2024-01-08; https://busco-data.ezlab.org/v5/data/lineages/fungi_odb10.2024-01-08.tar.gz) to establish a consistent benchmark. For each of the 1,341 assemblies, we extracted all proteins from the Ensembl Fungi release 59 GFF annotation and its corresponding genome using gffread^112^. We then ran BUSCO^113^ in protein mode (`-m proteins`) on each proteome. BUSCO classifies each ortholog as “Complete (single-copy or duplicated)”, “Fragmented”, or “Missing”, providing a standardized completeness score across all annotations (Figure 1-figure supplement 4G–I).

#### TFRecord generation of one-hot encoded genomic windows with exon and repeat masks

We loaded the repetitive, homologous, and paralogous-cleaned 16,384 bp windows with pysam (http://code.google.com/p/pysam/). For each window we: (1) one-hot encoded the DNA sequence; (2) computed exon masks by projecting the exons of all annotated genes onto the window and trimming 2 bp from each exon boundary within the window (the “chew” region); (3) derived repeat masks by flagging lowercase bases; and (4) assigned a species index. We flattened the resulting arrays (“sequence”, “exon_mask”, “repeat_mask”, “species”) to bytes and serialized them as TensorFlow Example protocol buffers. Finally, we wrote ZLIB-compressed TFRecord shards containing 32 examples each.

### Shorkie LM training

We trained and evaluated Shorkie LM using the ZLIB-compressed TFRecord shards described above via a custom version of the baskerville API, called baskerville-yeast (https://github.com/calico/baskerville-yeast). To avoid leakage across closely related sequences, we defined validation and test splits using S. cerevisiae chromosomes only (chrXI, chrXIII, and chrXV for validation; chrXII, chrXIV, and chrXVI for test). The training set comprised all remaining sequences from multiple species after filtering homologous regions, with the larger dataset constructed as a strict superset of the smaller dataset. Notably, the 80 *S. cerevisiae* strain sequences were excluded from these datasets; including them would result in an 80-fold oversampling of the *S. cerevisiae* genome, causing the model to overfit rather than learn broadly across other fungal species during pretraining. Training proceeded in epochs that sampled up to 150 minibatches (batch size = 8) from an in-memory shuffle buffer of 256 records, with full reshuffling between epochs.

At each step, we masked 15% of positions per sequence (*m* = ⌊0.15 × 16,384⌋ = 2,457). Following the BERT protocol^114^: 80 % of masked sites were properly masked, 10 % were substituted by a random nucleotide, and 10 % were left unchanged to prevent the model from over-relying on the mask token distribution. To exploit the fact that double-stranded DNA is symmetric under reverse complementation, we applied reverse-complement augmentation to each input sequence with probability 0.5, thereby encouraging the model to learn strand-invariant features, a strategy shown to substantially improve performance^115,116^.

We computed categorical cross-entropy over masked positions only and weighted each masked position by genomic region: 1 outside exons and repeats, 0.1 within annotated exons or repeat-masked regions, and 0.01 where both overlap. This down-weighting focuses the model on regulatory sequence, following the success of this strategy for repeats in prior work^64^. We trained the model using the Adam optimizer^117^ (learning rate = 1 × 10⁻⁴; β_1_= 0.7; β_2_= 0.9), global clip-norm = 0.1, and a linear warmup over the first 20,000 steps.

Training proceeded for up to 10,000 epochs (minimum = 100; maximum = 10,000), each capped at 150 steps, with early stopping according to the validation loss (patience = 1,000 epochs). At the end of each epoch, we averaged validation loss over five independent mask-and-predict passes per example (repeat_eval = 5) to reduce sampling noise. We retained the checkpoint with the lowest validation loss as the final model.

### Shorkie LM evaluation

We evaluated held-out test windows using the same masking objective as training via a Monte Carlo masking scheme. For each test sequence, we iteratively masked 15 % of positions (2,457 positions) and predicted them until all positions had been covered. We specified the `--rc` flag to average predictions from both the forward sequence and its reverse complement. The resulting per-position nucleotide probabilities quantify model uncertainty/enrichment under the MLM objective and are used to construct probability and information-content matrices for motif visualization. We then computed the per-position cross-entropy loss (Equation 1) for each 16k window *s* as:

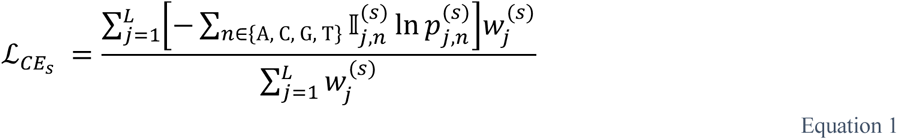

where *L* =16,384 is the window length, 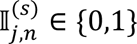 ∈ {0,1} is the one-hot indicator for base *n* at position *j* in window *s*, 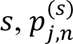 is the model’s predicted probability for base *n*, and 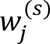 is a position-specific weight (exon/repeat scaling) at that position. The global test loss is the average over all *N* test sequences:

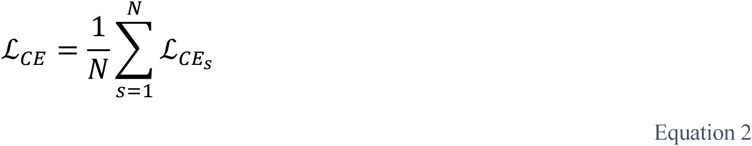

Perplexity is then defined as

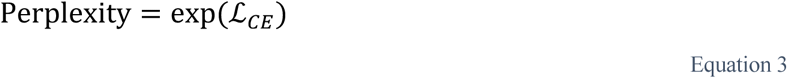

Finally, we calculated taxon-specific evaluation metrics by summing segment losses for each species and dividing by the number of segments for that species, yielding a species-level loss. In addition to loss and perplexity, we computed and stored a position probability matrix (PPM) for each sequence (`x_pred`), alongside one-hot inputs (`x_true`), species labels, and scaling weights for each base pair in a pred.npz file. Finally, we converted these PPMs into information content matrices (ICMs)^118,119^ to facilitate the identification of TF binding sites.

### Computation of Shorkie LM’s information content matrix

Let 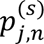 be the PPM probability for nucleotide *n* ∈ {A, C, G, T} at position *j* in the 16,384 bp window *s*. We transform this PPM into an ICM via the following steps. To avoid zeros, we first add a small pseudocount *ε*, yielding

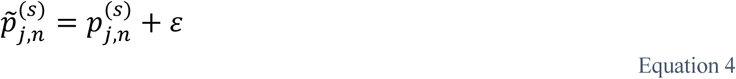

and then renormalize across nucleotides so that

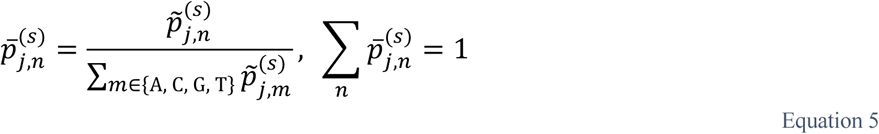

The Shannon entropy at position *j* is

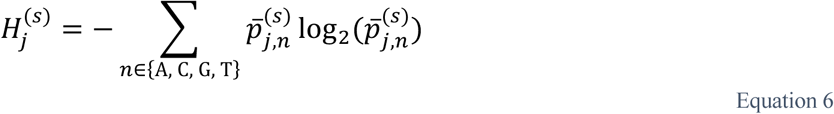

with larger 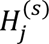 indicating greater nucleotide diversity (i.e. lower conservation). We then define the perposition information content as

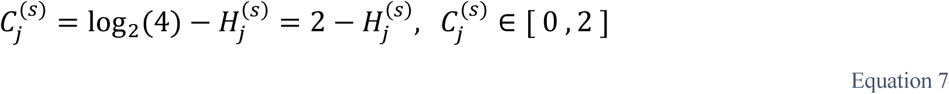

For logo visualization, each nucleotide’s column height is set to

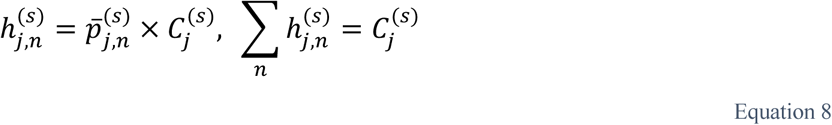

Stacking letters in ascending 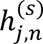 order produces a DNA logo whose total column height reflects the information content, 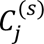, and whose individual letter heights encode the normalized nucleotide probabilities,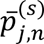

### Constructing a DNA logo of the SMT3 promoter region using SpeciesLM

We adapted the Python notebook workflow of SpeciesLM^64^ (https://github.com/gagneurlab/dependen-cies_DNALM/blob/main/compute_and_visualize_dep_maps.ipynb). First, we loaded a pretrained BERT-style masked-language model (`BertForMaskedLM`; “johahi/specieslm-fungi-upstream-k1”) and its `AutoTokenizer` via Hugging Face Transformers^120^. Following the notebook, we applied this model to the 1 kb upstream region of SMT3 in *S. cerevisiae*. For each sequence, we tokenized the nucleotides and pre-pended a proxy-species identifier. At inference, a softmax produced per-position base-probability distributions. We then computed per-position information content (IC) from the reference (unmutated) probabilities against the yeast genomic background. Finally, we generated a sequence logo by setting the height of nucleotide *b* at position *j* to its predicted probability multiplied by the IC (Equation 8), so total stack height reflects conservation and relative letter heights encode base preferences.

### Six genomic datasets and PPM construction for Shorkie LM evaluation

We evaluated Shorkie LM across six diverse genomic datasets:

1. *S.* cerevisiae reference genome (R64).
2. Four randomly selected S. cerevisiae strains: S. cerevisiae YJM1202, YJM1400, YJM555, and YJM984.
3. Five genomes within the Saccharomycetales order: Candida albicans, Eremothecium gossypii FDAG1, Kluyveromyces lactis str. NRRL Y-1140, Komagataella phaffii CBS 7435, and Nakaseomyces glabratus.
4. Four genomes from the broader Ascomycota phylum: Aspergillus fumigatus, Neurospora crassa, Penicillium chrysogenum str. P2niaD18, and Tuber melanosporum.
5. Four genomes from the Orbiliales order: Arthrobotrys flagrans str. CBS H-5679, Arthrobotrys oligospora ATCC 24927, Dactylellina haptotyla CBS 200.50, and Drechslerella stenobrocha 248.
6. Four genomes from the Schizosaccharomycetales order: Schizosaccharomyces cryophilus, Schizosaccharomyces japonicus, Schizosaccharomyces octosporus, and Schizosaccharomyces pombe.

Except for *S. cerevisiae* and other Saccharomycetales genomes, all others were held out during training. We then applied the approach described in the “Shorkie LM Data Preprocessing” section to segment genomes into 16,384 bp windows, create ZLIB-compressed TFRecord shards, and predict with Shorkie LM.

### Motif discovery with TF-MoDISco-Lite

#### TF-MoDISco run on the *S. cerevisiae* genome

To identify salient sequence motifs captured by the pretrained Shorkie LM, we applied TF-MoDISco-Lite (https://github.com/jmschrei/tfmodisco-lite)^52^ to attribution maps derived from the model’s per-base predictive distributions. We first collected Shorkie LM evaluation outputs containing the one-hot encoded input sequence (`x_true`) and the model-predicted nucleotide probabilities (`x_pred`) for 16,384-bp windows.

We converted predicted probabilities into per-base contribution scores as follows. Let *p_i_*_,j_ denote the predicted probability for nucleotide *i* ∈ {A, C, G, T} at position *j* within a window of length *L*. We added a small pseudocount *ε* = 1 × 10⁻⁴ for numerical stability and computed a per-window background frequency for each nucleotide

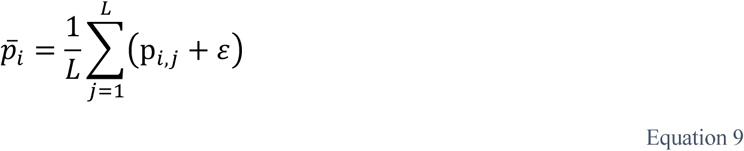

We then transformed probabilities into an information-weighted score:

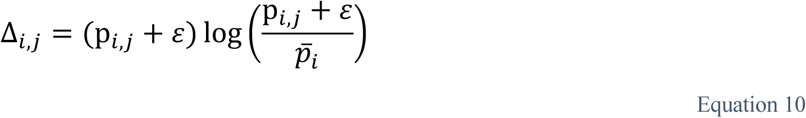

This yields a 4 × *L* attribution map per window (one channel per nucleotide) that accentuates positions where the model assigns unusually high probability to a specific base relative to the per-window mean predicted probability for that nucleotide. We transposed arrays to TF-MoDISco’s channel-first format and saved the one-hot inputs and attribution maps as .npz files (x_true.npz, x_pred.npz). We then ran TF-MoDISco-Lite (`modisco motifs -s x_true.npz -a x_pred.npz -n 1000000 -w 16384`) to sample one million seqlets across all 16,384 bp windows. TF-MoDISco-Lite then clustered those seqlets into consolidated motifs by first computing cosine similarity on gapped k-mer representations and next refining clusters via fine-grained realignment. For each resulting cluster, it computes both a contribution weight matrix (CWM) and a PWM. The `modisco report` step then generated motif logos and an interactive HTML summary.

#### Curating known TF-binding motifs from multiple yeast motif databases

We curated known TF-binding motifs from six publicly available databases, covering literature-curated associations, computational predictions, and high-throughput assay-derived specificity profiles. We included YEASTRACT, which provides 732 curated yeast motifs (average width 9.8 bp)^57^ (https://yeastract.com/); SwissRegulon Yeast, with 158 genome-wide motifs^56^ (https://swissregulon.unibas.ch/pages/); UniPROBE Yeast (GR09), containing 89 PBM-derived motifs^55^ (http://thebrain.bwh.harvard.edu/uniprobe/); MacIsaac v1, offering 124 phylogenetically conserved motifs^54^ (https://fraenkel-nsf.csbi.mit.edu/improved_map/); SCPD, supplying 24 promoter-derived motifs (https://esefinder.ahc.umn.edu/cgi-bin/tools/ESE3/esefinder.cgi)^58^; and YeTFaSCo, a comprehensive compendium of 1,709 motifs for 256 yeast proteins calculated and quality-assessed by multiple metrics (http://yetfasco.ccbr.utoronto.ca/)^53^. All motifs were downloaded in MEME format from the respective websites and merged to obtain a comprehensive yeast TF-binding motif database for integration into our downstream analyses.

#### Genome-wide enrichment analysis of TF-MoDISco motifs around TSS

To map TF-MoDISco-derived motifs to the genome coordinates and assess their proximity to transcription start sites (TSS), we first parsed the HDF5 output from the TF-MoDISco-Lite run (`modisco_results.h5`) to extract seqlet positions (start, end, example index, strand) for each pattern. We then converted these coordinates to genomic coordinates by adding the seqlet offsets to the corresponding entries in the original BED file of one-hot input windows, yielding a unified BED of motif hits annotated with species, chromosome, start, end, strand, and the best-matching known motif with its associated q-value.

Next, we generated a comprehensive TSS annotation by parsing GTF files. For each gene, we recorded the 5’ end of its transcripts (start coordinate for “+” strand, end for “–” strand) and sorted these TSS positions per chromosome. For each motif hit, we computed its midpoint and located the nearest TSS via binary search on the sorted list of transcripts, defining signed distance as negative for upstream (“–”) and positive for downstream (“+”) relative to the TSS strand.

To establish a background distribution, we sampled one random position per seqlet from the metaclusters on the same chromosome, drawing positions uniformly across chromosome lengths, and calculated each position’s closest-TSS distance using the same procedure. Finally, we compared the observed and background distance distributions by plotting histograms over a ±2.5 kb window centered on the TSS to highlight motif enrichment patterns.

### Embedding t-SNE clustering using Shorkie LM

To generate low-dimensional embeddings of genomic contexts, we first defined three genomic feature classes across all sixteen *S. cerevisiae* chromosomes: (1) Promoters: 500 bp immediately upstream of each start codon; (2) gene bodies: the span between each gene’s start and end coordinates; and (3) intergenic regions: all regions not annotated as gene, exon, or CDS. Using the Ensembl GTF biotypes, we further retained gene-body intervals corresponding to protein-coding genes, tRNAs, and transposable elements. Together with promoters and intergenic regions, these yielded 16,378 elements across five feature classes: “Protein-coding gene,” “tRNA,” “Transposable element,” “Promoter” and “Intergenic region”.

We extracted 500 bp of native genomic sequence centered on each element, reverse-complementing minus-strand elements, and centered this sequence within the LM input. We took the output of the first transformer block and averaged activations with weights proportional to real-sequence occupancy, excluding padding. We applied the same extraction and pooling to an untrained model with identical architecture. We used t-SNE ^121^ (scikit-learn `TSNE` with n_components = 2) to project the embeddings into two dimensions. The resulting 2D coordinates were visualized to assess clustering patterns by genomic features.

### Nucleotide dependency maps of Shorkie LM

We computed nucleotide dependency maps^63^ of Shorkie LM at six S. cerevisiae loci: the GAL1–GAL10 UAS (668 bp); the PHO5, CLN2 and HSP12 promoters (600 bp upstream to 100 bp downstream of the start codon); the RPL13B–RPS16A UAS (720 bp); and the DTD1 intron (196 bp), each centred in its native 16,384-bp context in genomic forward orientation. Before examining any model output, we defined motif occurrences as exact matches on either strand to the consensus sequences of the Gal4 UAS (CGG-N11-CCG), Pho4 E-box (CACGTK), SCB (CRCGAAA), Rap1 site (RMACCCANNCAYY), STRE (AGGGG) and branch point (TACTAAC), plus the annotated DTD1 5′ splice site (first six intron bases) and 3′ splice site (last three intron bases).

For each query position, we substituted each alternative nucleotide in the four nucleotide input channels and recorded, at every target position, the largest absolute change in predicted log2-odds over the three substitutions and four target nucleotides. Reference and mutant inputs were unmasked; probabilities were clipped to [10⁻⁷, 1 − 10⁻⁷], and self-dependencies were set to zero and excluded from inference. Maps were computed for the Shorkie LM on native sequence, an architecture-matched untrained model with Glorot-initialized weights, and Shorkie LM on a dinucleotide shuffle of the locus with native flanks.

### Shorkie and Shorkie_Random_Init data pre-processing

#### RNA-seq perturbation experiments

The design of the inducible genetic perturbation experiments builds on methods developed for the Induction Dynamics gene Expression Atlas^34^, where hundreds of transcription factors were independently induced and resultant gene expression changes were profiled over time using RNA hybridization microarrays. New data used for Shorkie-supervised training were generated at Calico Life Sciences LLC using updated miniaturized chemostats, or ministats. The instrumentation and RNA-sequencing protocols are described here.

#### Strain construction and selection

Each time-course experiment uses a strain selected from the Yeast Estradiol strains with Titratable Induction (YETI) collection^122^, in which the native promoter for a gene of interest was replaced by the synthetic β-estradiol-inducible Z3EV promoter. The released data comprise 3,053 induction RNA-seq tracks from 337 induced genes in two series (Figure 3-figure supplement 2). (1) Eight transcription factors (ARG80, CUP2, GAT4, MET4, MSN2, MSN4, NDT80 and RPN4), matched to previously measured microarray data, were induced in replicate and sampled at 0, 5, 10, 15, 30, 45, 60 and 90 min (580 tracks; 4–12 tracks per factor and time point). (2) A broader panel of 329 genes, including kinases, phosphatases and other transcriptional regulators prioritized as likely to have downstream transcriptional changes using YeastMine annotations^123^, Phenome^124^, and Fitness Clusters^125^, was sampled at 0, 5, 10, 20, 40, 70, 120 and 180 min (2,473 tracks; mostly one per gene and time point). In total, 384 of the 3,053 tracks are pre-induction (T0) samples.

#### Growth conditions

For all experiments, cells were grown under continuous culturing conditions. Cultures were maintained under phosphate limitation in minimal chemically defined media prepared by mixing 20 mL of 1,000x vitamin solution, 20 mL of 1,000x metals, 40 mL of 10 g/L KH_2_PO_4_, 1 L of 40% dextrose, and 2 L of 10x salts solution in 16 L of Milli-Q water to bring the total volume to 20 L. (Stock solutions are defined in Table S1).

#### Instrumentation and perturbation experiments

To increase throughput and coverage of gene expression changes, we developed a ministat array system capable of growing 24 cultures of 30 mL each in parallel to steady state, applying a chemical perturbation, and generating samples for time-resolved omic measurements. The system is housed in a 30°C warm room and consists of four banks of six 100 mL round-bottom vessels, integrated via a 24-vessel manifold into a complete ministat array (Figure 3-figure supplement 3).

Each ministat array incorporates three commercial peristaltic pumps: (1) a media pump that delivers fresh media to each vessel at a constant rate (the dilution rate), (2) a sampling pump that extracts a bolus of cells from each culture, and (3) an input pump used initially to inoculate cultures and subsequently to deliver a chemical perturbation (e.g., β-estradiol). Each pump uses separate tubing for each of the 24 ministats. Each vessel is also equipped with a tube for the effluent, which is continuously weighed to monitor and control the dilution rate, and a tube for delivering air. Air is supplied via a Flow Master 2400 airflow regulator connected to a humidifier to minimize culture evaporation. The regulator also splits the airflow into 24 individually adjustable lines, one for each culture.

For each culture, a pre-induction (*T*_0_) sample was collected prior to perturbation. The input pump then delivered 100 µL of 500 µM *β*-estradiol in base media from a 96-well plate into each vessel to initiate the time course. Following induction, cultures were typically sampled at 8 time points using the sampling pump to collect each sample into 96 deep-well plates pre-filled with chilled lysis buffer containing RNase inhibitor (Takara). Samples from the 24 ministats were staggered across quadrants of each 96-well plate, enabling systematic re-arraying of the eight time point-specific plates into two consolidated 96-well plates for downstream processing. Sample plates were flash frozen in liquid nitrogen immediately after collection.

#### RNA-seq library preparation

Samples from four time points were combined into a single 96-well PCR plate using the Bravo BenchCel system and processed using a miniaturized, high-throughput adaptation of standard library preparation protocols. After addition of oligo-dT primer (sequence: AAGCAGTGGTATCAACGCAGAG-TACTTTTTTTTTTTTTTTTTTTTTTTTTTTTTTVN), samples were lysed by three freeze-thaw cycles and incubated at 42°C for 3 minutes. cDNA synthesis was performed using the SMARTScribe Reverse Transcriptase kit (Takara), with 2.4 µM LNA template switch oligo (sequence: AAGCAGTGGTATCAAC-GCAGAGTACrGrG+G) and RNase inhibitor. For cDNA amplification, SeqAmp DNA polymerase (Takara) and ISO PCR primer (sequence: AAGCAGTGGTATCAACGCAGAGT) were added. The samples were placed in a thermocycler with the following conditions: 95°C for 1 min, 10 cycles of [98°C for 10 s, 65°C for 30 s, 68°C for 3 min], 72°C for 10 min. Cleanup was performed using RNAclean XP beads (Beckman Coulter) at 0.9x reaction volume.

Relative cDNA concentration was measured using Quant-iT PicoGreen (Fisher). For absolute quantification and quality control, 11 representative samples (chosen to span the range of PicoGreen values) were assessed using a High Sensitivity DNA kit (Agilent) on the BioAnalyzer. These values were used to generate a standard curve, and all samples were diluted to 200 pg/µL using a Mantis dispenser.

Library preparation was performed using the Nextera XT kit on cDNA from two ministat batches at a time, re-arrayed into a 384-well PCR plate. Samples were pooled, and pooled libraries underwent a double RNAclean XP bead cleanup. Final library concentration and quality were confirmed using a High Sensitivity DNA kit. Libraries were sequenced by Genewiz on the Illumina NovaSeq.

#### *S. cerevisiae* R64 reference genome pre-processing

Contigs from the *S. cerevisiae* R64 reference assembly were first split at assembly gaps and at hypervariable regions identified in the Rossi et al. study (these sites include the rDNA locus, tRNA genes and telomere regions and are available in 02_References_and_Features_Files at https://github.com/CEGRcode/2021-Rossi_Nature) hereafter referred to as rossi_mask.bed.

We then trimmed 1,024 bp from each contig end and discarded those shorter than 16,384 bp. Contigs longer than 786,432 bp were split in half to prevent large linkage blocks from biasing cross-validation partitions; this threshold represents 48 × the target window size, preserving sufficient sequence context while allowing granular randomization. The genome was then segmented into overlapping 16,384 bp windows with a 6,165 bp stride. These windows were shuffled and partitioned into eight cross-validation folds by balancing total nucleotide counts. Unmappable positions (from hypervariable regions described in rossi_mask.bed) were annotated, and any window with > 50% unmappability (`--umap_clip` 0.5) was removed.

#### ChIP-exo and ChIP-MNase sample preprocessing

We obtained paired-end sequencing data from Rossi et al.^65^ (GSE147927), aligned reads with BWA-MEM (v0.7.17)^126^, and removed multi-mapping reads with SAMtools^127^. For ChIP-exo data the position of the 5’ end of Read 1 was used, while the full span for MNase was used to generate tracks. We performed additional quality control on BAM files using Picard (https://github.com/broadinstitute/picard) to mark and remove duplicates. Reads overlapping regions described in rossi_mask.bed were removed and experiments with less than 10,000 remaining reads and greater than 75 percent duplication rate were dropped from the dataset. Lastly, BEDtools^128^ and bedGraphToBigWig (https://www.encodeproject.org/software/bedgraphtobigwig/) were used to transform the BAM files to BigWig tracks, yielding 1,128 ChIP-exo and 20 ChIP-MNase tracks.

#### RNA-seq sample preprocessing

We trimmed adapter sequences from FASTQ files using BBMap (https://github.com/BioInfoTools/BBMap), then aligned reads and quantified transcript abundance using STAR^129^. A genomic index was created using S288c_R63-3 and quantification was performed using GeneCounts with the following parameters `(--out-FilterMultimapNmax 1, --bamRemoveDuplicatesType UniqueIdentical, -- alignIntronMin 10, --alignIntronMax 2500, --alignMatesGapMax 2500`). BAM files were processed to remove PCR duplicates marked using PICARD^130^. Lastly, bam_cov.py (https://github.com/calico/basenji/blob/master/bin/bam_cov.py) was used to produce BigWig tracks. RNA-seq datasets from 1,000 strains (Caudal et al.) used unpaired reads^66^ and were filtered to retain samples with greater than 150,000 reads and less than 80 percent duplication rate while the in-house induction RNA-seq datasets used paired-end reads and were filtered to retain samples with greater than 150,000 reads and mean insert size greater than 250 bp. This produced 3,053 induction RNA-seq tracks and 1,014 1,000-strain RNA-seq tracks.

#### Validation

To ensure that gene expression dynamics measured using the new ministat array system are reliable and accurate, we compared data collected on the high-throughput ministat array to previously collected micro-array data from Hackett et al.^34^. To assess the correspondence between transcriptional profiles in each system, we calculated the Pearson correlation of gene expression fold-changes across all genes for a given matched time point post induction (Figure 3-figure supplement 4A), using both raw and log2 shrunken fold-changes. Next, to evaluate the new system’s sensitivity in detecting previously identified differentially expressed genes, we conducted an ROC analysis. To select a positive set of differentially expressed genes, we selected genes with an absolute log_2_ fold-change greater than one in the microarray data as a heuristic (Figure 3-figure supplement 4B). We then tested for differential expression in the new system by separately fitting a standard ordinary least squares linear regression for each gene’s expression time course following perturbation. In this model, gene expression relative to the mean *T*_0_ expression is described as a function of the time point, coded as a categorical variable for each time point, and a covariate for the culture vessel corresponding to an experimental replicate. Then we fit an ANOVA for each gene expression time course regression and used the F-statistic, which compares variance between time points with variance among replicates within a time point. By varying the cutoff F-statistic, we calculated the AUROC (Figure 3-figure supplement 4C-D).

#### Track data transformation

We processed 5,215 BigWig tracks: 3,053 induction time point RNA-seq datasets^34^, 1,014 RNA-seq datasets from various yeast strains^66^, 1,128 ChIP-exo tracks, and 20 ChIP-MNase tracks^65^. For each 16,384-bp window, we extracted per-base coverage and imputed missing values with the window’s median. To reduce edge effects, we cropped 1,024 bp from each end, retaining a 14,336-bp interior. We then summed coverage across consecutive 16-bp intervals to form 896 non-overlapping bins, yielding an 896-element vector per window. We saved these vectors as float16 in HDF5. Finally, we serialized the one-hot DNA sequence (via pysam), the corresponding binned coverage vector, and the unmappability mask into ZLIB-compressed TFRecord files (256 windows per file) organized by cross-validation fold.

### Shorkie model architecture and hyperparameters

Shorkie fine-tunes the pretrained Shorkie LM backbone (13.7 M parameters) to predict RNA-seq, ChIP-exo, and ChIP-MNase signals across 16,384 bp windows. Its trunk replicates Shorkie LM exactly:

• Initial Conv1D projection: 11 bp kernel × 96 filters (linear activation)

• Seven residual down-sampling blocks: each block is BatchNorm→GELU→Conv1D(5 bp) with filter counts of 96, 128, 160, 192, 256, 320, and 384, followed by 5% dropout, a skip connection, and Max-Pool1D (pool size = 2)

• Transformer bottleneck: eight layers operating over 128 positions. Each layer contains LayerNorm→4-head self-attention (model dimension = 384, key dimension = 64) with 20% dropout, followed by a feed-forward network (LayerNorm→Dense→ReLU→Dropout→Dense→Dropout) and residual connections.

The decoder uses the same U-Net up-sampling scheme as Shorkie LM but with only three up-sampling stages to restore the 16 bp resolution. At each stage, feature maps are batch-normalized, passed through a GELU nonlinearity, projected via a Dense layer (384 → 384), and doubled in length with UpSampling1D, then merged with the corresponding encoder output via a U-Net skip connection. At the final 16 bp resolution, a Cropping1D layer (cropping = 64) removes convolutional padding artifacts and is followed by GELU. A single Dense layer then projects each position into 5,215 channels: TF-perturbed RNA-seq (n = 3,053), 1,000-strain RNA-seq (n = 1,014), ChIP-exo (n = 1,128), and ChIP-MNase histone marks (n = 20), and a Softplus activation ensures all outputs remain positive.

We fine-tuned Shorkie with Adam (β_1_= 0.7, β_2_= 0.9; global clip-norm = 0.1), learning rate = 2 × 10^−5^ with 20,000 warm-up steps, batch = 8, using a Poisson + Multinomial loss (5x scaling of multinomial loss component) and early stopping with a patience of 150 epochs.

### Shorkie_Random_Init model architecture and hyperparameters

The Shorkie_Random_Init model uses the same architecture as Shorkie (including the same input/output setup and heads), but is trained with random weight initialization (no LM pretraining). We trained Shorkie_Random_Init in a supervised manner using Adam (β₁ = 0.9, β₂ = 0.999; global clip-norm = 0.1), learning rate = 5 × 10⁻^4^ with 5,000 warm-up steps, and otherwise identical hyperparameters. Shorkie_Random_Init thus serves as a baseline for quantifying the performance improvements attributable to transfer learning.

Because optimal optimization settings can differ between fine-tuning and training from random initialization, we additionally performed a learning-rate sweep for Shorkie_Random_Init over 5 × 10⁻⁶–5 × 10⁻³ under the same architecture, data splits, and training budget, and report the full learning curves and best-epoch summaries in Figure 3-figure supplement 8. Shorkie_Random_Init results in the main text use the best-performing learning rate from this sweep (LR = 5 × 10⁻³), providing a tuned from-scratch baseline for quantifying gains attributable to LM pretraining.

To rule out the possibility that the randomly initialized baseline was disadvantaged by over-parameterization and subsequent overfitting, we evaluated two reduced-capacity architectures trained from scratch: ConvNet_Smaller (3.40M parameters) and U-Net_Smaller (2.99M parameters). Reducing model capacity did not close the performance gap; both smaller architectures yielded validation Pearson’s *R* comparable to, or lower than, the full-size Shorkie_Random_Init (*R* = 0.523 ± 0.032), and all remained substantially below the pretrained Shorkie model (*R* = 0.633 ± 0.051) across validation metrics including *R* ^2^ (Figure 3-figure supplement 7).

### Shorkie and Shorkie_Random_Init bin-level, gene-level and track-level evaluation

#### Bin-level evaluation

We generated predictions at 16-bp resolution for each 16,384 bp input window by first averaging the model’s outputs across both the forward and reverse-complement strands (strand-ensemble) and then applying a shift-ensemble over offsets *S* = {0, 1} bp to smooth boundary artifacts. Specifically, the ensemble prediction at genomic position *i* and track *j* is defined as:

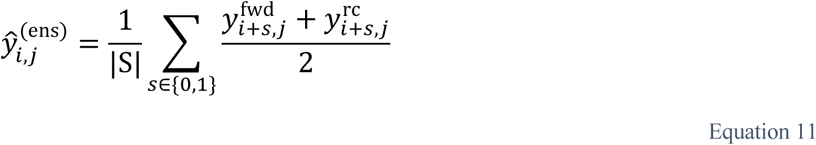

where 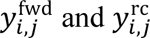 are the strand-specific model outputs. We compared 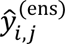 to measured coverage *y_i_*_,j_ and reported bin-level Pearson correlation and coefficient of determination (*R*^2^) across all tracks and cross-validation folds.

#### Raw (non-normalized) gene-level evaluation

Let *G* denote all annotated genes from the *S. cerevisiae* GTF. We assigned a given 16-bp output bin *i* to gene *g* if at least 50% of the bin’s width (*p*) overlaps an exon of gene *g*: ℬ(*g*) = {*b*: overlap(*b*, *g*) ≥ 0.5 *p*}.

For each gene *g* and track *j*, we aggregated the strand- and shift-ensemble predictions 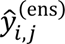 and the true coverages *y_i_*_,j_ over ℬ(*g*) and stabilized variance via a log_2_-transform with a pseudocount of 1:

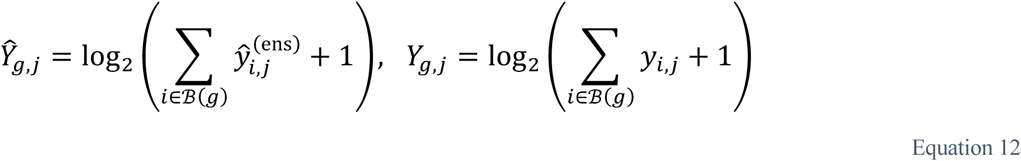

For each track *j*, we computed gene-level Pearson *r* and *R*^2^ between 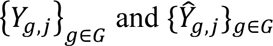.

#### Assay-normalized gene-level evaluation

To account for the differing dynamic ranges across assays, we first partitioned tracks into assay-specific groups *T_k_*, including TF-perturbed RNA-seq, 1,000-strain RNA-seq, ChIP-exo, and ChIP-MNase. Within each group *T_k_*, we independently applied quantile normalization (QN) to the log_2_-transformed aggregated coverages to equalize their distributions:

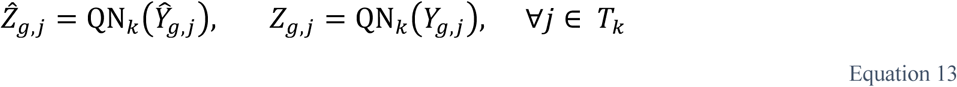

We then mean-centered each gene within its assay group:

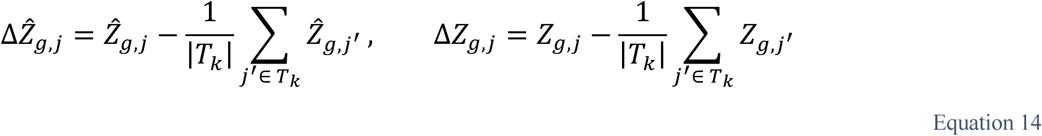

We summarized the gene-level concordance with Pearson *r* and *R*^2^ between {Δ*Z*_g,j_}_g∈*G*_ and {Δ*Ẑ*_g,j_}_g∈*G*_ for each *j*. Normalizing by assay-specific variability yields metrics that isolate predictive skill from absolute signal scale and enable evaluation of track specificity—that is, whether the model reproduces the differences among tracks within each assay.

#### Within-gene bin-level consistency

To evaluate how precisely Shorkie reconstructs fine-scale positional coverage within individual genes, we analyzed consistency at the bin-level within each gene. First, for each gene *g* and track *j*, we derived log_2_-transformed coverage vectors from the predictions and measured coverages:

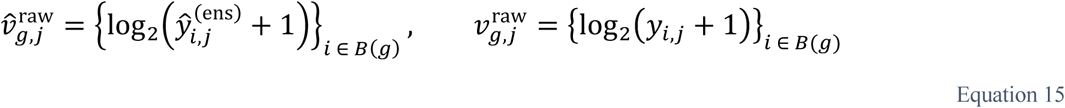

We computed Pearson *r* for gene-track pairs with non-trivial variation (both variances > 10^−6^). To de-emphasize nearly flat profiles, we further required 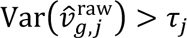, where *τ*_j_ is the 80th percentile of predicted variances across genes for track *j*. For each gene we summarized:

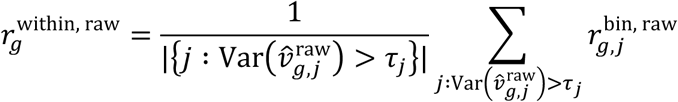

An identical procedure was conducted using quantile-normalized and mean-centered vectors to compute 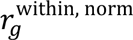.

#### Track-level evaluation

To evaluate Shorkie’s ability to accurately capture each gene’s pattern across RNA-seq assays specifically, we performed a track-level analysis focusing exclusively on RNA-seq tracks. For each gene *g*, we constructed vectors of coverage across all RNA-seq tracks, both in their raw and normalized (mean-centered and quantile-normalized) forms:

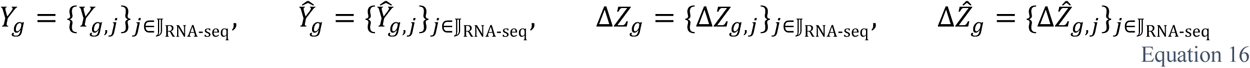

We reported Pearson *r* and *R*^2^ between predicted and observed vectors for both raw (*Y*_g_ and *Ŷ*_g_) and normalized (Δ*Z*_g_ and Δ*Ẑ*_g_) settings, summarizing per gene.

### Attention weight matrix visualization from selected Shorkie embeddings

To inspect how Shorkie’s transformer blocks focus on different parts of a gene, we extracted 16,384 bp windows centered on two example genes (EFM5 at chr VII: 489,391–505,775; RPL7A at chr VII: 356,973– 373,357) from the *S. cerevisiae* R64 reference using pysam. We center-trimmed or padded each window with “N”s to exactly 16,384 bp and parsed the Ensembl GTF with PyRanges^131^ to build an annotation table of gene bodies, exons, and 5’/3’ UTRs for downstream overlay.

We evaluated three models: Shorkie LM (pretrained LM), Shorkie (fine-tuned), and Shorkie_Random_Init (baseline). For each 16,384 × 4 input tensor, we averaged forward and reverse-complement predictions and concatenated results across replicates to yield a 1 × N_reps × 16,384 coverage array. To capture self-attention, we computed dot-products between learned query and key vectors at every position pair (*i*, *j*), applied softmax to obtain attention weights, and collected these across all eight cross-validation folds for (i) the first transformer block and (ii) the final two blocks. This yielded a single 128×128 attention map per block set.

For visualization, we clipped each fold-averaged attention map below 10⁻⁴ and above 0.05 to enhance contrast, then displayed it as a heatmap across the full window. We plotted predicted coverage as filled curves along the top and right margins. We converted genomic features into “attention-bin” coordinates (bin index = ⌊(position − window_start)/128⌋) and overlaid them as colored boxes for gene bodies or lines for UTRs and exons, with strand-specific coloring.

### *In silico* mutagenesis analysis of Shorkie and Shorkie_Random_Init

#### Sequence extraction and formatting

We applied *in silico* mutagenesis (ISM) to quantify the effect of every single-nucleotide variant (SNV) within yeast promoter regions based on Shorkie and Shorkie_Random_Init predictions. We defined input regions as 16,384-bp windows centered on promoter segments (450 bp upstream and 50 bp downstream of the TSS). The promoter set consisted of 137 ribosomal protein genes, 64 ribosome/rRNA biosynthesis (RRB) genes, and 3,258 additional protein-coding genes from the *S. cerevisiae* R64 reference genome. We extracted the corresponding sequences from the reference FASTA while preserving strand orientation.

For each 16 kb window, we computed ISM maps across the 500 bp promoter segment. For each reference sequence *s* and position *p* = 1,…, 500, we generated three mutant sequences by substituting the reference base with each of the three alternative nucleotides *n* ∈ {A, C, G, T} ∖ {ref}, yielding 3 × 500 mutant sequences per window.

#### ISM importance score matrix construction

We one-hot encoded all reference and mutant sequences and ran them through Shorkie and Shorkie_Random_Init with strand-ensemble averaging. For each sequence *s*, genomic bin *i* ∈ {1,…, 896}, and track *j*, let *y*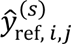 be the model prediction for the reference sequence, and 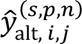 be the prediction for the variant at position *p* with nucleotide *n*. We computed the log_2_ fold-change score to quantify the variant effect:

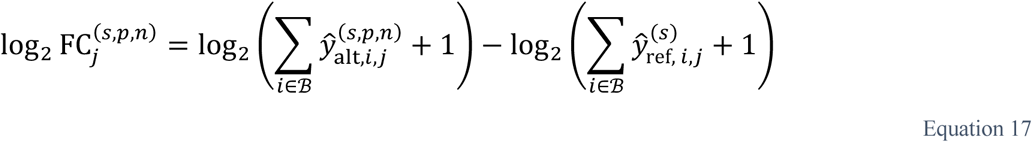

where ℬ is the set of output bins used for scoring. For Figure 4 and the motif analyses, ℬ contains all 896 output bins, so the score reflects the predicted coverage change across the whole 14,336 bp output window. For the time-course analyses in Figure 5, ℬ contains only the bins overlapping the annotated exons of the target gene, so the score measures the predicted effect on that gene. We saved the resulting log_2_ fold-change scores, together with metadata (chromosome, start, end, strand, reference, and alternate alleles), to an HDF5 file (scores.h5) via h5py (https://www.h5py.org/).

#### ISM importance score matrix normalization and ISM map visualization

To derive per-position importance profiles, we defined the set *J* of *T*_0_ RNA-Seq tracks. For each sequence *s*, mutation (*p*, *n*), and position *i*, we averaged the log_2_ fold-change scores across *T*_0_ tracks:

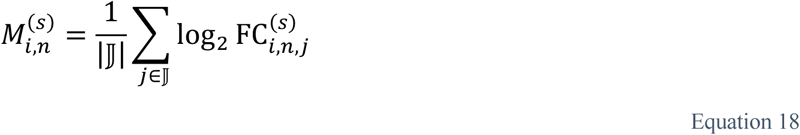

where *i* indexes positions and *n* denotes nucleotides. Next, we zero-mean normalized at each position *i* across the four nucleotides:

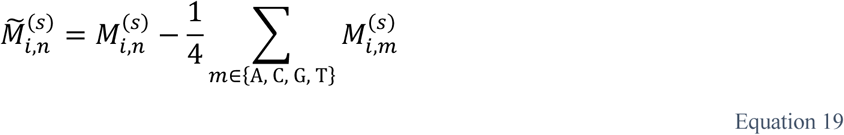

Finally, to focus on reference-base contributions, we computed:

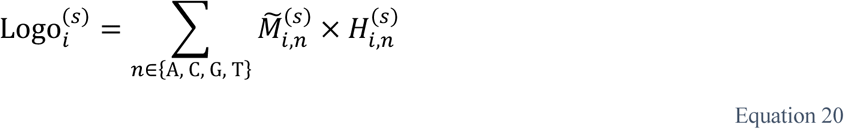

where 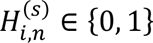 ∈ {0, 1} is the one-hot indicator for the reference base at position *i*. Finally, we visualized perposition scores 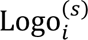 as DNA sequence logos (ISM maps), highlighting the magnitude and direction of variant effects across the promoter window.

### Quantitative comparison of known-motif recovery in ISM maps

#### Selecting promoter sequences for comparison

We analyzed 500-bp promoter windows (450 bp upstream to 50 bp downstream of the TSS, in gene orientation) from four cohorts: 64 ribosome and rRNA biosynthesis (RRB) genes and 387 ribosomal-protein and other protein-coding genes (RP/TSS) from the promoter set described above; 982 target genes of five transcription factors whose binding motifs are in the motif set below (Msn2, Msn4, Met4, Rpn4 and Swi4; regulons); and 256 protein-coding genes randomly selected without prior constraints on expression, motif content, or model predictions (independent panel). This independent panel was assembled via stratified sampling (16 genes from each of 16 bins defined by joint quartiles of promoter GC content and upstream intergenic length), strictly excluding genes belonging to the other cohorts.

#### Motif scanning and threshold calibration

We scanned both strands with 144 motifs: 139 high-confidence YeTFaSCo PWMs^53^ and five consensus motifs: PAC (GCGATGAG), RRPE (TGAAAAATTTT), TATA box (TATAAA), branch point (TACT-AAC) and 5′ splice site (GTATGT). We computed log-odds scores using a pseudocount of 10⁻³ and background composition estimated from 1,200 unselected promoter windows in the independent-panel sampling pool. We dinucleotide-shuffled each null window three times^132^ and calibrated motif-specific cutoffs at empirical tail probability ≤10⁻⁴. The motif-level false discovery rate (FDR) was the null exceedance rate multiplied by the number of scanned positions, divided by observed calls and capped at one.

#### Motif recovery scoring and normalization

We computed ISM maps for both Shorkie and Shorkie_Random_Init (Equations 17–20). We quantified motif recovery as the mean normalized absolute attribution score within the motif minus the mean score across its ±50 bp flanking regions, excluding other motif calls and ambiguous bases from the flanks. Finally, we excluded any motif calls with fewer than 20 eligible flanking nucleotides from downstream analyses.

#### Locus-level aggregation and statistical testing

We defined a locus as a single promoter window or a connected group of overlapping promoter windows on the same chromosome, irrespective of strand. After merging duplicate genomic intervals, the 1,685 pooled windows formed 1,440 loci.

We quantified between-model effect sizes as the median of the paired locus-level differences (Shorkie − Shorkie_Random_Init). We tested motifs with eligible occurrences in ≥15 loci (127 pooled motifs) using two-sided Wilcoxon signed-rank tests. Recovery above flanks was tested against zero using one-sided Wilcoxon signed-rank tests separately for each model and cohort. All Wilcoxon tests used asymptotic p-values. To control the family-wise error rate, we applied the Holm correction^133^ across the complete set of 144 motifs. We estimated pointwise 95% confidence intervals for the median paired difference using 10,000 paired-locus bootstrap resamples. Finally, to construct the empirical null band in Figure 4-figure supplement 3A, we evaluated 20 replicate sets of length-matched decoy sites placed at call-free, unambiguous positions within the same windows and plotted the central 95% distribution of their median paired differences.

#### Motif discovery using TF-MoDISco-Lite

To identify TF-binding motifs, we first constructed two arrays: ref.npz of shape (*N*_seq_, 4, *M*), containing one-hot encoded reference sequences, and pred.npz of shape (*N*_seq_, 4, *M*, |J|), containing per-replicate variant scores for *T*_0_ RNA-seq tracks.

Here, *N*_seq_ is the number of promoter windows, *M* = 500 is the promoter window length, and |J| is the number of *T*_0_ RNA-seq replicates. We reshaped pred.npz to (*N*_seq_ × |J|, 4, *M*) by concatenating across replicates, then ran TF-MoDISco-Lite (see “TF-MoDISco run on the *S. cerevisiae* genome” section) on these matrices to uncover recurring sequence motifs.

#### Time-series motif analysis of TF-induction with Shorkie

To capture dynamic motif changes during TF induction (e.g., MSN2, MSN4, MET4), we filtered the RNA-seq metadata to select only the tracks corresponding to each TF time course and grouped biological replicates by sampling time. We parsed time point annotations (e.g., “T_0”, “T_15”, “T_30”) directly from the track identifiers, ordered them chronologically, and for each time point *T_t_* defined J*_t_* as the set of associated RNA-seq track indices.

For each promoter window *s*, base position *i* ∈ {1, 2,…, 500}, nucleotide channel *n* ∈ {A, C, G, T}, and time point *T_t_*, we computed an ISM importance score matrix by averaging log_2_ fold-change scores across all replicate tracks:

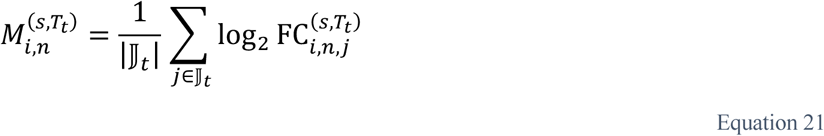

To focus on differential saliency relative to baseline expression, we baseline-corrected each time point by subtracting the *T*_0_ map:

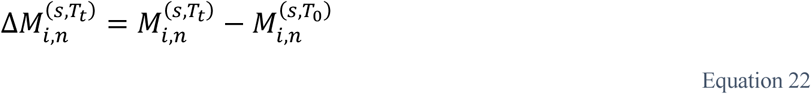

We then applied zero-mean normalization across nucleotides at each position to Δ*M* and visualized the resulting ΔISM maps for each *T_t_*. Finally, we reshaped per-time point ΔISM matrices to (*N*_seq_ × |J*_t_*|, 4, *M*) and applied TF-MoDISco-Lite to identify motifs whose importance trajectories changed over the induction time course.

#### Gene-level coverage calculation for experimental measurements and Shorkie predictions

To compare experimental and predicted gene-level coverages at Shorkie’s native 16 bp resolution, we summed bin values over each gene’s span. For gene *g* and track *j*, let ℬ(*g*) be the set of overlapping 16 bp bins. We define:

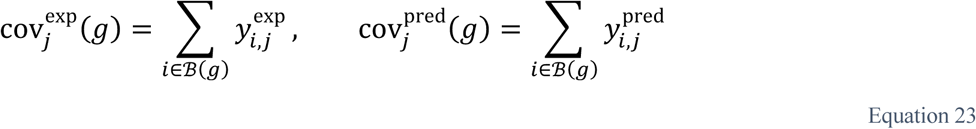

where ℬ(*g*) indexes the bins overlapping gene *g*. Next, we normalized to reads-per-million (RPM) using each sample’s library size libsize_j_,

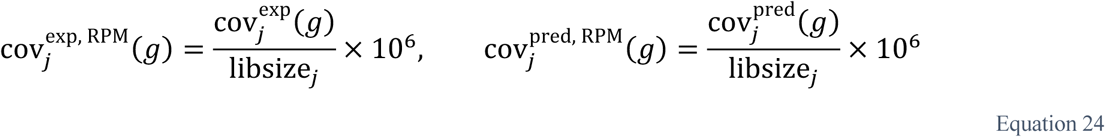

To characterize time-dependent gene expression trajectories in TF-induction experiments, we averaged RPM-normalized coverages across replicate tracks at each time point *T_t_*:

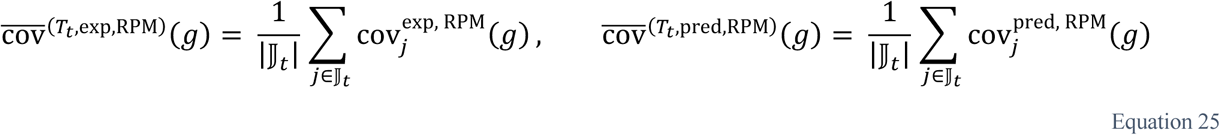

This yields matched, per-gene coverage trajectories for both experimental measurements and Shorkie predictions.

#### Euclidean distance calculation of ISM maps

To quantify motif importance changes during induction, we extracted the normalized importance score matrix **P***_s_*_,*t*_ ∈ ℝ*^L^*^×4^ for each promoter window *s* and time point *t*. We flattened this matrix into a vector **v***_s_*_,*t*_ ∈ ℝ^4*L*^, and computed pairwise Euclidean distances between time points:

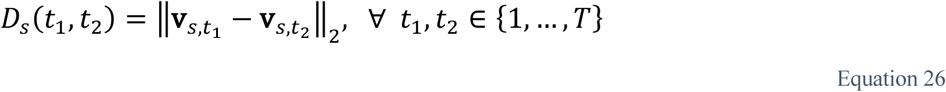

To summarize across all promoter windows, we conducted element-wise averaging:

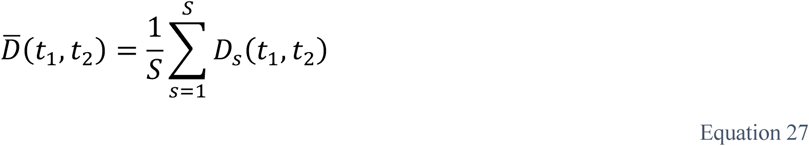

where *S* is the total number of promoter windows. We visualized the resulting mean Euclidean-distance matrix *D̄* as a heatmap, effectively capturing motif shifts over the TF induction time course.

### *cis*-eQTL analysis with Shorkie and DREAM challenge models

To benchmark *cis*-eQTL predictive performance, we compared Shorkie against the DREAM challenge models^94^ using two independent eQTL datasets. Because these datasets are association-based and not fine-mapped to a single causal allele, the reported effect sign can reflect the lead variant under linkage/allelic heterogeneity rather than the true causal direction. Accordingly, our primary evaluation targets regulatory impact magnitude, using absolute model-predicted variant effects to test for enrichment among putative eQTL lead SNPs relative to matched negative SNPs, rather than evaluating signed directionality.

#### Caudal et al. eQTLs from the pan-transcriptome of ∼1,000 yeast natural isolates

We imported the GWAS summary statistics^66^ (file: GWAS_combined_lgcCorr_ldPruned_noBonferroni_20221207.tab; downloaded from The 1002 Yeast Genome website: http://1002genomes.u-strasbg.fr/files/RNAseq) into a DataFrame and removed variants labeled as masked (ld_mask = “masked”). We normalized Phenotype (Pheno_pos) and SNP (ChrPos) positions by chromosome length, then classified a variant as *cis* if |ChrPos − Pheno_pos| ≤ 8,000 bp on the same chromosome and as *trans* otherwise. We further stratified variants by subtype (SNP vs. CNV) (Figure 7-figure supplement 1A).

We parsed the gVCF containing 1,011 yeast isolates (1011Matrix.gvcf; downloaded from http://1002genomes.u-strasbg.fr/files/)^96^ with pysam to extract reference and alternate alleles, chromosome, position, and quality. We then merged the *cis* and *trans* eQTL tables with the gVCF DataFrame on chromosome and position to identify intersecting and unique variant sets.

We removed variants with missing or non-positive P-values and computed significance as −log_10_(PValue). Finally, we generated a Manhattan plot by plotting −log_10_(PValue) against cumulative genomic position with alternating colors per chromosome and placed chromosome ticks at median cumulative positions (Figure 7-figure supplement 1B).

#### Kita et al. eQTLs from 85 diverse *S. cerevisiae* isolates

We imported the summary statistics (pnas.1717421114.sd01; downloaded from https://www.pnas.org/doi/suppl/10.1073/pnas.1717421114/suppl_file/pnas.1717421114.sd01.txt), yielding 1,640 eQTLs. From these, we selected 683 variants in four genomic contexts: Promoter, UTR5, UTR3, and ORF. For each variant, we computed the absolute distance between its genomic coordinate (ChrPos) and the target gene’s TSS, labeling it *cis* if |ChrPos − TSS| ≤ 8,000 bp and *trans* otherwise. Finally, we retrieved the corresponding allele sequences from 1011Matrix.gvcf using pysam.

#### Renganaath et al. causal cis-regulatory variants from an MPRA in a BY×RM yeast cross

We retrieved the variant-level MPRA summary statistics from Renganaath et al. (Supplementary File 3; https://cdn.elifesciences.org/articles/62669/elife-62669-supp3-v2.xlsx)^97^, which identified causal *cis*-regulatory variants in *S. cerevisiae* using a massively parallel reporter assay. The original study assayed 5,832 natural variants in the promoters of 2,503 genes between the laboratory strain BY and the vineyard isolate RM.

We restricted our evaluation exclusively to the “TSS” oligo library to ensure all variants fall within the core promoter region (≤144 bp upstream of the TSS). We purposefully excluded “distal” variants from the study’s “Upstream” library, as their synthetic MPRA context (which inserts an invariant HIS3 minimal promoter) diverges from the endogenous genomic sequence Shorkie relies on for its predictions. From the 6,361 tested variants evaluated via the mpra R package (5% FDR), we selected 142 true positive causal variants by requiring an adjusted P-value ≤ 0.05, a SNP-only classification (excluding indels), and a match between the significantly affected gene and the designed target gene (SignificantGeneIsDesignedGene column is True).

#### Negative-eQTL sampling

We generated four independent negative-eQTL sets by sampling common, non-coding variants that matched the positive eQTLs in allele composition and distance to the gene TSS. We extracted *S. cerevisiae* TSS coordinates from the Ensembl GTF. From the gVCF, we retained variants located outside CDS/exon intervals with allele frequency (AF) ≥ 0.05 and excluded any variant that matched a positive eQTL.

For each positive eQTL, we shuffled its matching candidate list of negative variants on the same chromosome and attempted to select one negative variant whose distance to a randomly chosen gene’s TSS matched the positive’s distance within ±100 bp (with a fallback window of ±200 bp if no match emerged). We enforced that each negative was used only once per iteration, yielding one negative per positive. We repeated this sampling four times to produce four distinct negative-eQTL sets.

#### Variant effect prediction with Shorkie

We predicted how individual variants alter gene-level coverage profiles with Shorkie. For each variant, we (1) extracted a 16,384 bp window centered on the SNP; (2) verified that the reference allele matched the extracted sequence; (3) generated one-hot encodings for both the reference and alternate alleles; and (4) averaged predictions over forward and reverse-complement strands. Within the window, we summed the predicted coverage values over all bins overlapping the annotated gene *g*’s exons:

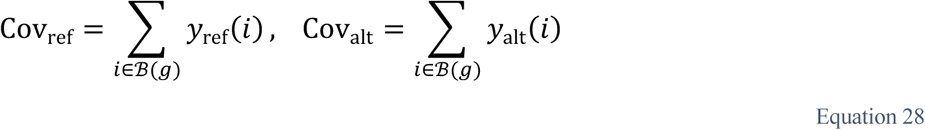

where ℬ(*g*) indexes the bins overlapping gene *g*. We then defined the log_2_ fold-change score as

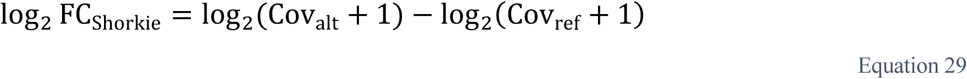

#### eQTL ISM analysis with Shorkie

To probe local sequence drivers, we applied the ISM pipeline to 80-bp windows centered on each SNP (to match DREAM input length), generated an 80 × 3 Δ matrix per variant, formed the reference-average matrix, and rendered sequence logos.

#### Predicting eQTL SNPs with DREAM challenge models

To benchmark Shorkie against established DREAM challenge models, we evaluated three pretrained models: convolutional (DREAM-CNN), recurrent (DREAM-RNN), and attention-based (DREAM-Atten), on both positive eQTLs and the four independently sampled negative sets.

#### Sequence extraction and formatting

For each variant (positive or negative), we extracted an 80-bp window centered on the SNP from the *S. cerevisiae* genome (40 bp upstream and 39 bp downstream), then prepended a fixed 17-bp upstream flank (TGCATTTTTTTCACATC) and appended a 13 bp downstream flank (GGTTACGGCTGTT), following the DREAM models’ expected input format, yielding final input sequences of 110 bp.

#### Model inference and log fold-change score calculation

We one-hot encoded the 110-bp reference and alternate sequences and scored them on a GPU to obtain scalar predictions, then converted them to log_2_ FC using the shared definition.

#### ISM analysis of DREAM-RNN

We performed single-nucleotide ISM on the 110-bp inputs by iterating over each position *p* = 1,…, 110 and substituting the native nucleotide with each of the three alternatives (*n* ∈ {A, C, G, T} ∖ {ref}); we then computed Δ(*p*, *n*), constructed the reference-average matrix, and visualized sequence logos.

#### Massively parallel reporter assay (MPRA) evaluation of promoter variants

We evaluated Shorkie’s predictive performance for regulatory variants measured by a publicly available MPRA experiment^94^. The MPRA included five single-sequence sub-libraries: high_exp, low_exp, yeast_exp, random_exp, challenging_exp, each containing unique 110 bp promoter sequences, and three dual-sequence sub-libraries: SNVs_exp, motif_perturbation_exp, motif_tiling_exp, consisting of paired reference and alternate sequences. We excluded constructs present in any public leaderboard and randomly drew a uniform sample of 1,000 sequences without replacement from each category.

#### Gene selection and library sampling

We stratified *T*_0_ RNA-seq expression into three quantiles (5–25%, 25–75%, 75–95%) separately by strand. From each quantile, we randomly chose three to four genes, resulting in 22 genes:

• Forward strand genes (10 total):

- 5–25%: GPM3/SLI1/VPS52
- 25–75%: YMR160W/MRPS28/YCT1
- 75–95%: RDL2/PHS1/RTC3/MSN4

- Reverse strand genes (12 total):
- 5–25%: COA4/ERI1/RSM25/AIM11
- 25–75%: ERD1/MRM2/SNT2/MRPL1
- 75–95%: CSI2/RPE1/PKC1/MAE1

#### Promoter insertion design

To examine how Shorkie’s predictions vary with distance from the TSS, we first defined a promoter insertion window of 110 bp (±55 bp) and enforced a minimum 100-bp offset from the TSS to avoid direct overlap. We then chose eleven distinct insertion offsets (ranging from 100 to 200 bp, incremented by 10 bp). For each selected gene, we calculated the insertion midpoints as follows:

• Forward strand: midpoint = TSS – offset

• Reverse strand: midpoint = TSS + offset

Each midpoint defined a 110 bp replacement window from (midpoint − 55 bp) to (midpoint + 55 bp). We then replaced these native genomic windows with each MPRA-derived sequence and used Shorkie to predict the regulatory impact. In contrast, DREAM challenge models directly utilized the 110 bp MPRA windows as input to predict scalar expression scores.

#### Promoter variant effect quantification

To quantify gene-level effects from Shorkie’s predictions, we identified model-predicted coverage bins *B*(*g*) overlapping the exonic regions of each target gene (*g*).

For single-sequence libraries, we computed Cov_native_ = ∑*_i_*_∈ℬ(g)_ *y*_native_(*i*) and Cov_MPRA_ = ∑*_i_*_∈ℬ(g)_ *y*_MPRA_(*i*). We reported the regulatory effect as log_2_ FC_single_ = log_2_(Cov_MPRA_ + 1) − log_2_(Cov_native_ + 1). log_2_ FC_single_ reflects changes in predicted coverage due to the MPRA insert relative to the native sequence.

In dual-sequence libraries, each construct includes both reference and alternate promoter variants. Shorkie predicted REF and ALT coverage separately, and we computed log_2_ FC_ref_ and log_2_ FC_alt_ relative to the native promoter and summarized the predicted differential effect as Δlog_2_ FC_dual_ = log_2_ FC_alt_ − log_2_ FC_ref_.

Experimental reporter assays yielded corresponding expression scores for the reference (*S*_ref_) and alternate (*S*_alt_) sequences. We quantified their difference as Δ*S* = *S*_alt_ − *S*_ref_. Finally, we visualized scatterplots to assess the concordance between the Shorkie-predicted Δlog_2_ FC_dual_ and the experimentally derived Δ*S* scores.

## Data and Code Availability

• The new Induction Dynamics Gene Expression Atlas RNA-seq datasets generated by Calico Life Sciences LLC are hosted on Google Cloud Storage (GCS). Source FASTQ files are available at gs://shorkie-paper-raw/data/supervised/raw/rnaseq/ and gs://shorkie-paper-rawdata/supervised/raw/novaseq-2021/. Corresponding adapter-trimmed paired-end FASTQs, sample sheets, quality-control files, and preprocessing outputs are under gs://shorkie-paper-raw/data/supervised/raw/ID2/.

• BigWig coverage tracks used for supervised training—including the new IDEA RNA-seq data, published yeast-strain RNA-seq data, and ChIP-exo/ChIP-MNase data—are available at gs://shorkie-paper/data/supervised/bigwigs/, with processed TFRecords at gs://shorkie-paper/data/supervised/processed.

• The genomes used for self-supervised language-model pretraining are at gs://shorkie-paper/data/unsupervised/genome/, with corresponding TFRecords at gs://shorkie-paper/data/unsupervised/processed/.

• The Shorkie LM and Shorkie models are implemented in TensorFlow. Shorkie LM is available at: gs://seqnn-share/shorkie_models/shorkie_lm/. Shorkie models are available at: gs://seqnnshare/shorkie_models/shorkie/. Shorkie_Random_Init models are available at: gs://seqnnshare/shorkie_models/shorkie_random_init/

• Parameters, training code and evaluation scripts for both the Shorkie LM and Shorkie models are available under the Apache-2.0 license at https://github.com/calico/baskerville-yeast and https://github.com/calico/shorkie-paper.

## Author Contributions

J.L. and D.R.K. conceived the project.

K-H.C., M.M.M., J.L., and D.R.K. designed the research.

K-H.C., J.L., and D.R.K. developed the baskerville-yeast repository.

S.H. led design of the yeast induction experiments and instrumentation.

E.S. optimized library preparation miniaturization, processed samples and generated sequencing libraries for the induction experiments.

M.M.M. processed the RNA-seq, ChIP-exo, and ChIP-MNase data for model training.

K-H.C. and J.L. trained the Shorkie LM models.

K-H.C. and M.M.M. trained the Shorkie_Random_Init models.

K-H.C. trained the Shorkie models.

K-H.C. and J.L. conducted analyses on Shorkie LM transcription factor motif inference and Shorkie model interpretability.

K-H.C. and J.L. conducted the MPRA and eQTL analyses.

K-H.C., M.M.M., E.S., S.H., J.L., and D.R.K. wrote the manuscript.

## Funding

This research was supported in part by the U.S. National Institutes of Health (NIH) under grants R01-HG006677 and R35-GM156470, and by the U.S. National Science Foundation (NSF) under grant DBI-2412449. Computational analyses were performed using resources provided by the Advanced Research Computing at Hopkins (ARCH) core facility, supported in part by NSF grant OAC-1920103. Additional funding was provided by Calico Life Sciences LLC.

## Acknowledgements

We gratefully acknowledge all members of the Kelley Lab at Calico Life Sciences LLC for their insightful discussions. We thank David Botstein for guidance and feedback on planning the induction time course experiments. We also are grateful for contributions from Griffin Kim for designing the ministat arrays, Rebecca Wang for adapting the library prep and automation of the higher throughput miniaturized vessels, and Thomas Li for assistance generating the experimental data. We also thank Steven Salzberg and Mihaela Pertea for their valuable ideas and thoughtful feedback. The Shorkie logo was generated with the help of OpenAI in the style of Borzoi (https://github.com/calico/borzoi/blob/main/borzoi_logo.png).

## Notes

https://github.com/calico/shorkie-paper

https://github.com/calico/baskerville-yeast

### Competing Interest Statement

The authors have declared no competing interest.

### Summary of Updates

Summary of major revisions * Corrected baseline metrics and confidence intervals: Against the tuned baseline, Shorkie improves median bin-level Pearson's R from 0.70 to 0.78 and gene-level Pearson's R from 0.80 to 0.88. All RNA-seq metrics of Figure 3 now report paired cross-fold uncertainty (Figure 3-figure supplement 9). * Narrowed the two claims flagged in the assessment. We no longer describe the Saccharomycetales corpus as an evolutionary optimum, and we no longer state that Shorkie LM learns regulatory grammar. We now describe the corpus as the best-performing initialization among those tested, and the interpretation analyses as evidence of motif recovery and, at selected loci, of inter-element dependencies. * MPRA with Shorkie_Random_Init. Shorkie outperforms Shorkie_Random_Init in all five DREAM MPRA categories, most strongly for native promoters and motif perturbation effects (Figure 6-figure supplement 3). * Length and padding control for Figure 2E. Embeddings were re-extracted from a fixed 500-bp native context with occupancy-weighted pooling and compared with an untrained network (Figure 2-figure supplement 3); Figure 2E is replaced. * Nucleotide dependency maps at six regulatory loci (Figure 2-figure supplement 4). * Quantitative ISM motif recovery of Shorkie versus Shorkie_Random_Init with paired, multiple-testing-corrected statistics (Figure 4-figure supplement 3), replacing the qualitative statement. * Target-gene ISM for Figure 5, which now sums predicted coverage only over the target gene (Figure 5; Response Figure 1). * Documentation and figures describing the induction RNA-seq resource (Figure 3-figure supplement 2), the loss weighting, the Figure 6B-C colours and the Figure 5 legend.

## References

1. Harbison, C. T. et al. Transcriptional regulatory code of a eukaryotic genome. Nature 431, 99–104 (2004).

2. He, Q., Johnston, J. & Zeitlinger, J. ChIP-nexus enables improved detection of in vivo transcription factor binding footprints. Nat Biotechnol 33, 395–401 (2015).

3. Lee, T. I. et al. Transcriptional Regulatory Networks in *Saccharomyces cerevisiae*. Science 298, 799– 804 (2002).

4. Rhee, H. S. & Pugh, B. F. Comprehensive Genome-wide Protein-DNA Interactions Detected at Single-Nucleotide Resolution. Cell 147, 1408–1419 (2011).

5. Skene, P. J. & Henikoff, S. An efficient targeted nuclease strategy for high-resolution mapping of DNA binding sites. eLife 6, e21856 (2017).

6. Struhl, K. Molecular mechanisms of transcriptional regulation in yeast. Annual review of biochemistry 58, 1051–1077 (1989).

7. Venters, B. J. et al. A Comprehensive Genomic Binding Map of Gene and Chromatin Regulatory Proteins in Saccharomyces. Molecular Cell 41, 480–492 (2011).

8. Venters, B. J. & Pugh, B. F. A canonical promoter organization of the transcription machinery and its regulators in the *Saccharomyces* genome. Genome Res. 19, 360–371 (2009).

9. Weiner, A. et al. High-Resolution Chromatin Dynamics during a Yeast Stress Response. Molecular Cell 58, 371–386 (2015).

10. Zentner, G. E., Kasinathan, S., Xin, B., Rohs, R. & Henikoff, S. ChEC-seq kinetics discriminates transcription factor binding sites by DNA sequence and shape in vivo. Nat Commun 6, 8733 (2015).

11. Zhu, J. et al. Integrating large-scale functional genomic data to dissect the complexity of yeast regulatory networks. Nat Genet 40, 854–861 (2008).

12. Beer, M. A. & Tavazoie, S. Predicting Gene Expression from Sequence. Cell 117, 185–198 (2004).

13. Sharon, E. et al. Inferring gene regulatory logic from high-throughput measurements of thousands of systematically designed promoters. Nat Biotechnol 30, 521–530 (2012).

14. De Boer, C. G. et al. Deciphering eukaryotic gene-regulatory logic with 100 million random promoters. Nat Biotechnol 38, 56–65 (2020).

15. Vaishnav, E. D. et al. The evolution, evolvability and engineering of gene regulatory DNA. Nature 603, 455–463 (2022).

16. Zhou, Z., et al. DNABERT-2: Efficient Foundation Model and Benchmark For Multi-Species Genome. Preprint at 10.48550/ARXIV.2306.15006 (2023).

17. Ji, Y., Zhou, Z., Liu, H. & Davuluri, R. V. DNABERT: pre-trained Bidirectional Encoder Representations from Transformers model for DNA-language in genome. Bioinformatics 37, 2112–2120 (2021).

18. Benegas, G., Batra, S. S. & Song, Y. S. DNA language models are powerful predictors of genome-wide variant effects. Proc. Natl. Acad. Sci. U.S.A. 120, e2311219120 (2023).

19. Zhai, J. et al. Cross-species modeling of plant genomes at single nucleotide resolution using a pretrained DNA language model. Preprint at 10.1101/2024.06.04.596709 (2024).

20. Sanabria, M., Hirsch, J., Joubert, P. M. & Poetsch, A. R. DNA language model GROVER learns sequence context in the human genome. Nat Mach Intell 6, 911–923 (2024).

21. Karollus, A. et al. Species-aware DNA language models capture regulatory elements and their evolution. Genome Biol 25, 83 (2024).

22. Dalla-Torre, H. et al. Nucleotide Transformer: building and evaluating robust foundation models for human genomics. Nat Methods 22, 287–297 (2025).

23. Dao, T. & Gu, A. Transformers are SSMs: Generalized Models and Efficient Algorithms Through Structured State Space Duality. Preprint at 10.48550/ARXIV.2405.21060 (2024).

24. Brixi, G. et al. Genome modeling and design across all domains of life with Evo 2. Preprint at 10.1101/2025.02.18.638918 (2025).

25. Schiff, Y., et al. Caduceus: Bi-Directional Equivariant Long-Range DNA Sequence Modeling. Preprint at 10.48550/ARXIV.2403.03234 (2024).

26. Nguyen, E., et al. HyenaDNA: Long-Range Genomic Sequence Modeling at Single Nucleotide Resolution. Preprint at 10.48550/ARXIV.2306.15794 (2023).

27. Nguyen, E. et al. Sequence modeling and design from molecular to genome scale with Evo. Science 386, eado9336 (2024).

28. Hayes, T. et al. Simulating 500 million years of evolution with a language model. Science 387, 850– 858 (2025).

29. Madani, A. et al. Large language models generate functional protein sequences across diverse families. Nat Biotechnol 41, 1099–1106 (2023).

30. Nijkamp, E., Ruffolo, J. A., Weinstein, E. N., Naik, N. & Madani, A. ProGen2: Exploring the boundaries of protein language models. Cell Systems 14, 968–978.e3 (2023).

31. Bhatnagar, A. et al. Scaling unlocks broader generation and deeper functional understanding of proteins. Preprint at 10.1101/2025.04.15.649055 (2025).

32. Lin, Z. et al. Evolutionary-scale prediction of atomic-level protein structure with a language model. Science 379, 1123–1130 (2023).

33. Shen, X.-X. et al. Tempo and Mode of Genome Evolution in the Budding Yeast Subphylum. Cell 175, 1533–1545.e20 (2018).

34. Hackett, S. R. et al. Learning causal networks using inducible transcription factors and transcriptome-wide time series. Molecular Systems Biology 16, e9174 (2020).

35. Avsec, Ž., et al. Effective gene expression prediction from sequence by integrating long-range interactions. Nat Methods 18, 1196–1203 (2021).

36. Linder, J., Srivastava, D., Yuan, H., Agarwal, V. & Kelley, D. R. Predicting RNA-seq coverage from DNA sequence as a unifying model of gene regulation. Nat Genet 10.1038/s41588-024-02053-6 (2025) doi:10.1038/s41588-024-02053-6.

37. Ronneberger, O., Fischer, P. & Brox, T. U-Net: Convolutional Networks for Biomedical Image Segmentation. in Medical Image Computing and Computer-Assisted Intervention – MICCAI 2015 (eds. Navab, N., Hornegger, J., Wells, W. M. & Frangi, A. F.) vol. 9351 234–241 (Springer International Publishing, Cham, 2015).

38. Huerta-Cepas, J., Serra, F. & Bork, P. ETE 3: Reconstruction, Analysis, and Visualization of Phylogenomic Data. Mol Biol Evol 33, 1635–1638 (2016).

39. Letunic, I. & Bork, P. Interactive Tree of Life (iTOL) v6: recent updates to the phylogenetic tree display and annotation tool. Nucleic Acids Research 52, W78–W82 (2024).

40. Letunic, I. & Bork, P. Interactive Tree Of Life (iTOL): an online tool for phylogenetic tree display and annotation. Bioinformatics 23, 127–128 (2007).

41. Marçais, G. et al. MUMmer4: A fast and versatile genome alignment system. PLoS Comput Biol 14, e1005944 (2018).

42. Ondov, B. D. et al. Mash: fast genome and metagenome distance estimation using MinHash. Genome Biol 17, 132 (2016).

43. Bailey, T. L. et al. MEME SUITE: tools for motif discovery and searching. Nucleic Acids Research 37, W202–W208 (2009).

44. Flynn, J. M., et al. RepeatModeler2 for automated genomic discovery of transposable element families. Proc. Natl. Acad. Sci. U.S.A. 117, 9451–9457 (2020).

45. Smit, A. F. A., Hubley, R. & Green, P. RepeatMasker Open-4.0. (2013).

46. Storer, J., Hubley, R., Rosen, J., Wheeler, T. J. & Smit, A. F. The Dfam community resource of transposable element families, sequence models, and genome annotations. Mobile DNA 12, 2 (2021).

47. Ou, S. & Jiang, N. LTR_retriever: A Highly Accurate and Sensitive Program for Identification of Long Terminal Repeat Retrotransposons. Plant Physiol. 176, 1410–1422 (2018).

48. Price, A. L., Jones, N. C. & Pevzner, P. A. De novo identification of repeat families in large genomes. Bioinformatics 21, i351–i358 (2005).

49. Li, H. Minimap2: pairwise alignment for nucleotide sequences. Bioinformatics 34, 3094–3100 (2018).

50. Rafi, A. M., Kiyota, B., Yachie, N. & De Boer, C. Detecting and avoiding homology-based data leakage in genome-trained sequence models. Preprint at 10.1101/2025.01.22.634321 (2025).

51. Sahu, B. et al. Sequence determinants of human gene regulatory elements. Nat Genet 54, 283–294 (2022).

52. Shrikumar, A., et al. Technical Note on Transcription Factor Motif Discovery from Importance Scores (TF-MoDISco) version 0.5.6.5. Preprint at 10.48550/arXiv.1811.00416 (2020).

53. De Boer, C. G. & Hughes, T. R. YeTFaSCo: a database of evaluated yeast transcription factor sequence specificities. Nucleic Acids Research 40, D169–D179 (2012).

54. MacIsaac, K. D. et al. An improved map of conserved regulatory sites for Saccharomyces cerevisiae. BMC Bioinformatics 7, (2006).

55. Newburger, D. E. & Bulyk, M. L. UniPROBE: an online database of protein binding microarray data on protein-DNA interactions. Nucleic Acids Research 37, D77–D82 (2009).

56. Pachkov, M., Erb, I., Molina, N. & Van Nimwegen, E. SwissRegulon: a database of genome-wide annotations of regulatory sites. Nucleic Acids Research 35, D127–D131 (2007).

57. Teixeira, M. C., et al. The YEASTRACT database: an upgraded information system for the analysis of gene and genomic transcription regulation in*Saccharomyces cerevisiae*. Nucl. Acids Res. 42, D161– D166 (2014).

58. Zhu, J. & Zhang, M. Q. SCPD: a promoter database of the yeast Saccharomyces cerevisiae. Bioinformatics 15, 607–611 (1999).

59. Yabana, N. & Yamamoto, M. *Schizosaccharomyces pombe map1*^+^ Encodes a MADS-Box-Family Protein Required for Cell-Type-Specific Gene Expression. Molecular and Cellular Biology 16, 3420– 3428 (1996).

60. Nielsen, O., Friis, T. & Kjærulff, S. The Schizosaccharomyces pombe map1 gene encodes an SRF / MCM1-related protein required for P-cell specific gene expression. Mol Gen Genet 253, 387–392 (1996).

61. Casselton, L. A. Mate recognition in fungi. Heredity 88, 142–147 (2002).

62. Oliva, A. et al. The Cell Cycle–Regulated Genes of Schizosaccharomyces pombe. PLoS Biol 3, e225 (2005).

63. Tomaz Da Silva, P., et al. Nucleotide dependency analysis of genomic language models detects functional elements. Nat Genet 57, 2589–2602 (2025).

64. Tomaz Da Silva, P. et al. Nucleotide dependency analysis of DNA language models reveals genomic functional elements. Preprint at 10.1101/2024.07.27.605418 (2024).

65. Rossi, M. J. et al. A high-resolution protein architecture of the budding yeast genome. Nature 592, 309–314 (2021).

66. Caudal, É. et al. Pan-transcriptome reveals a large accessory genome contribution to gene expression variation in yeast. Nat Genet 56, 1278–1287 (2024).

67. Routhier, E., Pierre, E., Khodabandelou, G. & Mozziconacci, J. Genome-wide prediction of DNA mutation effect on nucleosome positions for yeast synthetic genomics. Genome Res. 31, 317–326 (2021).

68. Martin, D. E., Soulard, A. & Hall, M. N. TOR Regulates Ribosomal Protein Gene Expression via PKA and the Forkhead Transcription Factor FHL1. Cell 119, 969–979 (2004).

69. Schawalder, S. B. et al. Growth-regulated recruitment of the essential yeast ribosomal protein gene activator Ifh1. Nature 432, 1058–1061 (2004).

70. Rudra, D., Zhao, Y. & Warner, J. R. Central role of Ifh1p–Fhl1p interaction in the synthesis of yeast ribosomal proteins. EMBO J 24, 533–542 (2005).

71. Reja, R., Vinayachandran, V., Ghosh, S. & Pugh, B. F. Molecular mechanisms of ribosomal protein gene coregulation. Genes Dev. 29, 1942–1954 (2015).

72. Shore, D. RAP1: a protean regulator in yeast. Trends in Genetics 10, 408–412 (1994).

73. Wade, J. T., Hall, D. B. & Struhl, K. The transcription factor Ifh1 is a key regulator of yeast ribosomal protein genes. Nature 432, 1054–1058 (2004).

74. Warner, J. R. The economics of ribosome biosynthesis in yeast. Trends in Biochemical Sciences 24, 437–440 (1999).

75. Jorgensen, P. et al. A dynamic transcriptional network communicates growth potential to ribosome synthesis and critical cell size. Genes Dev. 18, 2491–2505 (2004).

76. Wade, C. H., Umbarger, M. A. & McAlear, M. A. The budding yeast rRNA and ribosome biosynthesis (RRB) regulon contains over 200 genes. Yeast 23, 293–306 (2006).

77. Arnone, J. T. & McAlear, M. A. Adjacent Gene Pairing Plays a Role in the Coordinated Expression of Ribosome Biogenesis Genes *MPP10* and *YJR003C* in Saccharomyces cerevisiae. Eukaryot Cell 10, 43–53 (2011).

78. Hughes, J. D., Estep, P. W., Tavazoie, S. & Church, G. M. Computational identification of Cis-regulatory elements associated with groups of functionally related genes in Saccharomyces cerevisiae 1 1Edited by F. E. Cohen. Journal of Molecular Biology 296, 1205–1214 (2000).

79. Jorgensen, P., Nishikawa, J. L., Breitkreutz, B.-J. & Tyers, M. Systematic Identification of Pathways That Couple Cell Growth and Division in Yeast. Science 297, 395–400 (2002).

80. Brown, S. J., Cole, M. D. & Erives, A. J. Evolution of the holozoan ribosome biogenesis regulon. BMC Genomics 9, 442 (2008).

81. Schirman, D., Yakhini, Z., Pilpel, Y. & Dahan, O. A broad analysis of splicing regulation in yeast using a large library of synthetic introns. PLoS Genet 17, e1009805 (2021).

82. Moore, M. J., Query, C. C., Sharp, P. A., & others. Splicing of precursors to mRNAs by the spliceosome. Cold Spring Harbor Monograph Series 24, 303–303 (1993).

83. Parker, R., Siliciano, P. G. & Guthrie, C. Recognition of the TACTAAC box during mRNA splicing in yeast involves base pairing to the U2-like snRNA. Cell 49, 229–239 (1987).

84. Zavanelli, M. I. & Ares, M. Efficient association of U2 snRNPs with pre-mRNA requires an essential U2 RNA structural element. Genes & Development 5, 2521–2533 (1991).

85. Robinson, J. T., Thorvaldsdottir, H., Turner, D. & Mesirov, J. P. igv.js: an embeddable JavaScript implementation of the Integrative Genomics Viewer (IGV). Bioinformatics 39, btac830 (2023).

86. Engel, S. R., et al. *Saccharomyces* Genome Database: advances in genome annotation, expanded bio-chemical pathways, and other key enhancements. GENETICS 229, iyae185 (2025).

87. Garmendia-Torres, C., Goldbeter, A. & Jacquet, M. Nucleocytoplasmic Oscillations of the Yeast Transcription Factor Msn2: Evidence for Periodic PKA Activation. Current Biology 17, 1044–1049 (2007).

88. Ni, L. et al. Dynamic and complex transcription factor binding during an inducible response in yeast. Genes Dev. 23, 1351–1363 (2009).

89. Martínez-Pastor, M. T. et al. The Saccharomyces cerevisiae zinc finger proteins Msn2p and Msn4p are required for transcriptional induction through the stress response element (STRE). The EMBO Journal 15, 2227–2235 (1996).

90. Gasch, A. P. et al. Genomic Expression Programs in the Response of Yeast Cells to Environmental Changes. MBoC 11, 4241–4257 (2000).

91. Causton, H. C. et al. Remodeling of Yeast Genome Expression in Response to Environmental Changes. MBoC 12, 323–337 (2001).

92. Kuras, L., Barbey, R. & Thomas, D. Assembly of a bZIP-bHLH transcription activation complex: formation of the yeast Cbf1-Met4-Met28 complex is regulated through Met28 stimulation of Cbf1 DNA binding. The EMBO Journal 16, 2441–2451 (1997).

93. Lee, T. A. et al. Dissection of Combinatorial Control by the Met4 Transcriptional Complex. MBoC 21, 456–469 (2010).

94. Rafi, A. M. et al. A community effort to optimize sequence-based deep learning models of gene regulation. Nat Biotechnol 10.1038/s41587-024-02414-w (2024) doi:10.1038/s41587-024-02414-w.

95. Kita, R., Venkataram, S., Zhou, Y. & Fraser, H. B. High-resolution mapping of *cis*-regulatory variation in budding yeast. Proc. Natl. Acad. Sci. U.S.A. 114, (2017).

96. Peter, J. et al. Genome evolution across 1,011 Saccharomyces cerevisiae isolates. Nature 556, 339–344 (2018).

97. Renganaath, K. et al. Systematic identification of cis-regulatory variants that cause gene expression differences in a yeast cross. eLife 9, e62669 (2020).

98. Kuras, L. et al. Dual Regulation of the Met4 Transcription Factor by Ubiquitin-Dependent Degradation and Inhibition of Promoter Recruitment. Molecular Cell 10, 69–80 (2002).

99. Leroy, C., Cormier, L. & Kuras, L. Independent Recruitment of Mediator and SAGA by the Activator Met4. Molecular and Cellular Biology 26, 3149–3163 (2006).

100. Lin, L., Chamberlain, L., Zhu, L. J. & Green, M. R. Analysis of Gal4-directed transcription activation using Tra1 mutants selectively defective for interaction with Gal4. Proc. Natl. Acad. Sci. U.S.A. 109, 1997–2002 (2012).

101. Chandrasekaran, S. & Skowyra, D. The emerging regulatory potential of SCFMet30-mediated polyubiquitination and proteolysis of the Met4 transcriptional activator. Cell Div 3, 11 (2008).

102. Tang, Z., Somia, N., Yu, Y. & Koo, P. K. Evaluating the representational power of pre-trained DNA language models for regulatory genomics. Genome Biol 26, 203 (2025).

103. De Boer, C. G. & Taipale, J. Hold out the genome: a roadmap to solving the cis-regulatory code. Nature 625, 41–50 (2024).

104. Hendrycks, D. & Gimpel, K. Gaussian Error Linear Units (GELUs). Preprint at 10.48550/ARXIV.1606.08415 (2016).

105. Kurtz, S. et al. Versatile and open software for comparing large genomes. Genome Biol 5, R12 (2004).

106. Baker, D. N. & Langmead, B. Genomic sketching with multiplicities and locality-sensitive hashing using Dashing 2. Genome Res. gr.277655.123 (2023) doi:10.1101/gr.277655.123.

107. Ertl, O. SetSketch: Filling the Gap between MinHash and HyperLogLog. 10.48550/ARXIV.2101.00314 (2021) doi:10.48550/ARXIV.2101.00314.

108. Ertl, O. ProbMinHash – A Class of Locality-Sensitive Hash Algorithms for the (Probability) Jaccard Similarity. IEEE Trans. Knowl. Data Eng. 1–1 (2020) doi:10.1109/TKDE.2020.3021176.

109. Broder, A. Z. On the resemblance and containment of documents. in Proceedings. Compression and Complexity of SEQUENCES 1997 (Cat. No.97TB100171) 21–29 (IEEE Comput. Soc, Salerno, Italy, 1998). doi:10.1109/SEQUEN.1997.666900.

110. Jaccard, P. Étude comparative de la distribution florale dans une portion des Alpes et des Jura. Bull Soc Vaudoise Sci Nat 37, 547–579 (1901).

111. Wheeler, T. J. et al. Dfam: a database of repetitive DNA based on profile hidden Markov models. Nucleic Acids Research 41, D70–D82 (2012).

112. Pertea, G. & Pertea, M. GFF Utilities: GffRead and GffCompare. F1000Res 9, 304 (2020).

113. Simão, F. A., Waterhouse, R. M., Ioannidis, P., Kriventseva, E. V. & Zdobnov, E. M. BUSCO: assessing genome assembly and annotation completeness with single-copy orthologs. Bioinformatics 31, 3210–3212 (2015).

114. Devlin, J., Chang, M.-W., Lee, K. & Toutanova, K. BERT: Pre-training of Deep Bidirectional Transformers for Language Understanding. Preprint at 10.48550/ARXIV.1810.04805 (2018).

115. Mallet, V. & Vert, J.-P. Reverse-complement equivariant networks for DNA sequences. Advances in neural information processing systems 34, 13511–13523 (2021).

116. Zhou, H., Shrikumar, A. & Kundaje, A. Towards a better understanding of reverse-complement equivariance for deep learning models in genomics. in Machine Learning in Computational Biology 1–33 (PMLR, 2022).

117. Kingma, D. P. & Ba, J. Adam: A Method for Stochastic Optimization. Preprint at 10.48550/ARXIV.1412.6980 (2014).

118. Schneider, T. D., Stormo, G. D., Gold, L. & Ehrenfeucht, A. Information content of binding sites on nucleotide sequences. Journal of Molecular Biology 188, 415–431 (1986).

119. Schneider, T. D. & Stephens, R. M. Sequence logos: a new way to display consensus sequences. Nucl Acids Res 18, 6097–6100 (1990).

120. Wolf, T., et al. HuggingFace’s Transformers: State-of-the-art Natural Language Processing. Preprint at 10.48550/ARXIV.1910.03771 (2019).

121. Van der Maaten, L. & Hinton, G. Visualizing data using t-SNE. Journal of machine learning research 9, (2008).

122. Arita, Y. et al. A genome-scale yeast library with inducible expression of individual genes. Molecular Systems Biology 17, e10207 (2021).

123. Balakrishnan, R. et al. YeastMine—an integrated data warehouse for Saccharomyces cerevisiae data as a multipurpose tool-kit. Database 2012, (2012).

124. Turco, G. et al. Global analysis of the yeast knockout phenome. Sci. Adv. 9, eadg5702 (2023).

125. Hou, J. et al. The Hidden Complexity of Mendelian Traits across Natural Yeast Populations. Cell Reports 16, 1106–1114 (2016).

126. Li, H. Aligning sequence reads, clone sequences and assembly contigs with BWA-MEM. Preprint at 10.48550/ARXIV.1303.3997 (2013).

127. Li, H. et al. The Sequence Alignment/Map format and SAMtools. Bioinformatics 25, 2078–2079 (2009).

128. Quinlan, A. R. & Hall, I. M. BEDTools: a flexible suite of utilities for comparing genomic features. Bioinformatics 26, 841–842 (2010).

129. Dobin, A. et al. STAR: ultrafast universal RNA-seq aligner. Bioinformatics 29, 15–21 (2013).

130. Picard toolkit. Broad Institute, GitHub repository (2019).

131. Stovner, E. B. & Sætrom, P. PyRanges: efficient comparison of genomic intervals in Python. Bioinformatics 36, 918–919 (2020).

132. Significance of nucleotide sequence alignments: a method for random sequence permutation that preserves dinucleotide and codon usage. Molecular Biology and Evolution 10.1093/oxfordjournals.molbev.a040370 (1985) doi:10.1093/oxfordjournals.molbev.a040370.

133. Holm, S. A simple sequentially rejective multiple test procedure. Scandinavian journal of statistics 65–70 (1979).

