## Supplemental Table for "Predicting dynamic expression patterns in budding yeast with a fungal DNA language model"

**Supplemental Table 1.** Composition of the phosphate-limited, minimal chemically defined medium used in continuous culture

| **Componenet** | **Quantity** |
|---|---|
| **1000X metals stock solution in 1L of sterile water** | |
| Boric Acid | 500 mg |
| Copper Sulfate | 40 mg |
| Potassium Iodide | 100 mg |
| Ferric Chloride | 200 mg |
| Manganese Sulfate | 400 mg |
| Sodium Molybdate | 200 mg |
| Zinc Sulfate | 400 mg |
| **1000X vitamins stock solution in 1L of sterile water** | |
| Biotin | 2 mg |
| Calcium Pantothenate(B-5) | 400 mg |
| Folic acid | 2 mg |
| Inositol (myo-inositol) | 2000 mg |
| Niacin (nicotinic acid) | 400 mg |
| p-aminobenzoic acid | 200 mg |
| Pyridoxine hydrochloric acid | 400 mg |
| Riboflavin | 200 mg |
| Thiamine | 400 mg |
| **10X salts stock solution in 2L of sterile water** | |
| calcium chloride | 20g |
| sodium chloride | 20g |
| magnesium sulfate | 100g |
| ammonium sulfate | 1000g |
| potassium chloride | 200g |
