## Supplemental Figures for "Predicting dynamic expression patterns in budding yeast with a fungal DNA language model"

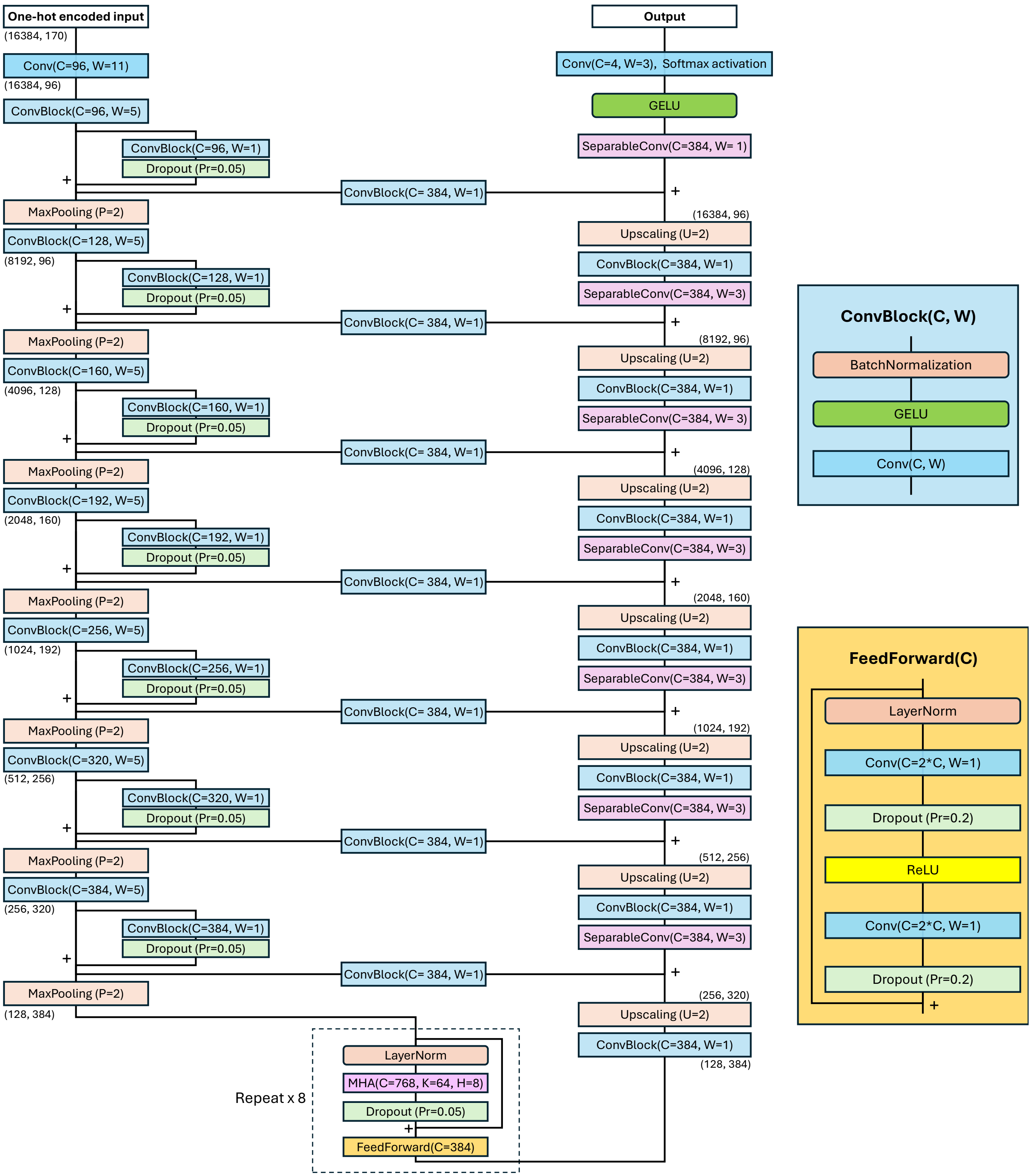


**Figure 1-figure supplement 1.** Architecture of the Shorkie LM model. The 13.7-million-parameter model predicts per-base nucleotide probabilities from genomic windows of 16,384 bp. Input sequences are augmented with stochastic shifts and passed through a 1D convolution layer (kernel size = 11, filters = 96), generating a 16,384 × 96 feature map. The encoder consists of seven residual down-sampling blocks, reducing the map to 128 × 384. This is followed by an eight-layer Transformer bottleneck that integrates long-range sequence context. The decoder mirrors the encoder with seven up-sampling stages, U-Net skip connections, and depthwise-separable convolutions, ultimately restoring the sequence to its original length of 16,384 positions. A final 1×1 convolutional layer and a softmax activation function across four channels produce per-position nucleotide probability distributions for A/C/G/T.

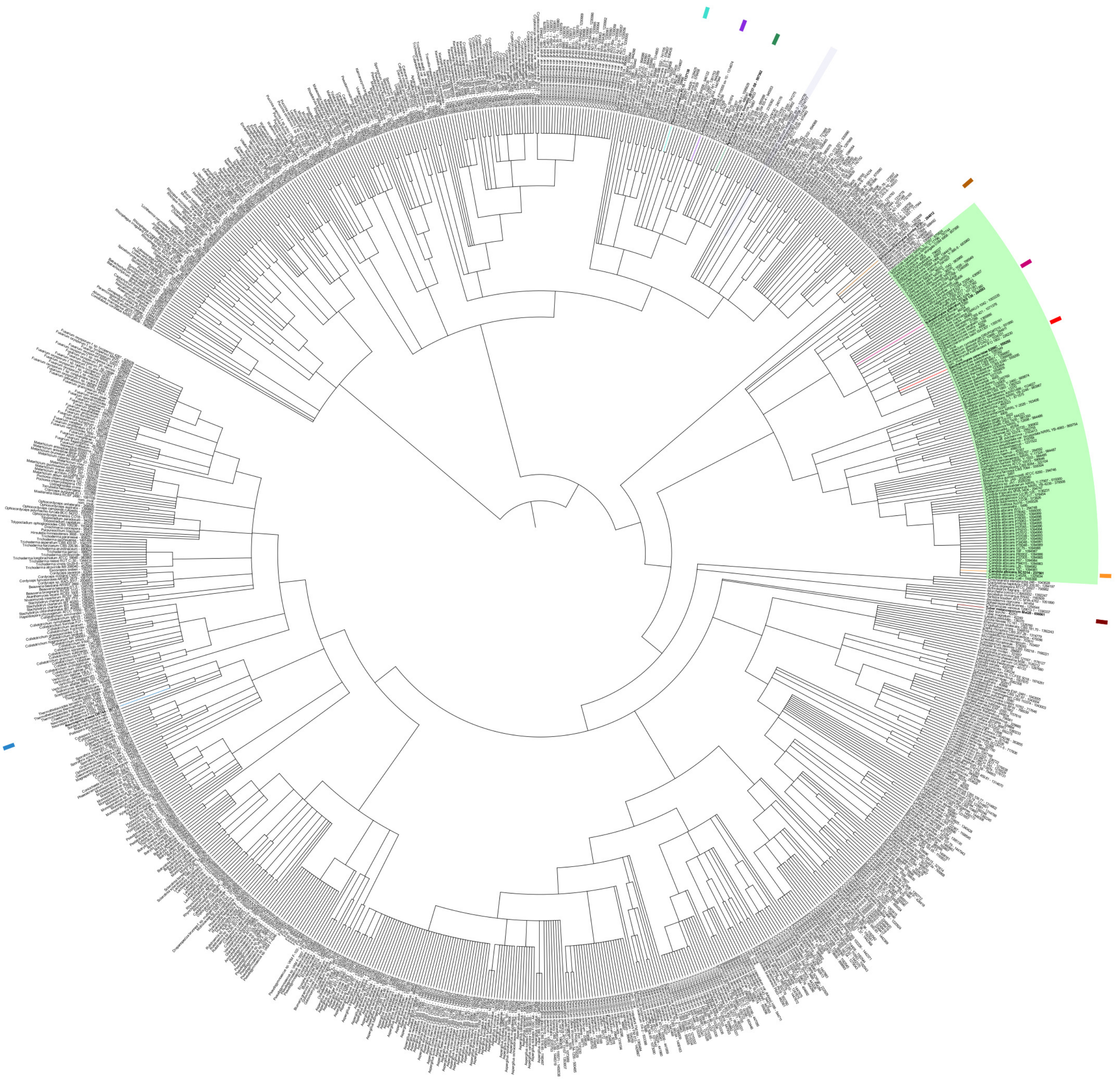

**Figure 1-figure supplement 2.** Phylogenetic tree of 1,341 fungal genomes. Species were assigned to NCBI Taxonomy IDs using the ETE Toolkit, and a minimum spanning tree of the target taxa was constructed with ete-ncbiquery. The tree, formatted in Newick, was visualized using iTOL v5 with a circular layout and annotated with color-coding for taxonomic groups. The order Saccharomycetales is highlighted in light green. Example annotated species include *Pleurotus ostreatus* (oyster mushroom) in cyan, *Lentinula edodes* (shiitake) in purple, *Agaricus bisporus* (white button mushroom) in green, *Tuber melanosporum* (black truffle) in brown, *Schizosaccharomyces pombe* (fission yeast) in orange, and *Saccharomyces cerevisiae* (brewer’s yeast) in red.

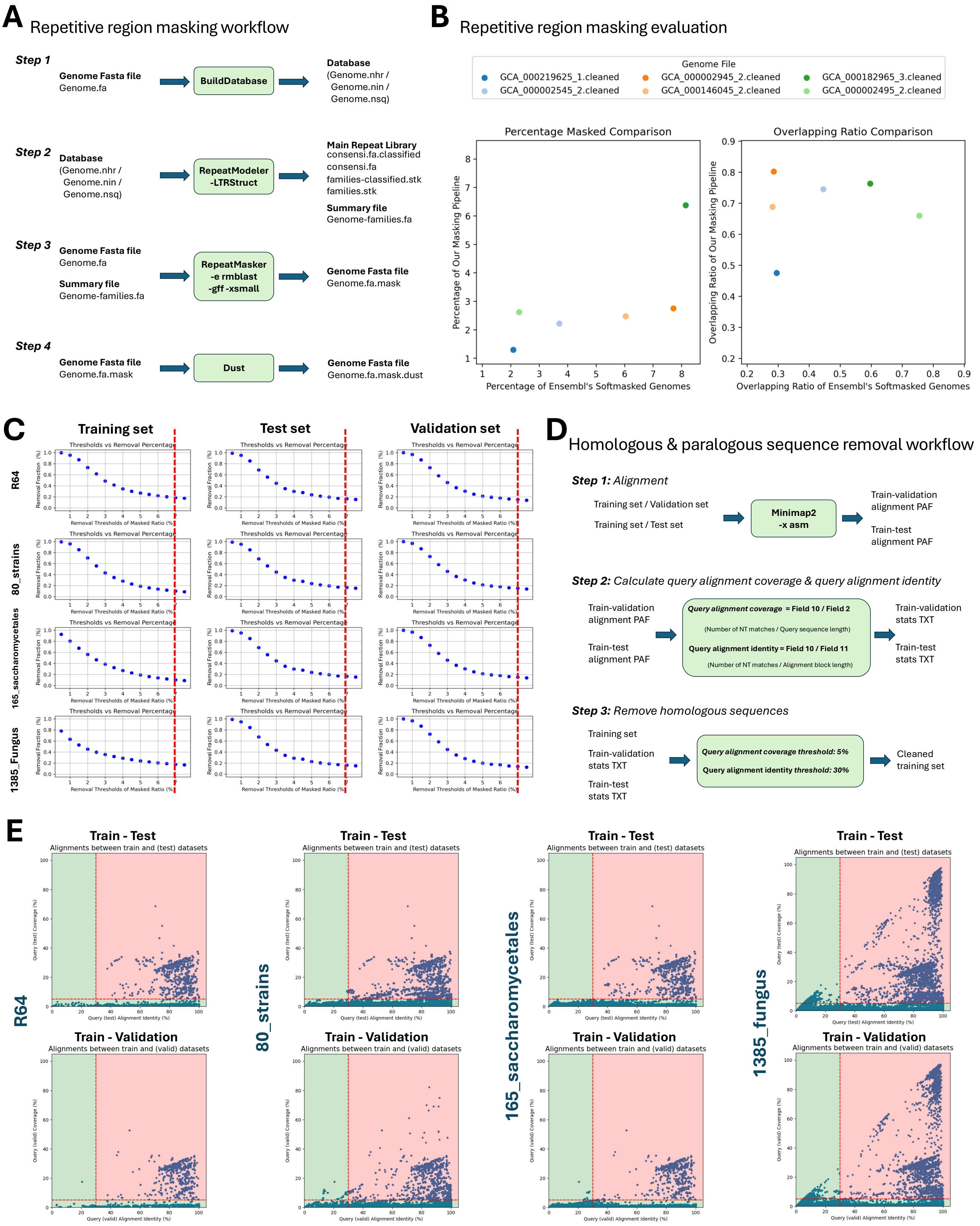
A
Repetitive region masking workflow
Step 1
Genome Fasta file
Genome.fa
BuildDatabase
Database
(Genome.nhr /
Genome.nin /
Genome.nsq)
Step 2
Database
(Genome.nhr /
Genome.nin /
Genome.nsq)
RepeatModeler
-LTRStruct
Main Repeat Library
consensi.fa.classified
consensi.fa
families-classified.stk
families.stk
Summary file
Genome-families.fa
Step 3
Genome Fasta file
Genome.fa
Summary file
Genome-families.fa
RepeatMasker
-e rmblast
-gff -xsmall
Genome Fasta file
Genome.fa.mask
Step 4
Genome Fasta file
Genome.fa.mask
Dust
Genome Fasta file
Genome.fa.mask.dust
B
Repetitive region masking evaluation
Genome File
GCA_000219625_1.cleaned
GCA_000002545_2.cleaned
GCA_000002945_2.cleaned
GCA_000146045_2.cleaned
GCA_000182965_3.cleaned
GCA_000002495_2.cleaned
Percentage Masked Comparison
Percentage of Our Masking Pipeline
Percentage of Ensembl's Softmasked Genomes
Overlapping Ratio Comparison
Overlapping Ratio of Our Masking Pipeline
Overlapping Ratio of Ensembl's Softmasked Genomes
C
Training set
Test set
Validation set
R64
80_strains
165_saccharomycetales
1385_Fungus
Thresholds vs Removal Percentage
Removal Fraction (%)
Removal Thresholds of Masked Ratio (%)
D
Homologous & paralogous sequence removal workflow
Step 1: Alignment
Training set / Validation set
Training set / Test set
Minimap2
-x asm
Train-validation
alignment PAF
Train-test
alignment PAF
Step 2: Calculate query alignment coverage & query alignment identity
Train-validation
alignment PAF
Train-test
alignment PAF
Query alignment coverage = Field 10 / Field 2
(Number of NT matches / Query sequence length)
Query alignment identity = Field 10 / Field 11
(Number of NT matches / Alignment block length)
Train-validation
stats TXT
Train-test
stats TXT
Step 3: Remove homologous sequences
Training set
Train-validation
stats TXT
Train-test
stats TXT
Query alignment coverage threshold: 5%
Query alignment identity threshold: 30%
Cleaned
training set
E
R64
80_strains
165_saccharomycetales
1385_fungus
Train - Test
Alignments between train and (test) datasets
Query (test) Coverage (%)
Query (test) Alignment Identity (%)
Train - Validation
Alignments between train and (valid) datasets
Query (valid) Coverage (%)
Query (valid) Alignment Identity (%)

**Figure 1-figure supplement 3.** Shorkie LM data preprocessing and homology filtering. **(A)** Two-tiered repeat masking approach: Each fungal genome from Ensembl was soft-masked using a custom library built with RepeatModeler v2.0 (incorporating RepeatScout, RECON, and LTR_retriever) merged with Dfam entries. RepeatMasker (RMBlast) was used with the "`-xsmall`" and "`-gff`" options to soft-mask interspersed repetitive elements, while residual low-complexity regions were masked using DUST from the MEME suite (with a score threshold of 20). **(B)** Comparison of the custom repeat masks against the original soft-masking from Ensembl in six representative genome assemblies. **(C)** Fully masked genomes were divided into 16,384 bp windows with a 4,096 bp stride; windows containing >7% masked content were excluded, resulting in the removal of approximately 20% of windows across all data splits. **(D)** Homology filtering to avoid data leakage: All training windows were aligned against the validation and test sets using Minimap2 ("`-x asm`"). For each alignment, coverage (percentage of query length covered) and identity (percentage of alignment matching the query) were computed. Any training window with ≥ 5% coverage and ≥ 30% identity to any validation or test window was discarded. **(E)** Scatter plots of coverage versus identity for all training-validation and training-test alignments, highlighting the thresholds used to define non-redundant training sequences.

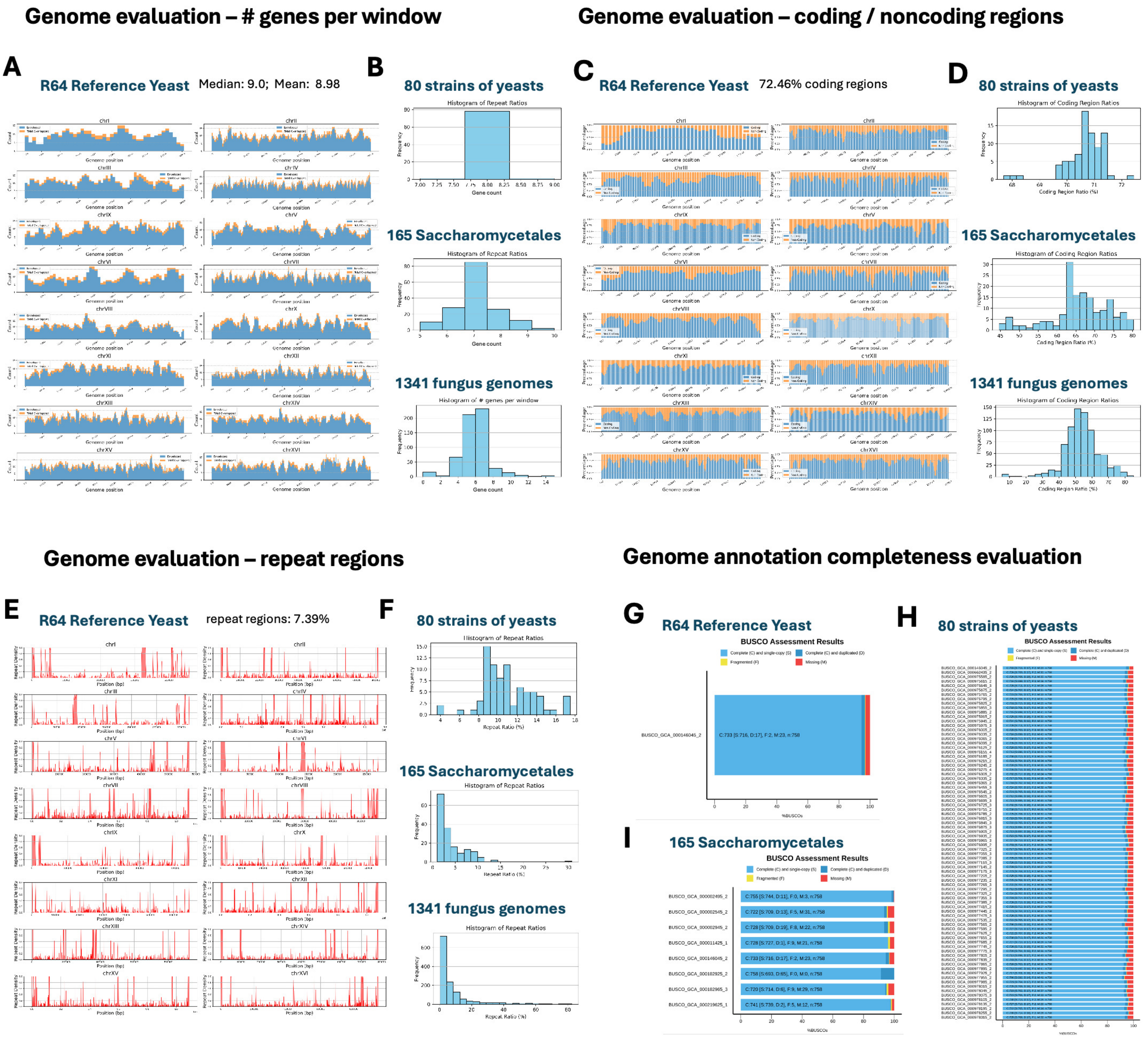


**Figure 1-figure supplement 4.** Window-based analysis of genomic features and BUSCO annotation completeness across fungal assemblies. Genomes were partitioned into 16,384 bp windows with a 4,096 bp stride, and the following analyses were performed: **(A)** gene count per window in the *S. cerevisiae* reference genome; **(B)** distribution of gene counts across 80 *S. cerevisiae* strains, 165 Saccharomycetales genomes, and 1,341 fungal assemblies; **(C)** ratio of coding-to-noncoding sequences per window in the *S. cerevisiae* reference genome; **(D)** distribution of coding-to-noncoding ratios across the three genome sets; **(E)** fraction of each window that is soft-masked as repetitive sequence in the *S. cerevisiae* reference genome; **(F)** distribution of repeat-masked fractions across the three genome sets; **(G–I)** BUSCO v5 (fungi_odb10) completeness scores: percentages of complete, fragmented, and missing orthologs for **(G)** the *S. cerevisiae* reference genome, **(H)** 80 *S. cerevisiae* strains, and **(I)** 165 Saccharomycetales genomes.

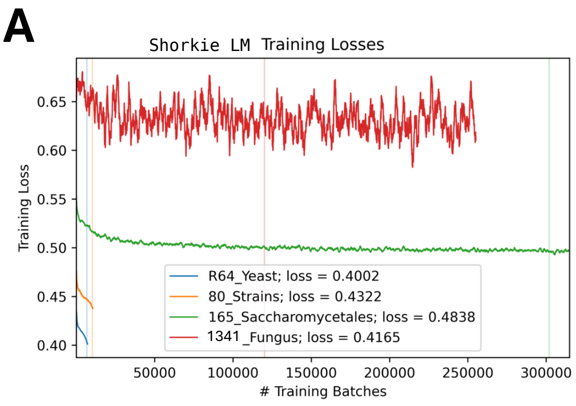


**B** Model architecture comparison

| Model | Architecture | Total Params | Key Features |
|---|---|---|---|
| Conv_Small | Conv1D + Residual blocks | 320,708 | Smaller conv width (64 channels). |
| Conv_big | Conv1D + Residual blocks | 3,642,116 | Larger conv width (256 channels). |
| Unet_small | U-Net + Transformers | 13,665,828 | Smaller conv width (96 channels) + 8 transformer blocks |
| Unet_big | U-Net + Transformers | 71,790,564 | Smaller conv width (384 channels) + 11 transformer blocks |

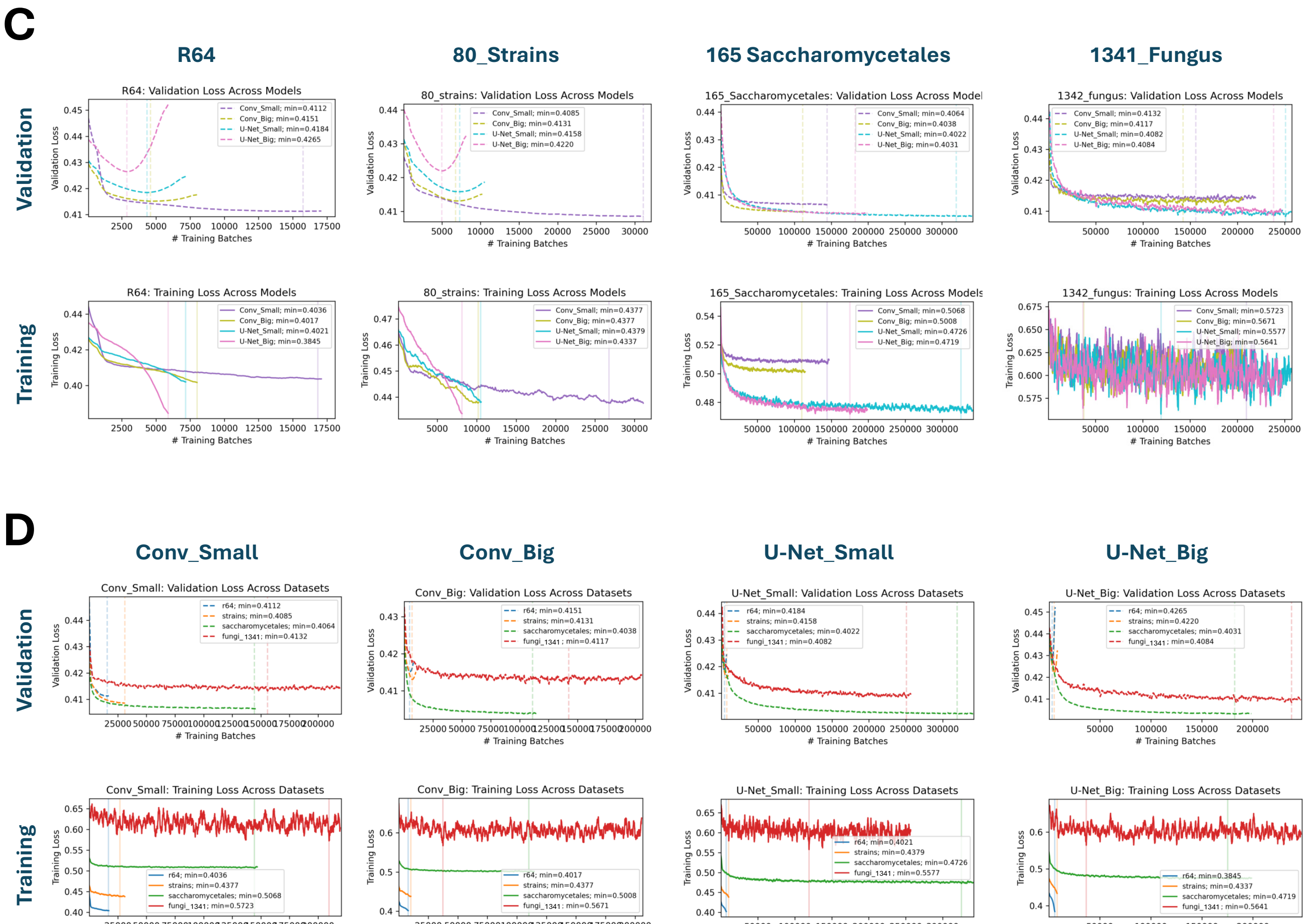


**Figure 1-figure supplement 5.** Evaluation of Shorkie LM training across evolutionary divergence and model architectures. **(A)** Training and validation loss curves for models trained on four fungal genomic datasets, ordered by increasing evolutionary divergence: R64 (blue; least divergent), 80_strains (orange), 165_Saccharomycetales (green), and 1341_Fungus (red; most divergent). Solid lines represent training loss, while dashed lines represent validation loss across training iterations. **(B)** Overview of the four neural network architectures assessed: (1) Conv_Small (64-channel Conv1D with residual blocks), (2) Conv_Big (256-channel Conv1D with residual blocks), (3) U-Net_Small (U-Net backbone with eight transformer blocks and 96-to-384-channel convolutions), and (4) U-Net_Big (U-Net backbone with 11 transformer blocks, 384-to-768-channel convolutions, and larger convulational widths). **(C)** Final training and validation loss values for each architecture across the four datasets. The x-axis represents datasets (R64,

80_strains, 165_Saccharomycetales, 1341_Fungus), and colored lines denote different architectures. Solid lines indicate training loss, while dotted lines indicate validation loss. **(D)** Final training and validation loss values for each dataset across the four architectures. The x-axis represents architectures (Conv_Small, Conv_Big, U-Net_Small, U-Net_Big), with colored lines corresponding to datasets. Solid curves represent training loss, while dotted curves represent validation loss. For visualization, loss traces are smoothed with a centered moving average (window = 21 training steps).

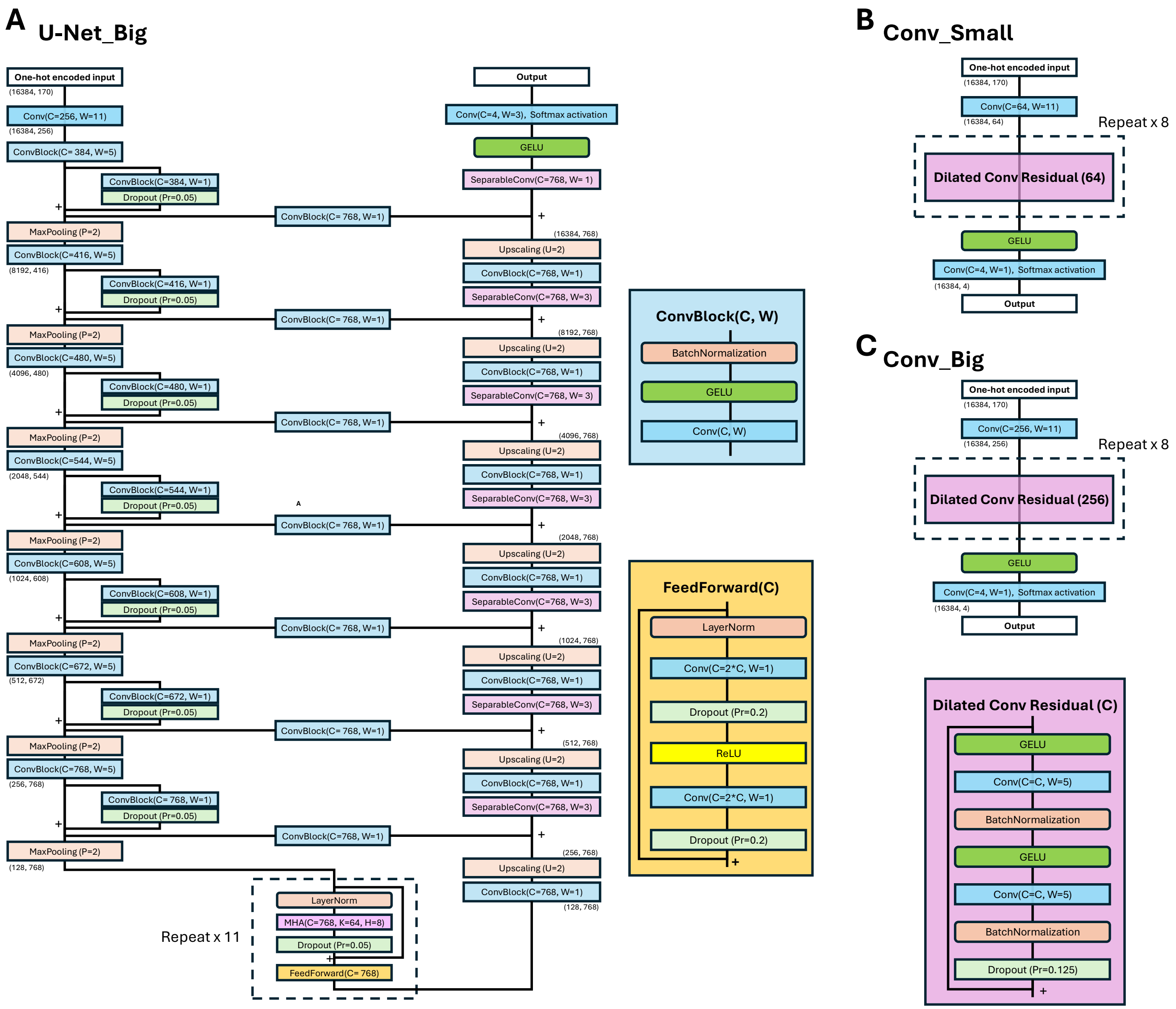


**Figure 1-figure supplement 6.** Schematic overviews of the three additional neural network architectures evaluated for Shorkie LM (U-Net_Small is described in Figure 1-figure supplement 1). **(A)** U-Net_Big: A U-Net–style encoder–decoder with 11 transformer blocks interleaved at the bottleneck, 384 convolutional channels per layer, skip connections between matching resolution stages, and a total of ~71.8 million parameters. **(B)** Conv_Small: A compact residual 1D convolutional network comprising four residual blocks with 64-channel convolutions, totaling ~320.7 thousand parameters. **(C)** Conv_Big: A deeper residual 1D convolutional network with four residual blocks using 256-channel convolutions, totaling ~3.64 million parameters. Each panel illustrates the flow of data from raw one-hot encoded sequence input through successive convolutional and, where applicable, transformer layers, to the final prediction head.

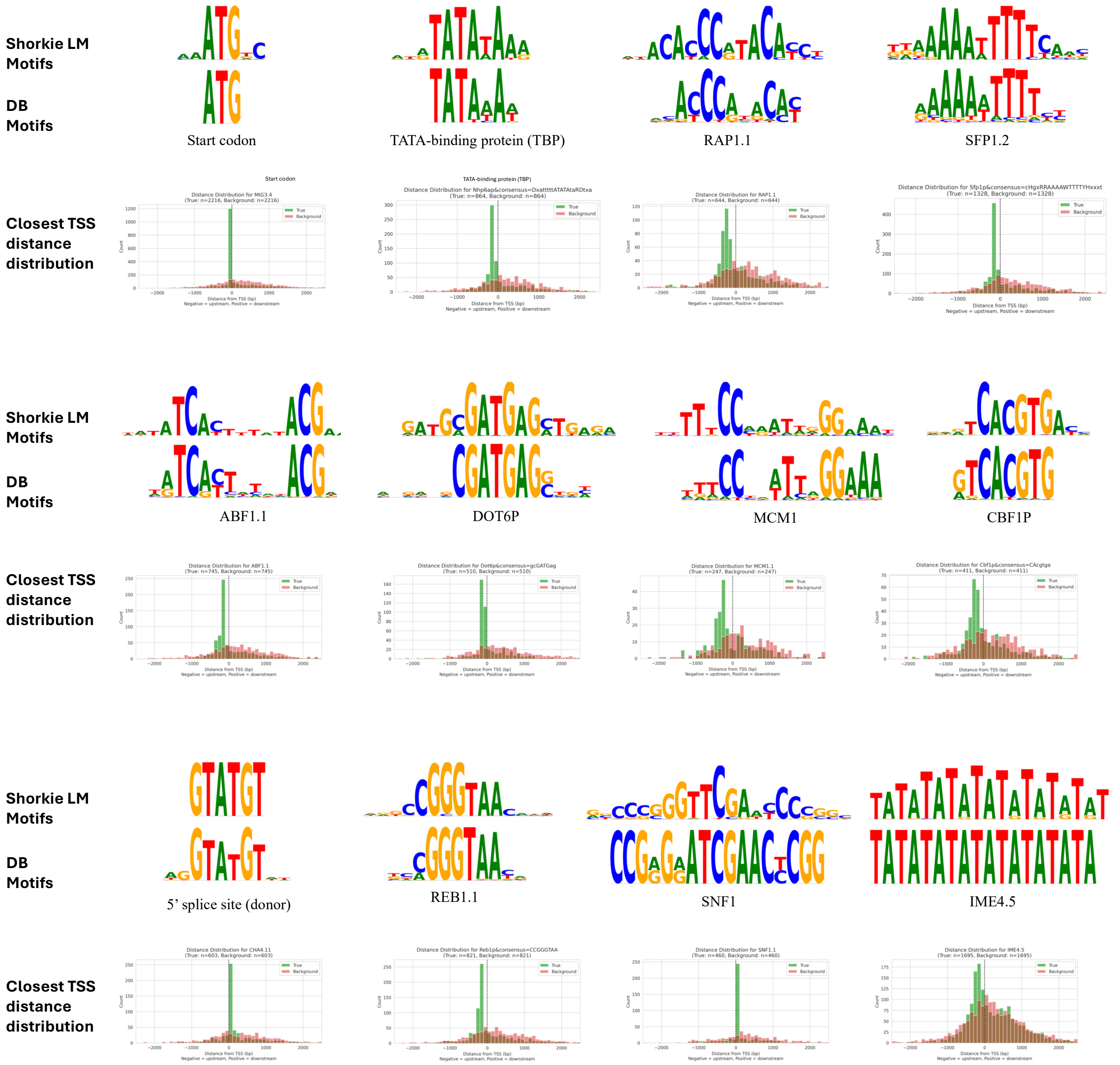


**Figure 2-figure supplement 1.** *De novo* motif discovery by Shorkie LM and positional enrichment relative to transcription start sites (TSSs). Using TF-MoDISco-Lite with importance scores derived from the *S. cerevisiae* reference genome, we recovered and clustered seqlets corresponding to the start codon, TATA-binding protein (TBP), Rap1.1, Sfp1.2, Abf1.1, Dot6p, Mcm1, Cbf1p, the 5′ splice-site donor motif, Reb1.1, Snf1 and Ime4.5. For each clustered motif, we plotted histograms of each motif’s detected seqlet distances to the nearest TSS versus a shuffled background.

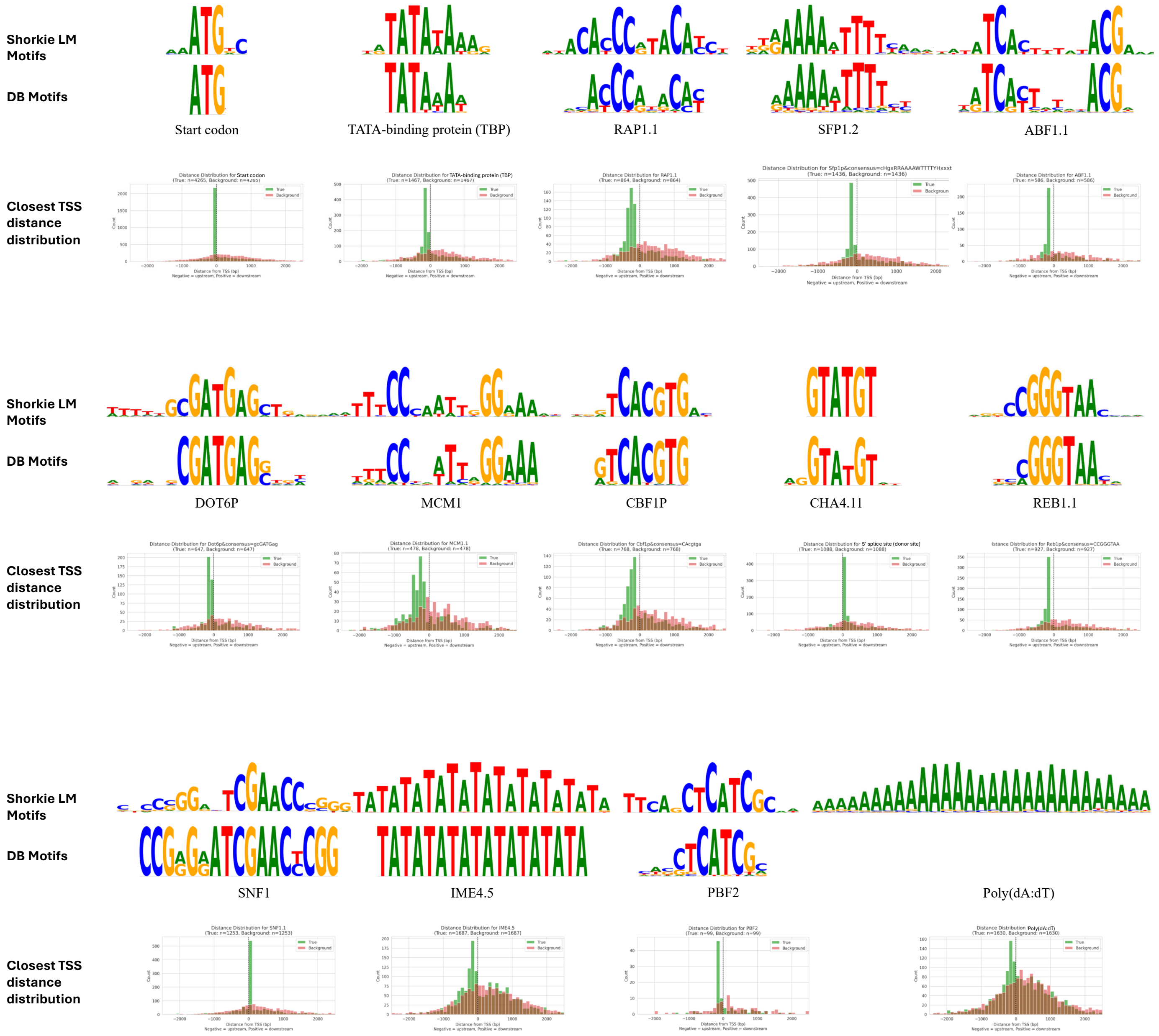


**Figure 2-figure supplement 2.** *De novo* motif discovery by Shorkie LM and positional enrichment relative to transcription start sites (TSSs). Using TF-MoDISco-Lite with importance scores derived from five Saccharomycetales genomes, we recovered and clustered seqlets corresponding to the start codon, TATA-binding protein, Rap1.1, Sfp1.2, Abf1.1, Dot6p, Mcm1, Cbf1p, the 5′ splice-site donor motif, Reb1.1, Snf1, Ime4.5, Pbf2, and Poly(dA:dT). For each clustered motif, we plotted histograms of each motif's detected seqlet distances to the nearest TSS versus a shuffled background.

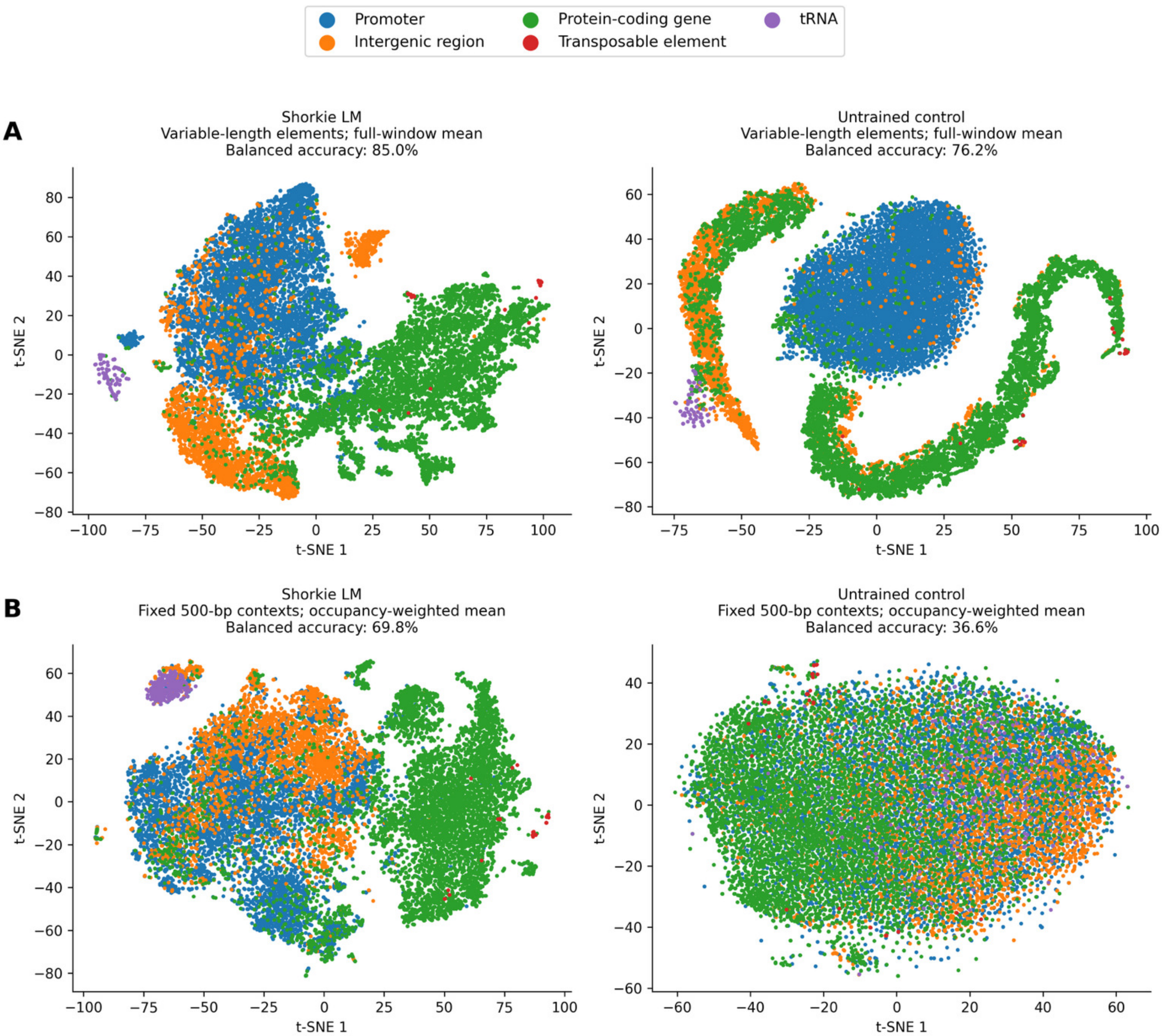


**Figure 2-figure supplement 3.** Element length and padding in the Figure 2E embeddings. t-SNE projections of embeddings of 16,378 annotated genomic elements (promoters, protein-coding genes, intergenic regions, tRNAs and transposable elements) from Shorkie LM (left) and an untrained network with the same architecture (right). (A) The extraction without element length control: each element centered in a 16,384-bp window, zero-padded and mean-pooled over the full window (median 73 bp for tRNAs, 3,468 bp for transposable elements). (B) 500 bp of native genomic context centered on each element, with occupancy-weighted pooling. Embeddings are the output of the first transformer block for both models.

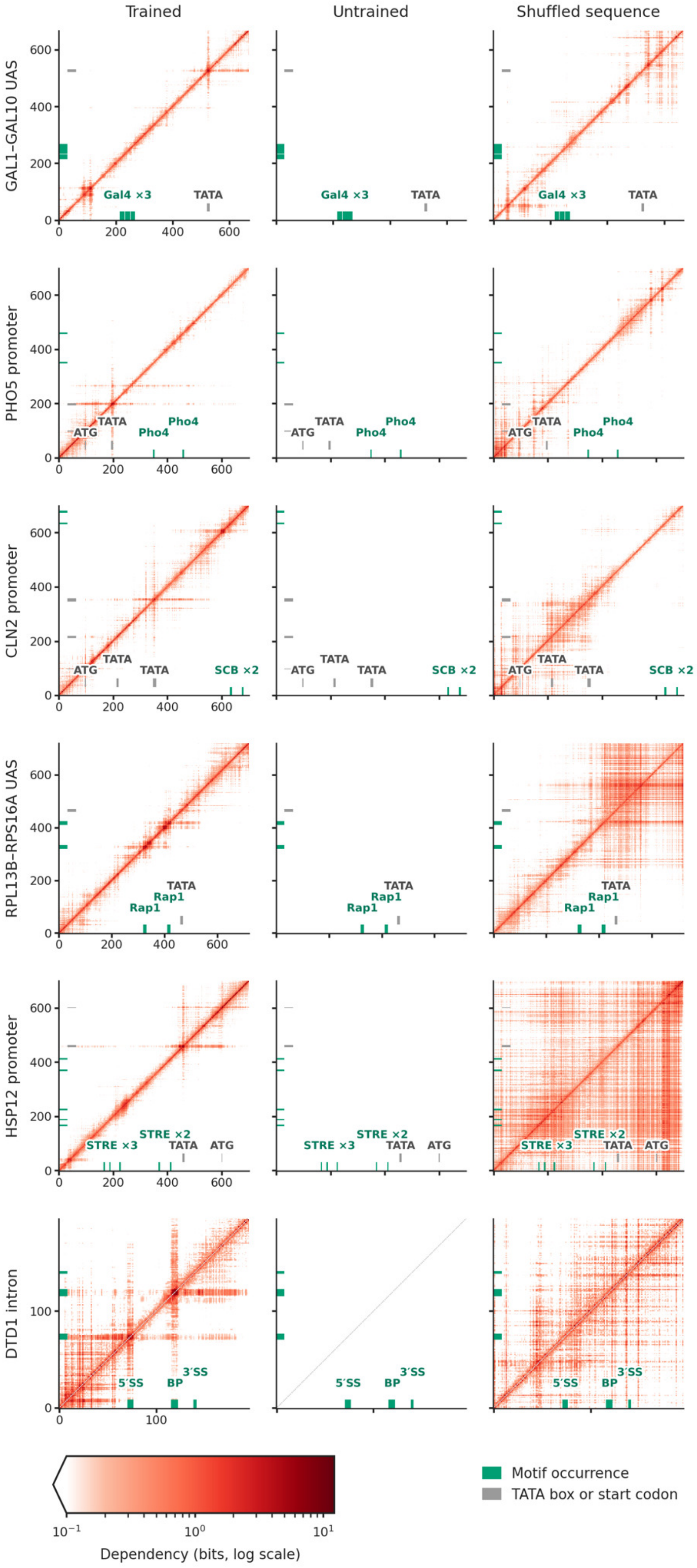
Trained
Untrained
Shuffled sequence
GAL1-GAL10 UAS
Gal4 ×3
TATA
PHO5 promoter
ATG
TATA
Pho4
Pho4
CLN2 promoter
ATG
TATA
TATA
SCB ×2
RPL13B-RPS16A UAS
TATA
Rap1
Rap1
HSP12 promoter
STRE ×3
STRE ×2
TATA
ATG
DTD1 intron
5′SS
BP
3′SS
Motif occurrence
TATA box or start codon
Dependency (bits, log scale)

**Figure 2-figure supplement 4.** Nucleotide dependency maps at six selected regulatory loci. Values show the largest absolute change in predicted log2-odds at each target position (x axis) after mutation of a query position (y axis). Columns compare Shorkie LM on native sequence, an untrained model with the same architecture, and Shorkie LM on a dinucleotide-shuffled locus. All maps use a shared logarithmic scale (0.1–12.2 bits). Teal and grey marks and labels indicate motif occurrences and TATA boxes (TATA) or start codons (ATG), respectively; BP, branch point; 5'SS and 3'SS, 5' and 3' splice sites.

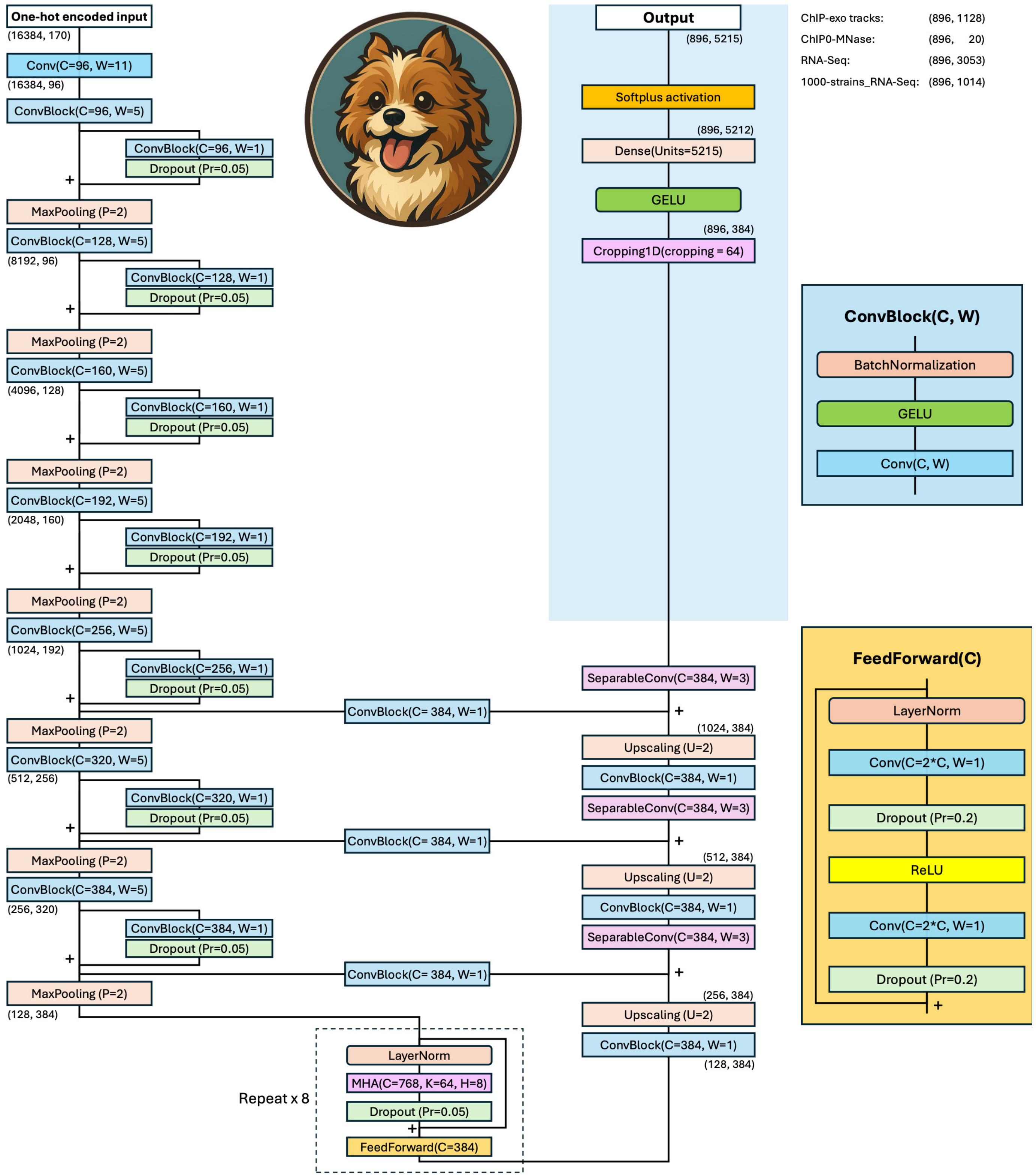


**Figure 3-figure supplement 1.** Shorkie model architecture. The Shorkie model, with 14.3 million parameters, processes one-hot-encoded 16,384 bp genomic windows and predicts 5,215 distinct genomic tracks at a resolution of 16 bp. The model architecture begins with a 1D convolutional block (96 filters, kernel size = 11) followed by seven residual down-sampling stages, reducing the feature map from 16,384 × 96 to 128 × 384. A transformer bottleneck with eight layers at 128 × 384 integrates long-range dependencies. The decoder mirrors the encoder through three up-sampling stages and U-Net-style skip connections, reconstructing a 1,024 × 384 feature map. All encoder, transformer, and decoder layers are initialized with pre-trained weights from the Shorkie language model. In the final prediction head (highlighted in light blue), the feature map is cropped by 64 bins on each side, resulting in a 896 × 384 feature map, which is then passed through a fully connected layer with Softplus activation. The output is a 896 × 5,215 matrix representing track predictions: 1,128 ChIP-exo tracks, 20 ChIP-MNase tracks, 3,053 IDEA RNA-Seq tracks, and 1,014 1000-strain RNA-Seq tracks.

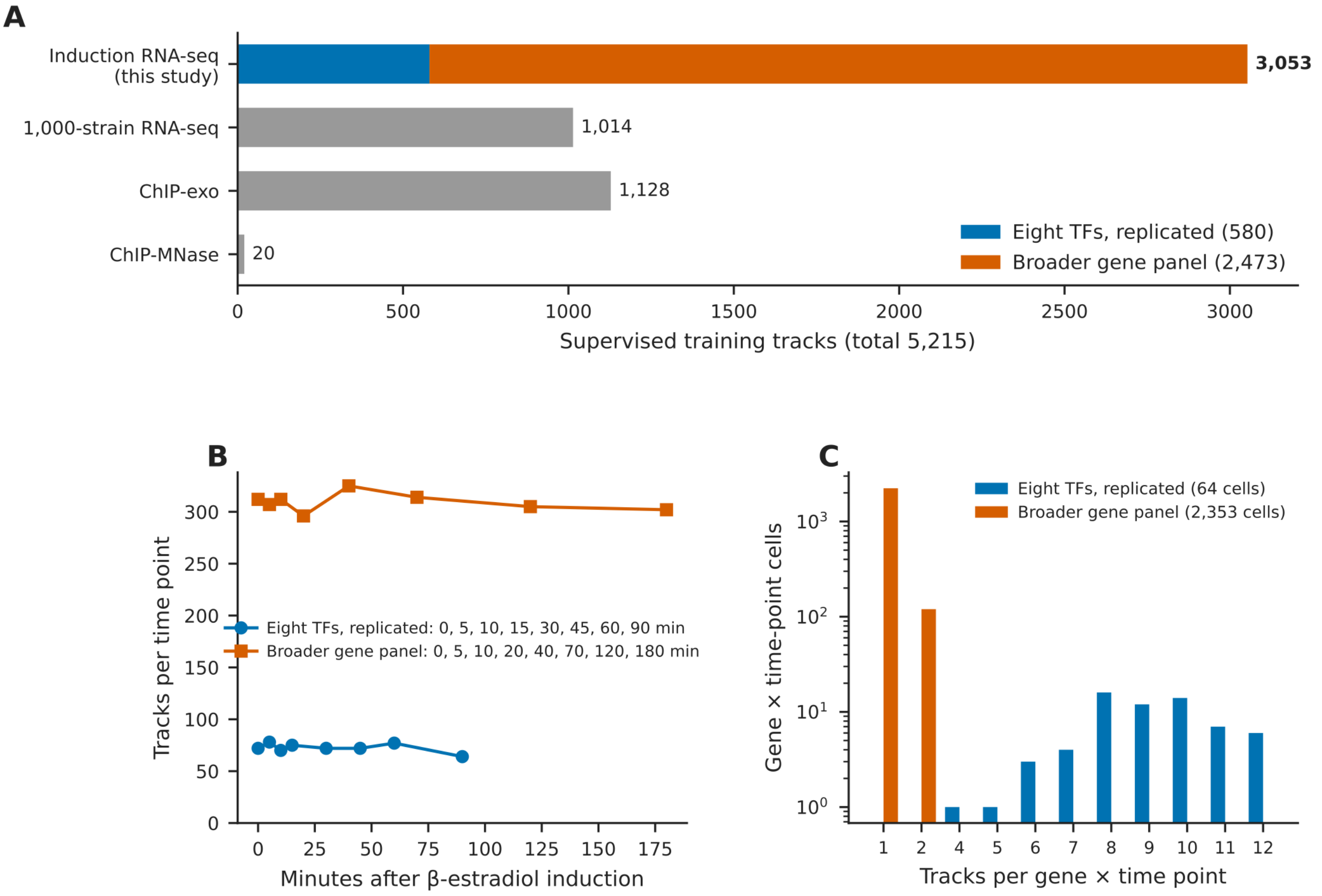


**Figure 3-figure supplement 2.** Composition of the induction RNA-seq resource. (A) The 5,215 supervised training tracks by assay. The 3,053 induction RNA-seq tracks generated in this study comprise the replicated eight-transcription-factor series (580 tracks) and the broader gene panel (2,473 tracks). (B) Tracks per time point; the two series were sampled on different schedules (legend). (C) Tracks per gene and time point (log scale): the eight transcription factors have 4–12 tracks at each time point, whereas most genes of the broader panel have one.

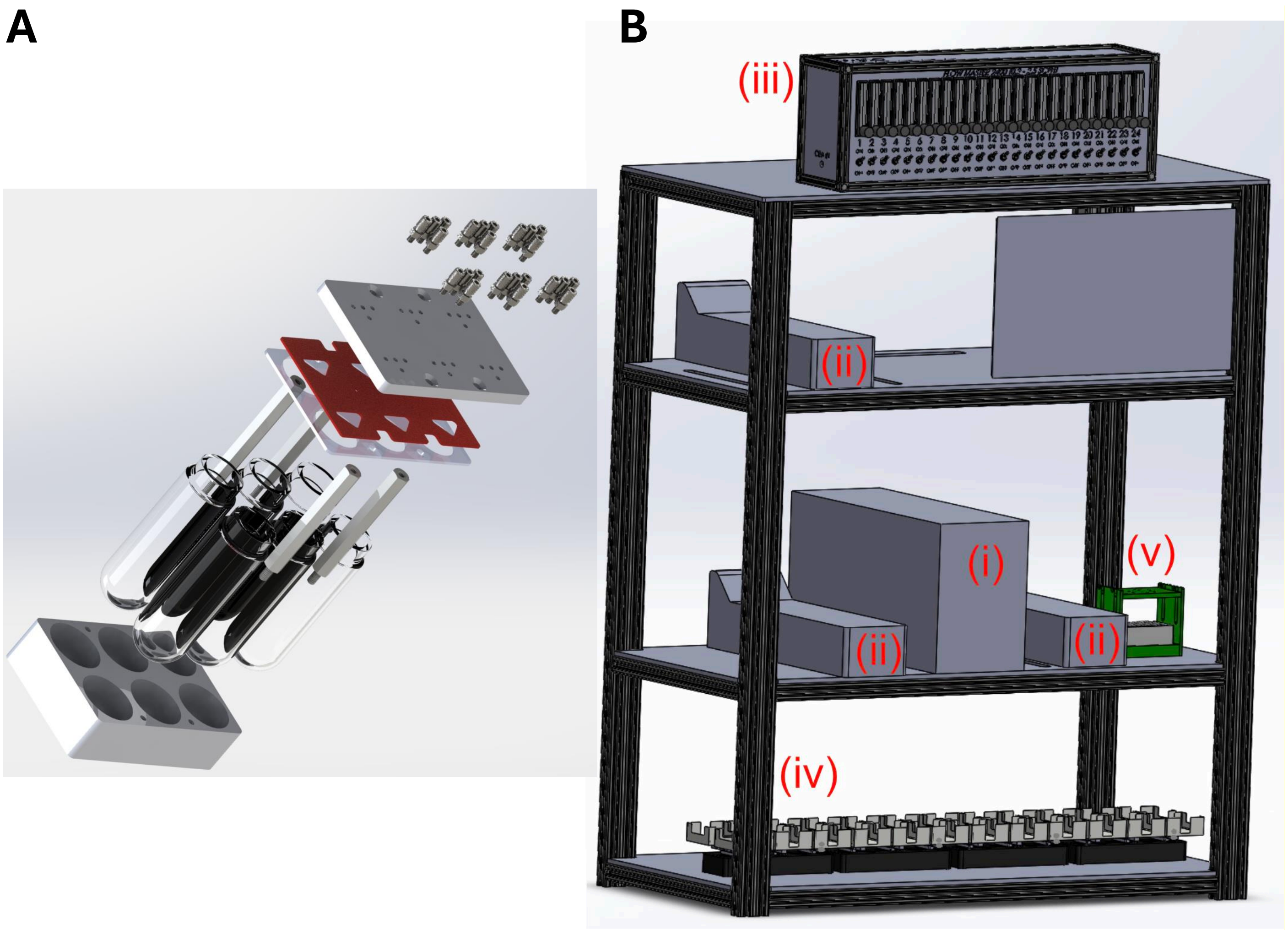


**Figure 3-figure supplement 3.** Overview of ministat array system. **(A)** Yeast were cultured in 100mL round bottom flasks housed in banks of 6-vessels. Each vessel supported connections for media, sampling, and an input for inoculating a bolus of cells or adding estradiol to activate transcriptional induction. **(B)** CAD mock up of major system components highlighted in red (i) 24-vessel manifold combined four 6-vessel manifolds into a complete ministat array (ii) three peristaltic pumps connected to each vessel via individual tubing for media, sampling, and cell/estradiol inputs (iii) air flow regulator with 24 lines whose flow rate was separately controlled (iv) balance system separately weighted the effluent of each ministat to calculate realized dilution rate for adding media to each vessel (v) sampling manifolds which housed plates for cells used for initial inoculation or estradiol for induction and a separate plate where final the samples for sequencing were collected.

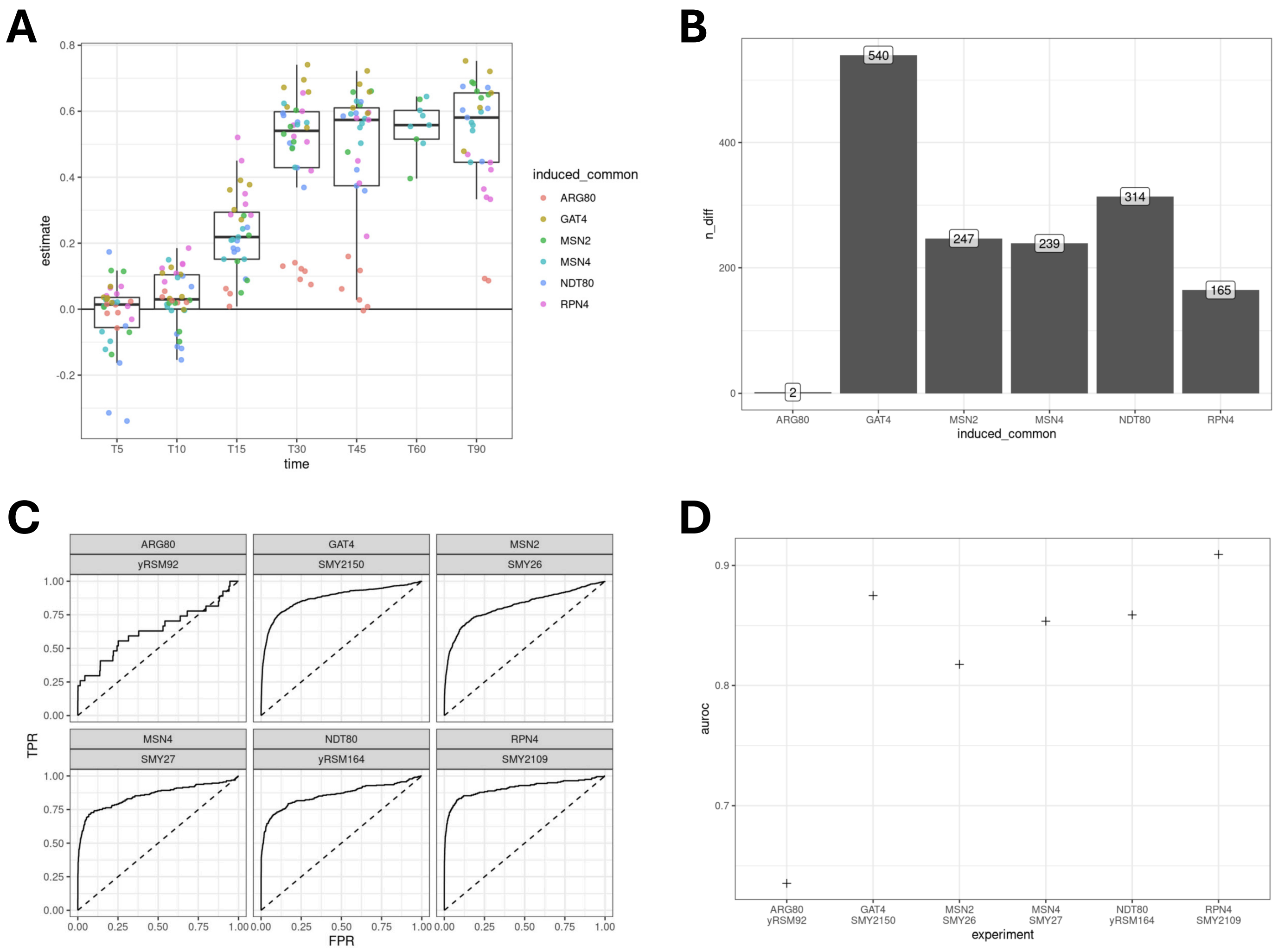


**Figure 3-figure supplement 4.** Ministat Array System Validation. **(A)** Box-and-whisker plots of Pearson correlations between $\log_2$ fold-changes measured by microarray versus RNA-seq for each replicate induction experiment; point colors correspond to the induced TF. **(B)** Bar plot of the number of differentially expressed genes for each TF perturbation. **(C)** ROC curves measuring accuracy in detecting differentially expressed genes for each TF perturbation using the new ministat array system. **(D)** AUROC values for each TF perturbation experiment.

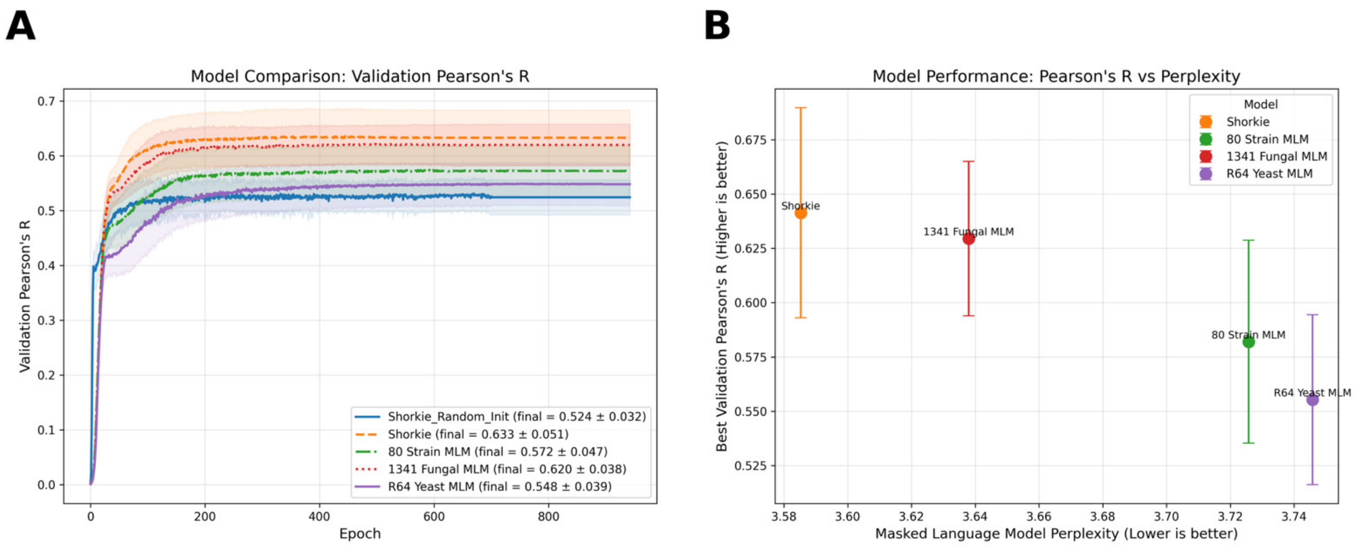


**Figure 3-figure supplement 5.** Downstream RNA-seq prediction performance for Shorkie initialized from LMs pretrained at different evolutionary scopes, and its relationship to LM perplexity. **(A)** Validation learning curves (Pearson's r) across training epochs for an identical supervised Shorkie architecture initialized either from random weights (Shorkie_Random_Init) or from masked language models pretrained on: the *S. cerevisiae* R64 reference only, 80 *S. cerevisiae* strains, 165 Saccharomycetales genomes (Shorkie_LM), or 1,341 fungal genomes. Curves show the mean across 8 cross-validation folds; shaded bands indicate the standard deviation across folds. Legends report the final-epoch mean ± s.d. across folds. **(B)** Best downstream validation performance (maximum Pearson's r across epochs) versus masked-LM perplexity for each LM-pretrained initialization. Points denote the fold-mean best performance; vertical error bars indicate the standard error across the 8 folds.

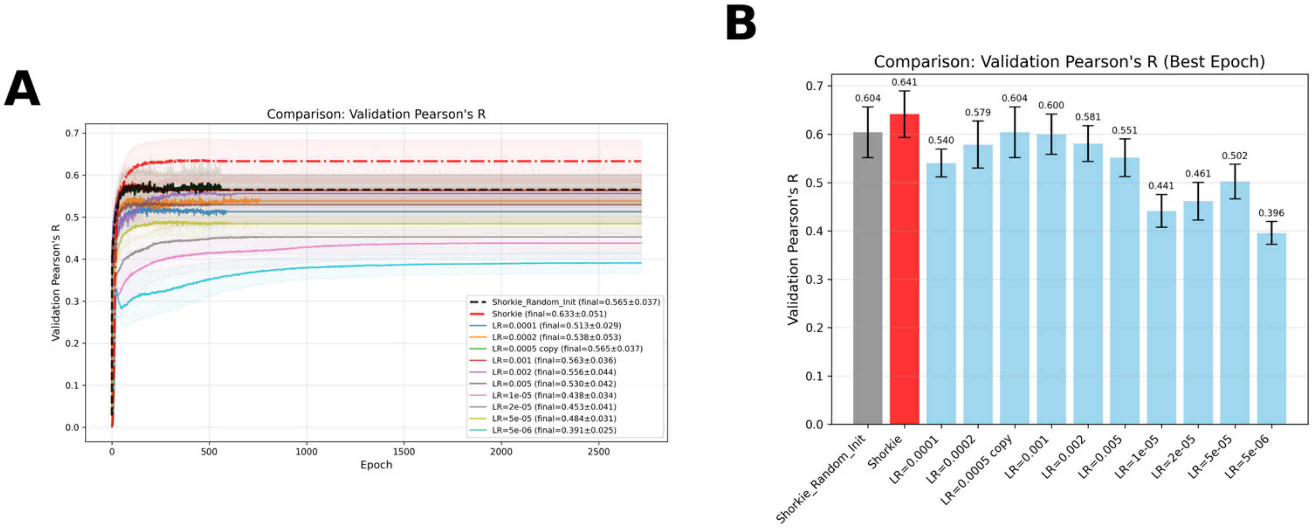


**Figure 3-figure supplement 6.** Learning-rate sweep to establish a strong from-scratch (randomly initialized) Shorkie baseline. **(A)** Validation Pearson correlation (r) versus training epoch for Shorkie fine-tuned from the pretrained Shorkie masked-LM weights (Shorkie; red dash-dot) and for randomly initialized models trained with learning rates spanning $5\times10^{-6}$–$5\times10^{-3}$ (colored curves). The black dashed trajectory indicates the Shorkie_Random_Init setting used throughout the manuscript. Lines and shaded envelopes show the mean ± s.d. across cross-validation folds. **(B)** Peak (best-epoch) validation Pearson r for each setting, summarized as bar heights (mean across folds) with error bars indicating ± s.d.

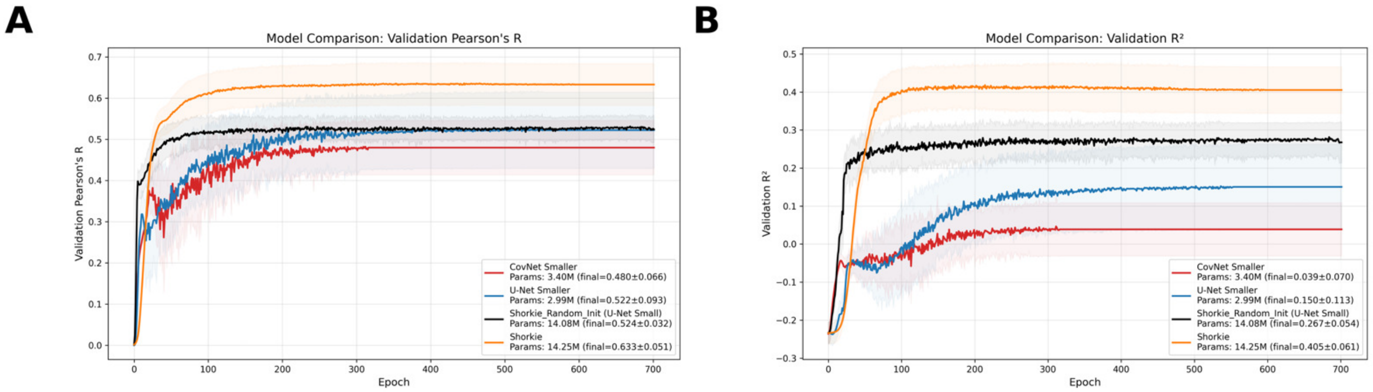


**Figure 3-figure supplement 7.** Reduced-capacity random-initialization baselines for supervised expression prediction. Validation performance of three randomly initialized supervised models: ConvNet_Smaller (3.40M parameters; red), U-Net_Smaller (2.99M; blue), and Shorkie_Random_Init using the U-Net_Small architecture (14.08M; black), compared to Shorkie fine-tuned from the pretrained LM using the same U-Net_Small backbone (14.25M; orange). All models were trained with the same data splits and training protocol. Shaded bands denote variability across cross-validation folds (mean ± s.d.). (A) Validation Pearson's R versus epoch. (B) Validation $R^2$ versus epoch.

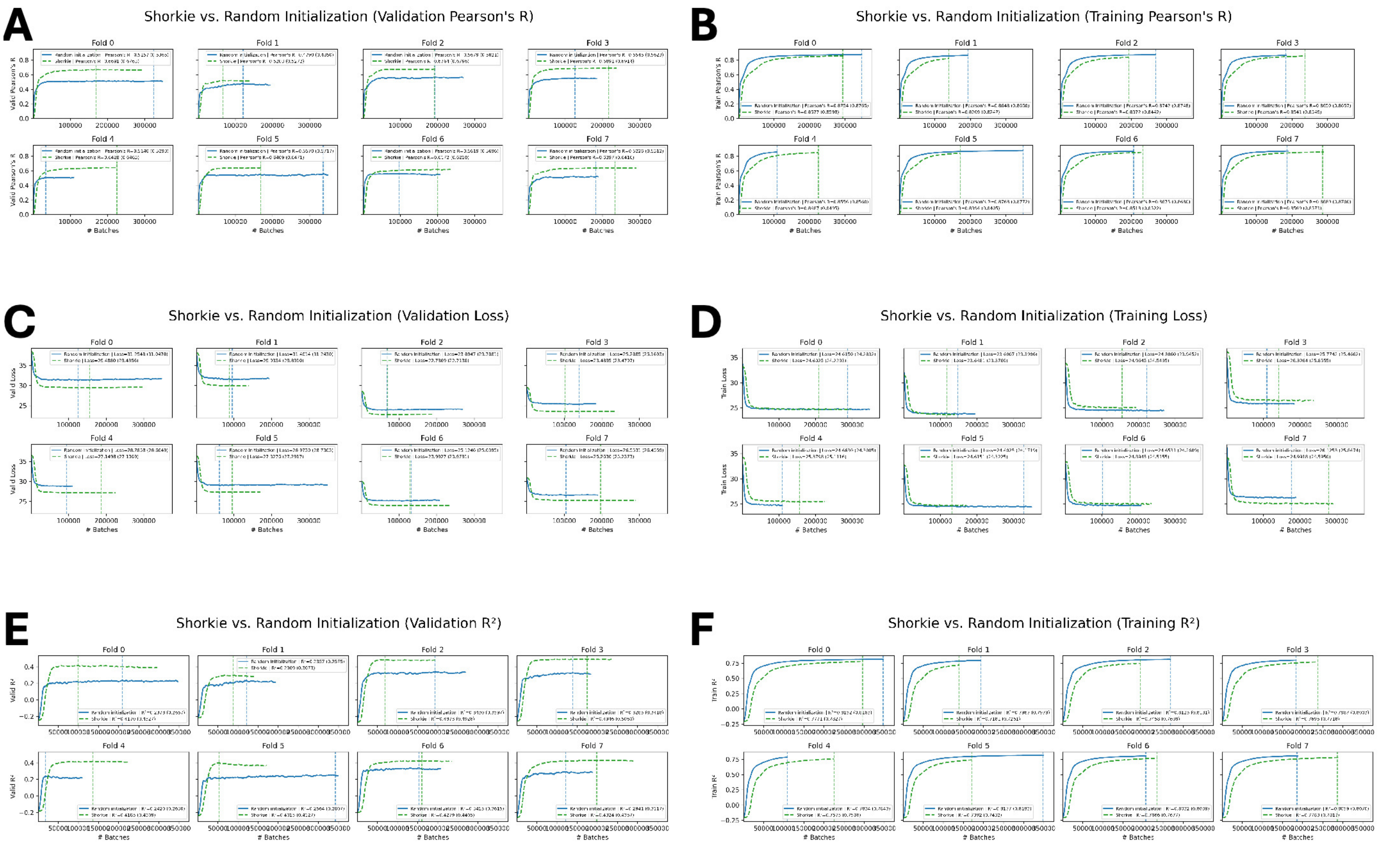


**Figure 3-figure supplement 8.** Training and validation performance of Shorkie and Shorkie_Random_Init. The figure shows six panels displaying the metric trajectories across training epochs for the Shorkie model (initialized with pretrained Shorkie LM weights; dashed lines) and the Shorkie_Random_Init model (initialized with random weights; solid lines). Panels A, C, and E represent validation metrics, while panels B, D, and F correspond to training metrics. **(A)** Validation Pearson's correlation coefficient (R); **(B)** training Pearson's correlation coefficient (R); **(C)** validation loss; **(D)** training loss; **(E)** validation R-squared ($R^2$); and **(F)** training R-squared ($R^2$).

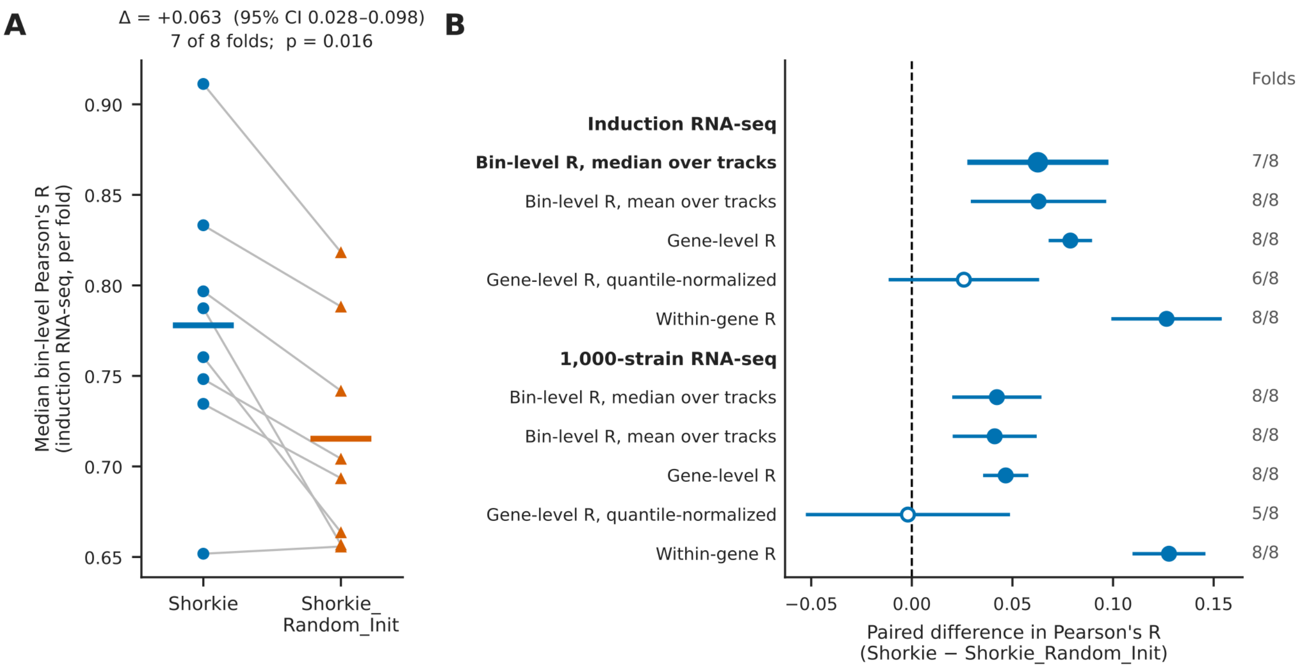


**Figure 3-figure supplement 9.** Cross-fold confidence intervals for the comparison between Shorkie and Shorkie_Random_Init on RNA-seq. Both models were evaluated on the same eight held-out chromosome folds. (A) Median bin-level Pearson's R for induction RNA-seq in each of eight held-out chromosome folds. Grey lines connect matched folds; bars show fold means. Shorkie is higher by +0.063 on average (95% CI 0.028 – 0.098) and in 7 of 8 folds. The paired difference represents the mean of per-fold differences in median R. (B) Paired difference in Pearson's R (Shorkie − Shorkie_Random_Init) for induction and 1,000-strain RNA-seq. Points represent mean paired differences across the eight folds; bars represent 95% confidence intervals; filled points indicate intervals above zero. Labels on the right show the number of folds favoring Shorkie.

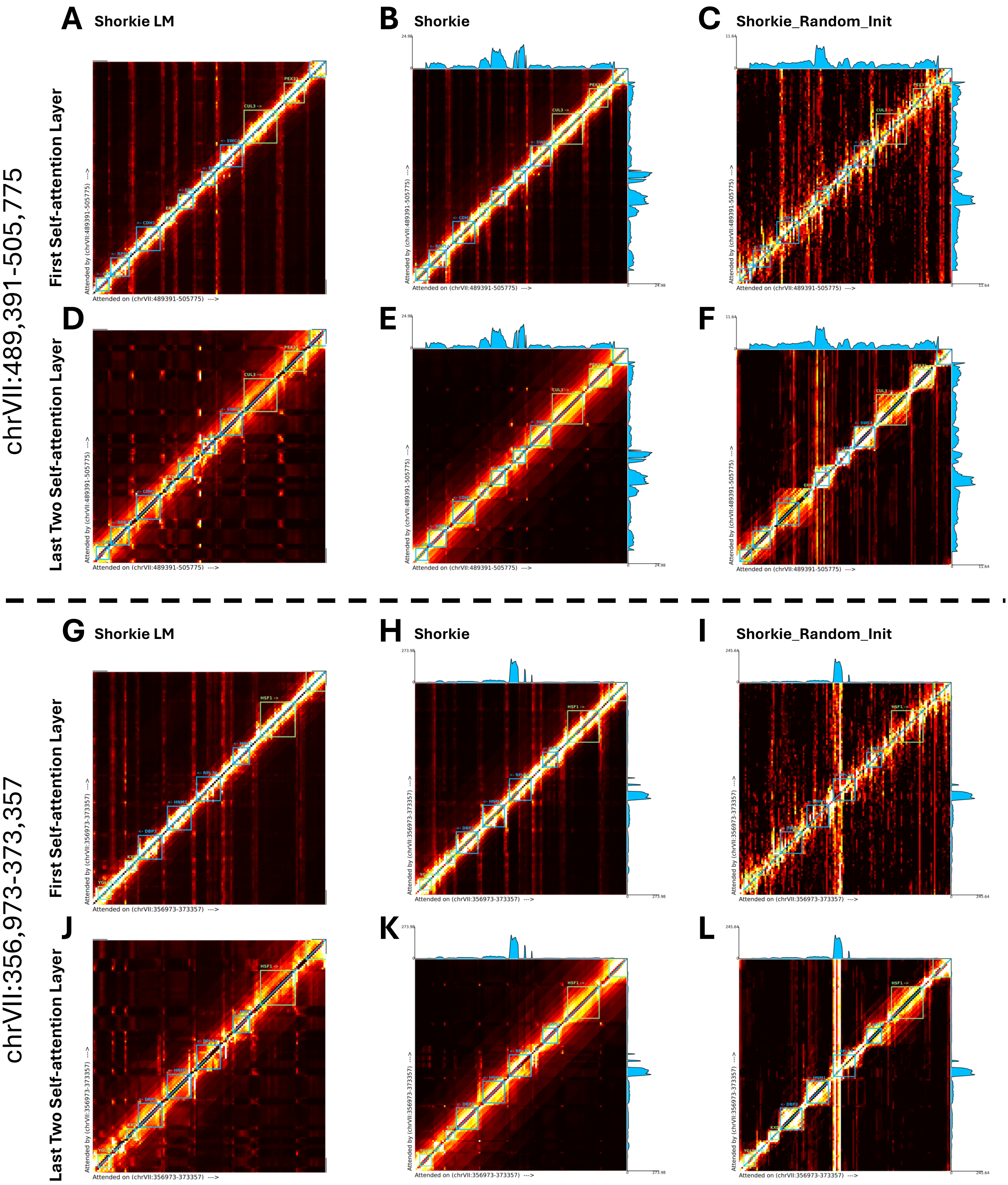


**Figure 3-figure supplement 10.** Self-attention reveals gene-structure encoding and refined regulatory focus. Self-attention scores, averaged across eight heads, are shown for (i) the first transformer block **(A–C, G–I)** and (ii) the last two self-attention blocks **(D–F, J–L)** for each model (Shorkie LM in column 1, Shorkie in column 2, and Shorkie_Random_Init in column 3, over 16,384 bp windows centered on the three-exon gene

**EFM5** (chrVII:489,391–505,775; top two rows) and the housekeeping gene **RPL7A** (chrVII:356,973–373,357; bottom two rows), using $T_0$ RNA-Seq tracks. All models focus attention on exonic segments within gene bodies, indicating robust gene-structure recognition. Shorkie LM and Shorkie additionally exhibit sharp peaks in flanking intergenic regions, whereas Shorkie_Random_Init shows a diffuse intergenic signal. Transfer learning further sharpens Shorkie's regulatory focus: last-layer attention **(D–F, J–L)** becomes tightly localized at specific intergenic loci, while first-layer attention **(A–C, G–I)** remains broadly distributed, reflecting progressive layer-wise specialization.

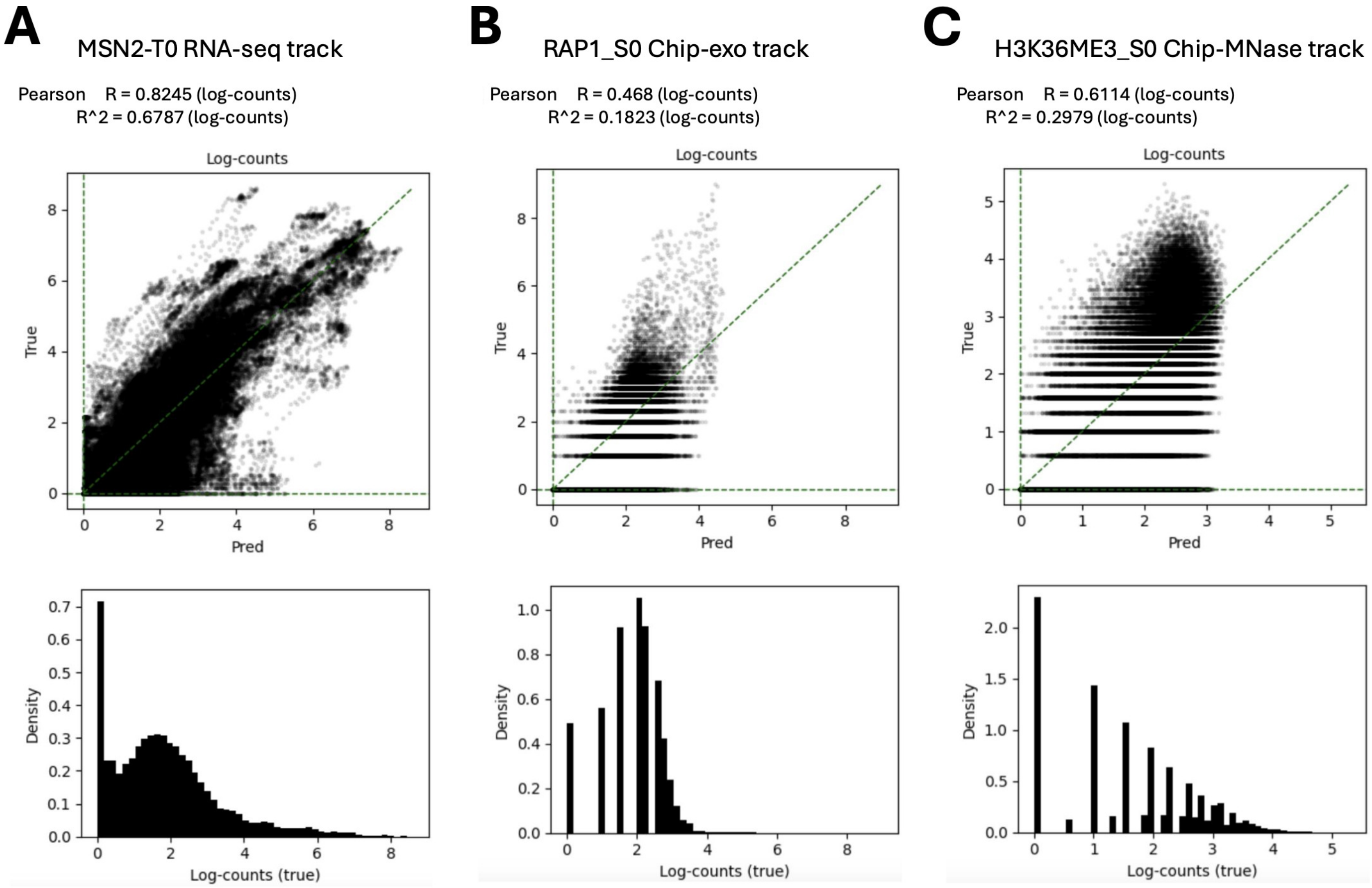


**Figure 3-figure supplement 11.** Bin-level predicted vs. measured signal across assays (fold-0 test set). The upper row shows scatterplots of model predictions (x-axis; log-counts) versus measurements (y-axis; log-counts) for representative tracks: **(A)** MSN2-T0 RNA-seq, **(B)** RAP1_S0 ChIP-exo, and **(C)** H3K36ME3_S0 ChIP–MNase. Each point is a genomic bin; the dashed diagonal marks the $y = x$ line (perfect agreement). Pearson's $R$ and $R^2$ are computed on log-counts and reported in each panel. The bottom row shows histograms of the measured (true) log-counts for the same bins, illustrating assay-specific dynamic range. All examples use held-out regions from the fold-0 model.

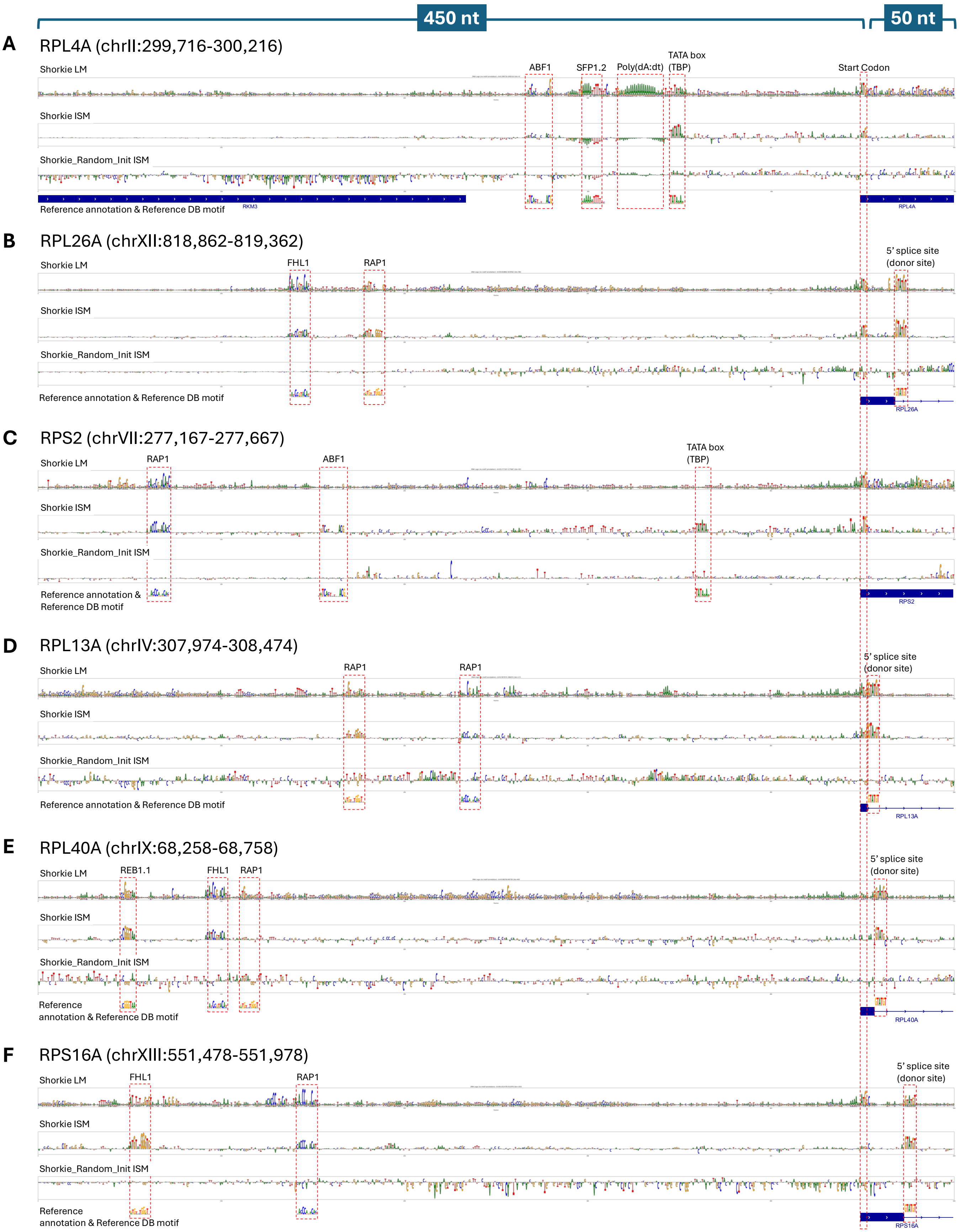


**Figure 4-figure supplement 1.** Motif analysis in promoter regions of six ribosomal protein genes. A 500-bp window spanning –450 to +50 bp around the TSS was extracted for **(A)** RPL4A (chrII:299,716–300,216), **(B)** RPL43B (chrXII:818,862–819,362), **(C)** RPS9A (chrVII:277,167–277,667), **(D)** RPL13A (chrIV:307,974–308,474), **(E)** RPL40A (chrIX:68,258–68,758) and **(F)** RPS16A (chrXIII:551,478–551,978). For each promoter: Row 1 shows conservation-based DNA logos produced by Shorkie LM. Row 2 shows ISM maps from the fine-tuned Shorkie model, where each nucleotide was systematically substituted and the effect on predicted RNA-Seq signal recorded. Row 3 shows ISM maps from Shorkie_Random_Init trained from scratch. Row 4 overlays the reference gene annotation and curated TF-binding motifs from the yeast motif database.

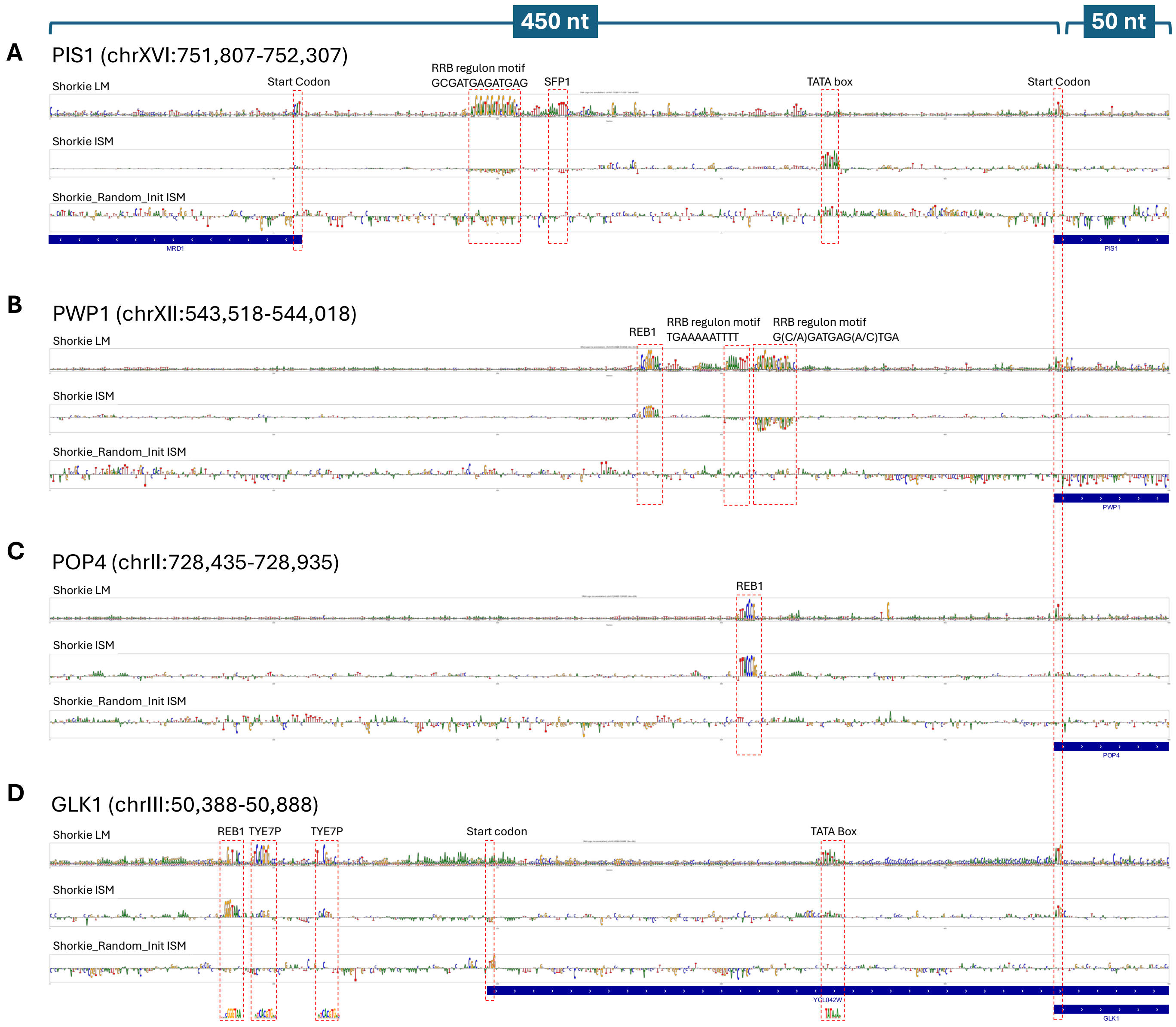


**Figure 4-figure supplement 2.** Motif analysis in promoter regions of four protein-coding genes. A 500-bp window spanning –450 to +50 bp around the TSS was extracted for **(A)** PIS1 (chrXVI:751,807–752,307), **(B)** PWP1 (chrXII:543,518–544,018), **(C)** POP4 (chrII:728,435–728,935), and **(D)** GLK1 (chrIII:50,388–50,888). For each promoter: Row 1 shows conservation-based DNA logos produced by Shorkie LM. Row 2 shows ISM maps from the fine-tuned Shorkie model, where each nucleotide was systematically substituted and the effect on predicted RNA-Seq signal recorded. Row 3 shows ISM maps from Shorkie_Random_Init trained from scratch. Row 4 overlays the reference gene annotation and curated TF-binding motifs from the yeast database, including the RRB regulon motifs (e.g. G(C/A)GATGAG(A/C)TGA and TGAAAAATTTT), REB1, TYE7, TATA box, and the annotated start codon.

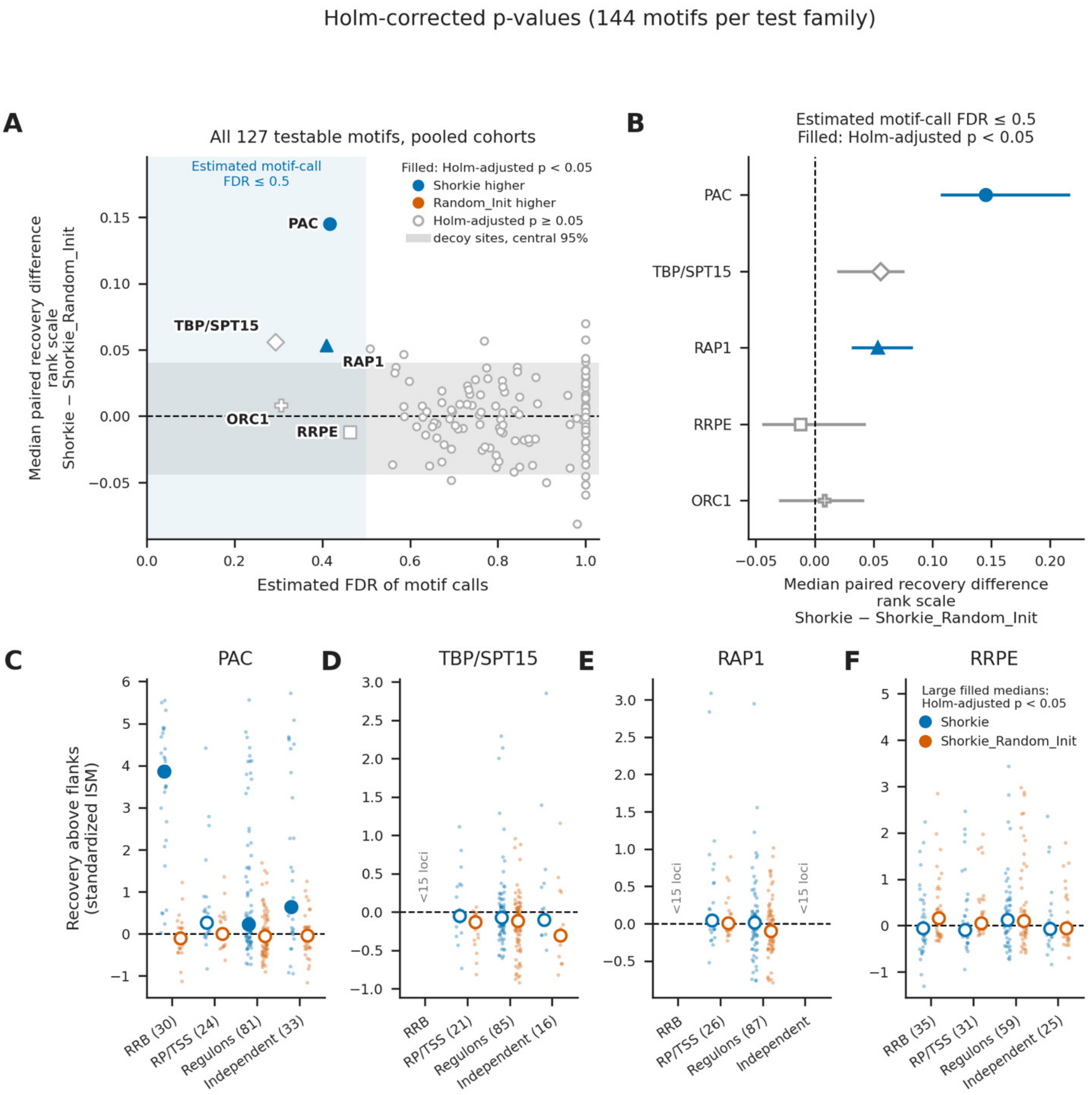


**Figure 4-figure supplement 3.** Quantitative comparison of known-motif recovery in Shorkie and Shorkie_Random_Init ISM maps. ISM maps of both models (fold 0) were compared across 1,685 500-bp promoter windows from the RRB, RP/TSS, regulon and independent cohorts. Recovery is the mean normalized absolute attribution at motif calls minus that in their eligible ±50-bp flanks. (A) Median paired recovery difference (Shorkie − Shorkie_Random_Init) versus estimated motif-call FDR for 127 motifs present in ≥15 loci. Blue shading marks FDR ≤0.5; the grey band spans the central 95% of decoy-set median differences. (B) The five motifs with FDR ≤0.5, with pointwise 95% paired-locus bootstrap intervals (blue, Shorkie higher; orange, Shorkie_Random_Init higher). (C–F) Standardized recovery for PAC, TBP, RAP1 and RRPE, respectively, by cohort (blue, Shorkie; orange, Shorkie_Random_Init; locus count next to each cohort). Small points denote loci; large circles denote medians. Filled circles indicate recovery above flanks for each model (one-sided Wilcoxon tests).

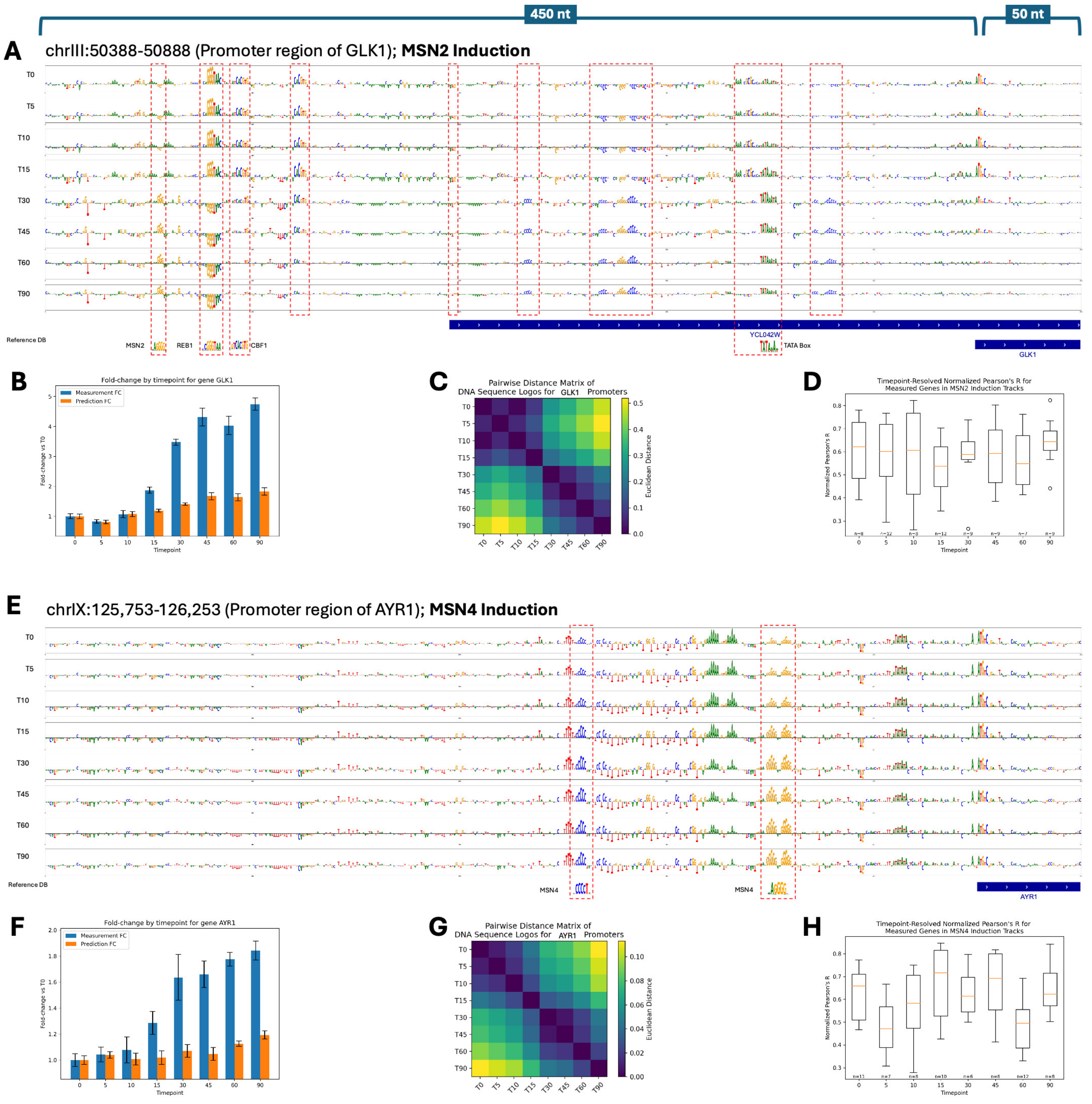


**Figure 5-figure supplement 1.** Time-course analysis of MSN2 and MSN4 induction in *Saccharomyces cerevisiae*. **(A–D)** MSN2 induction at the GLK1 promoter (chrIII:50,388–50,888): (A) Shorkie-generated ISM maps for the GLK1 promoter at eight timepoints following MSN2 induction (T0 = 0 min; T5 = 5 min; T10 = 10 min; T15 = 15 min; T30 = 30 min; T45 = 45 min; T60 = 60 min; T90 = 90 min). Rows correspond to successive timepoints, with the bottom row showing the reference sequence. Key motifs (TATA box, Cbf1, MSN2, Reb1) are annotated. **(B)** Experimental RNA-Seq fold change relative to T0 (blue) versus Shorkie predictions (orange) across the GLK1 locus. **(C)** Heatmap of pairwise Euclidean distances between the ISM maps for all MSN2 induction timepoint pairs. **(D)** Boxplot of normalized Pearson's R scores between experimental and predicted profiles across all yeast genes under MSN2 induction at each timepoint. **(E–H)** MSN4 induction at the AYR1 promoter (chrIX:125,753–126,253), with panels analogous to **A–D**: **(E)** ISM maps generated by Shorkie for the AYR1 promoter across the same eight timepoints.

**(F)** Experimental versus predicted fold change at different timepoints over the AYR1 locus. **(G)** Pairwise Euclidean distance heatmap of the AYR1 logos. **(H)** Boxplot of normalized Pearson's R scores for MSN4-induced samples at each timepoint.

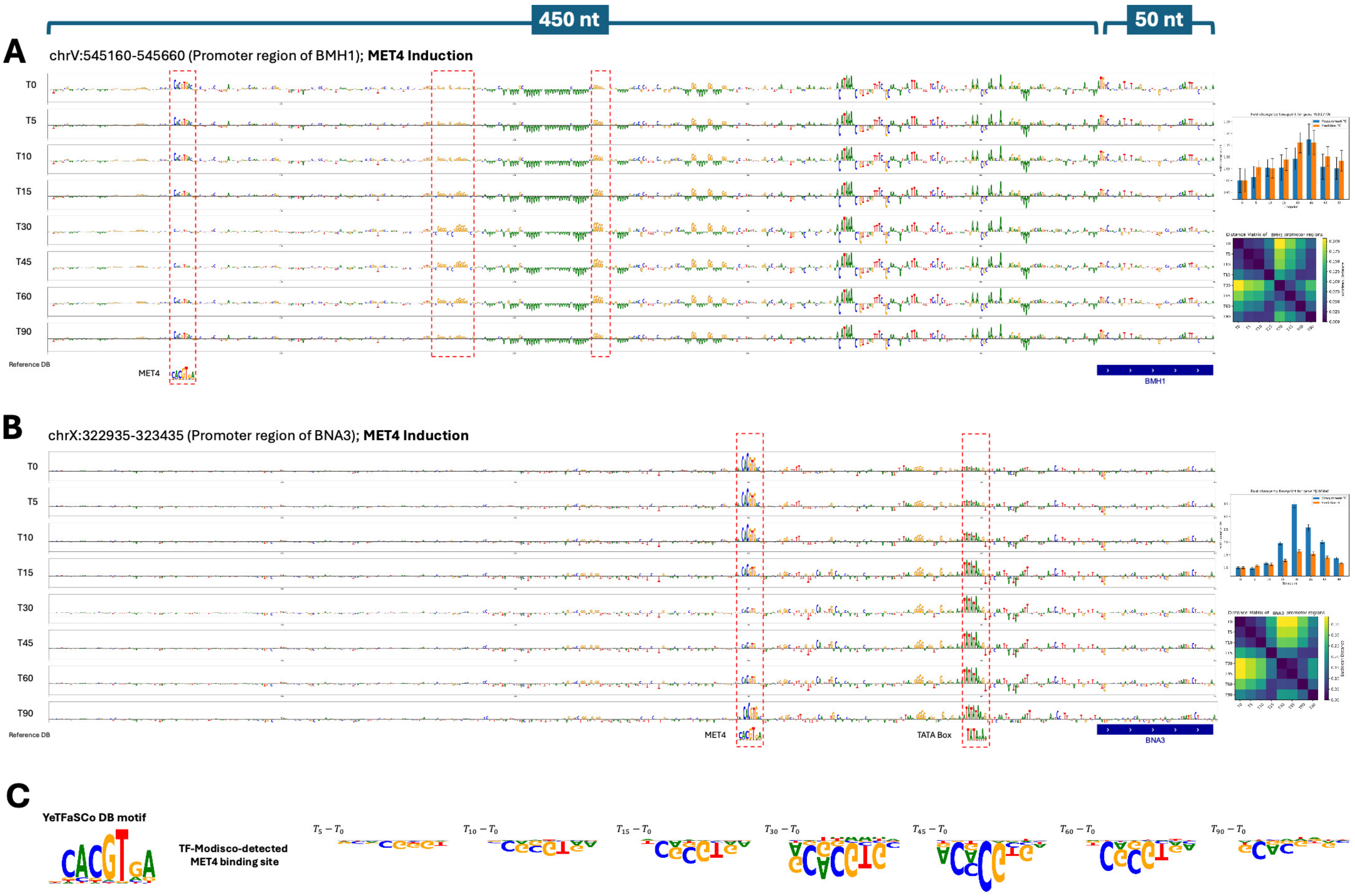


**Figure 5-figure supplement 2.** Time-course analysis of MET4 induction in *Saccharomyces cerevisiae*. Shorkie ISM was applied to 500 bp windows (–450 to +50 bp relative to the transcription start site, TSS) of the BMH1 (chrV:545,160–545,660) and BNA3 (chrX:322,935–323,435) promoters during MET4 induction. **(A)** ISM heatmaps for the BMH1 promoter at eight timepoints (T0 = 0 min, T5 = 5 min, T10 = 10 min, T15 = 15 min, T30 = 30 min, T45 = 45 min, T60 = 60 min, T90 = 90 min). Each row represents a logo at a single timepoint, with the bottom row showing the reference annotation and curated *Saccharomyces cerevisiae* transcription factor (TF)-binding motifs. Key regulatory elements, including the MET4 binding site and TATA box, are annotated. **(B)** ISM heatmaps for the BNA3 promoter, plotted as in **(A)**. **(C)** Consensus MET4 binding motif derived from TF-MoDISco, generated by clustering differential ISM scores across all timepoints relative to T0.

**A**

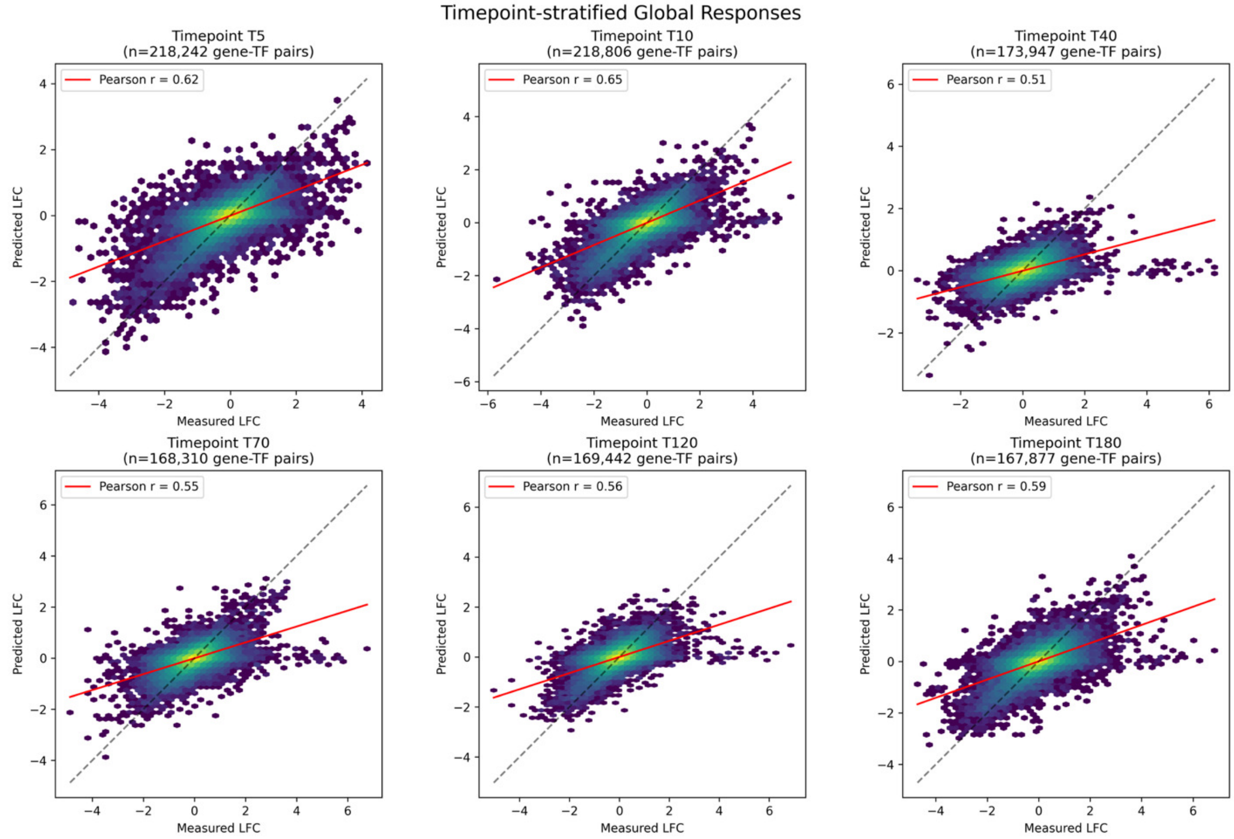
Timepoint-stratified Global Responses
Timepoint T5
(n=218,242 gene-TF pairs)
Pearson r = 0.62
Timepoint T10
(n=218,806 gene-TF pairs)
Pearson r = 0.65
Timepoint T40
(n=173,947 gene-TF pairs)
Pearson r = 0.51
Timepoint T70
(n=168,310 gene-TF pairs)
Pearson r = 0.55
Timepoint T120
(n=169,442 gene-TF pairs)
Pearson r = 0.56
Timepoint T180
(n=167,877 gene-TF pairs)
Pearson r = 0.59
Predicted LFC
Measured LFC

**B**

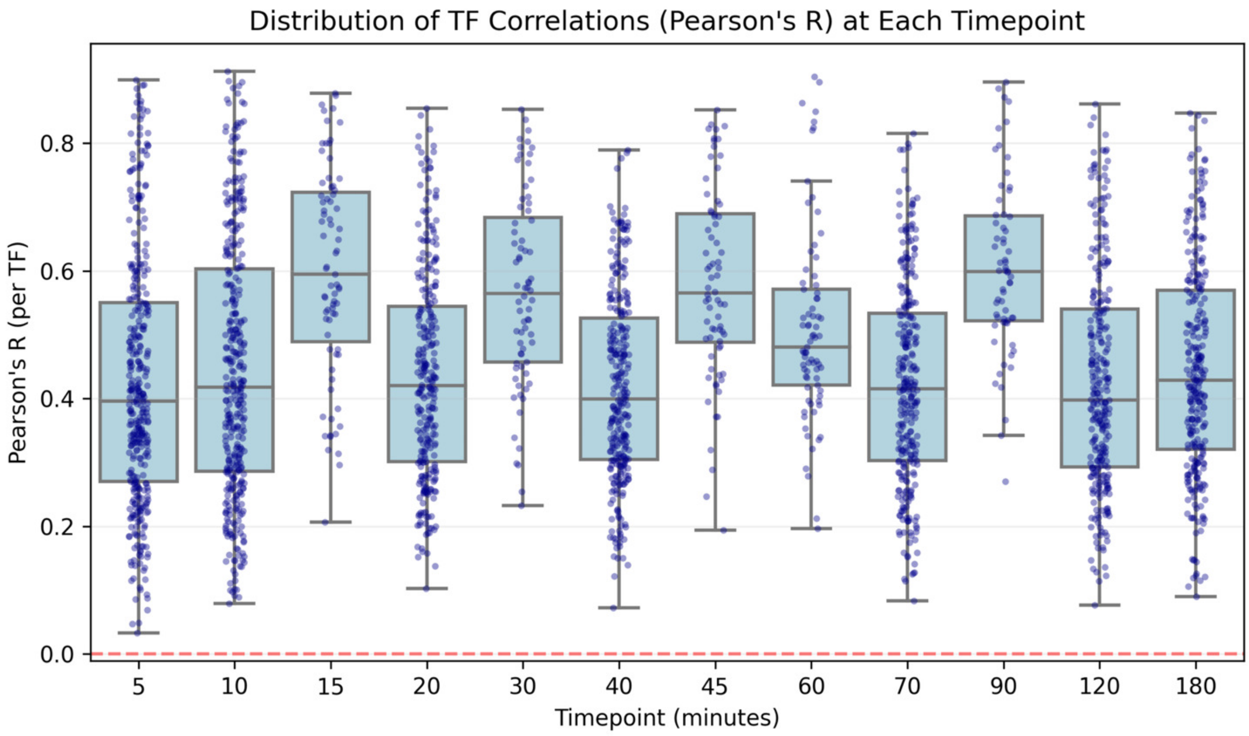
Distribution of TF Correlations (Pearson's R) at Each Timepoint
Pearson's R (per TF)
Timepoint (minutes)
5
10
15
20
30
40
45
60
70
90
120
180

**Figure 5-figure supplement 3.** Stratified global evaluation of predicted versus measured dynamic transcriptional responses. **(A)** High-density hexbin scatter plots of model-predicted versus experimentally measured RNA-seq $\log_2$ fold-changes (LFC) relative to baseline (T0). Data are stratified by progression timepoint, assessing up to 218,806 unique gene-TF induction pairs per temporal slice. A single "gene-TF pair" represents the dynamic LFC of one specific target gene (~6,000 genome-wide) under the induction of one specific transcription factor (>150 evaluated). The red line represents the linear regression fit (Pearson's R), with the dashed line indicating the $y = x$ axis of identity. **(B)** Distribution of Pearson's R coefficients verifying predictive performance across individual TFs over time. Each data point denotes the genome-wide correlation achieved by the model for a single induced TF.

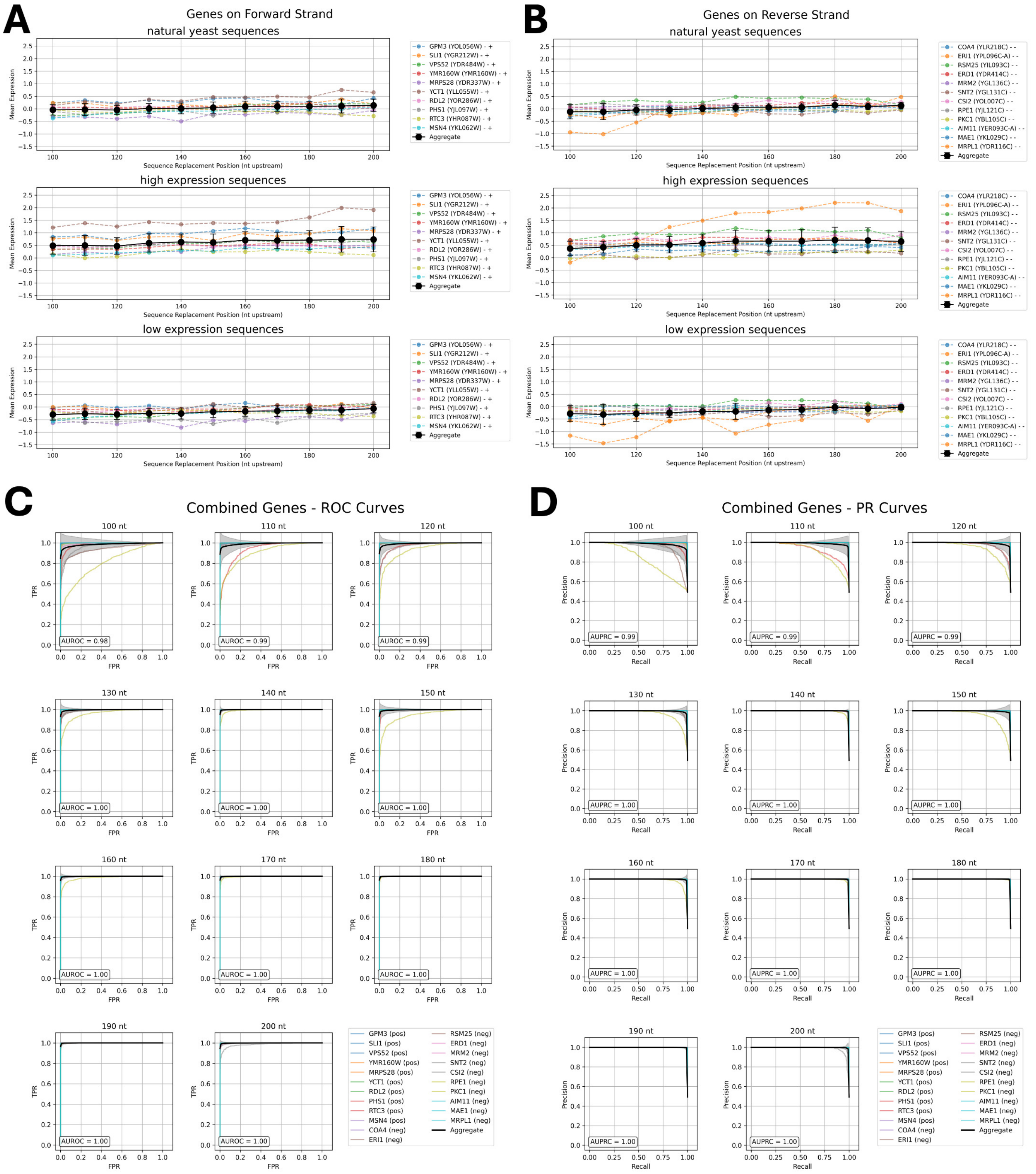


**Figure 6-figure supplement 1.** Shorkie prediction of cis-regulatory activity in a yeast MPRA assay. **(A–B)** Mean logSED scores predicted by Shorkie for native *S. cerevisiae* promoter sequences and for the highest- and lowest-expressing variants from the DREAM challenge, plotted at each nucleotide position relative to the transcription start site (TSS). **(A)** shows data for 10 genes on the forward strand, and **(B)** shows data for 12 genes on the reverse strand. Dashed colored lines represent the per-gene mean logSED across ~1,000 sequence variants at each position, while solid black lines indicate the overall mean ± standard error across all 22 genes. **(C–D)** Performance of Shorkie in classifying high- versus low-expression variants from 100 to 200 bp upstream of the TSS, evaluated by **(C)** receiver operating characteristic (ROC) analysis and **(D)** precision–recall (PR) analysis. Gene-specific curves are shown in color, with the overall mean curves shown in black.

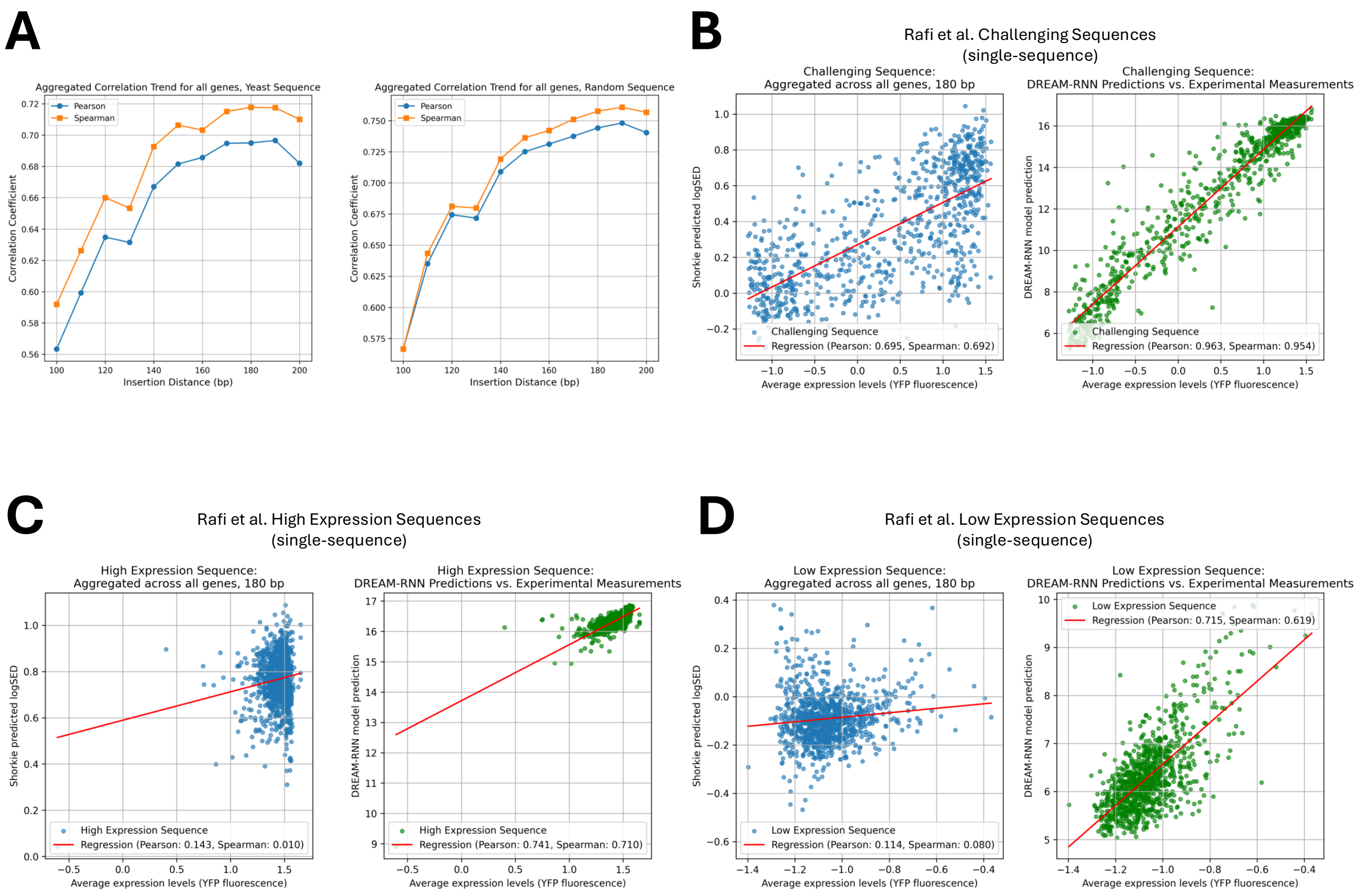


**Figure 6-figure supplement 2. (A)** Pearson's correlation coefficient (R) between Shorkie-predicted logSED scores at various positions upstream of the transcription start site (TSS) and mean YFP fluorescence levels measured by MPRA. Left: native yeast promoter sequences; right: randomized sequences. **(B–D)** Scatterplots of predicted logSED versus observed YFP expression for three distinct sequence subsets: **(B)** challenging constructs, **(C)** high-expression constructs, and **(D)** low-expression constructs. Each plot compares predictions from the Shorkie model (left) and the DREAM-RNN model (right).

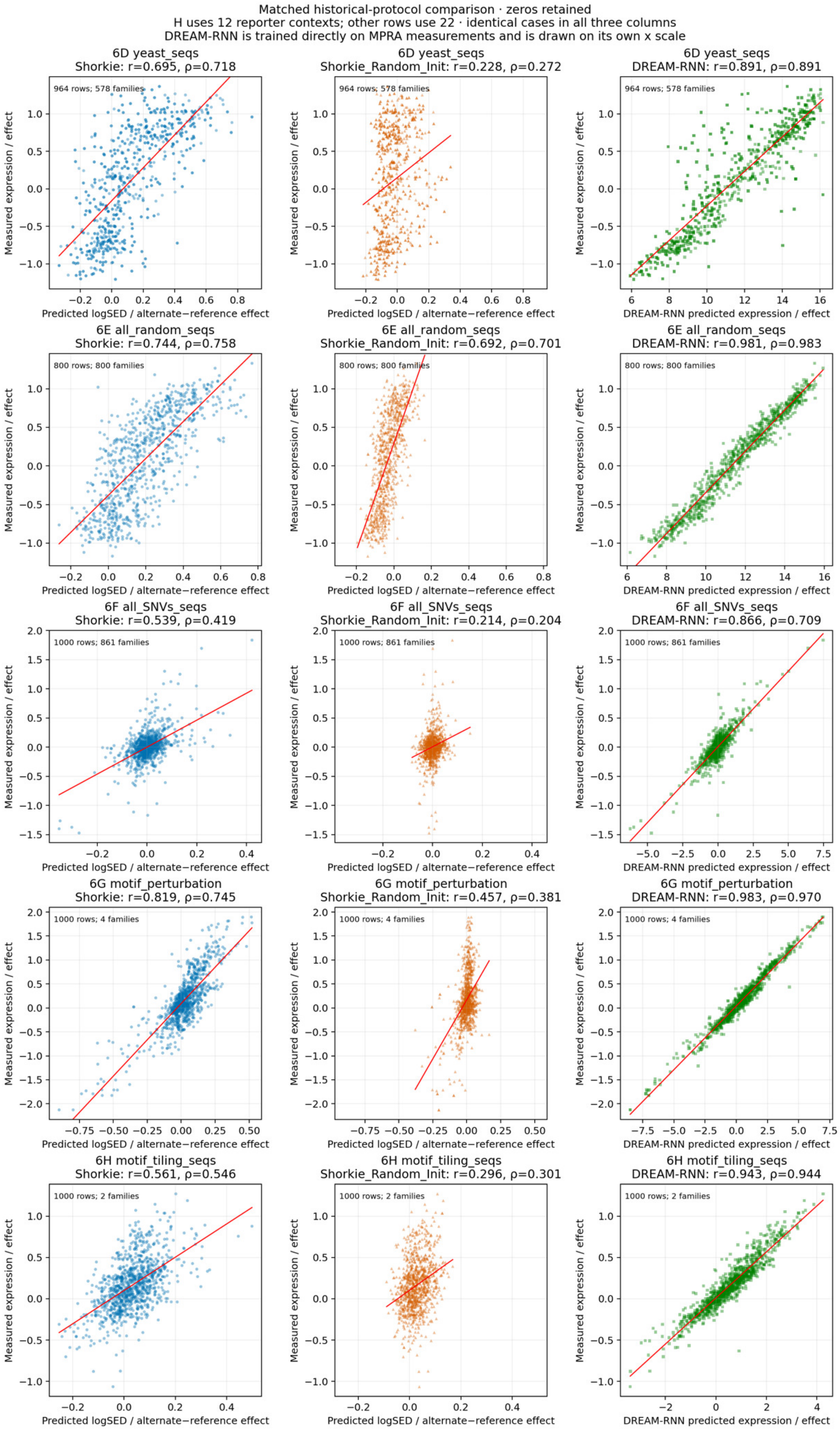
Matched historical-protocol comparison · zeros retained
H uses 12 reporter contexts; other rows use 22 · identical cases in all three columns
DREAM-RNN is trained directly on MPRA measurements and is drawn on its own x scale
6D yeast_seqs
Shorkie: r=0.695, ρ=0.718
6D yeast_seqs
Shorkie_Random_Init: r=0.228, ρ=0.272
6D yeast_seqs
DREAM-RNN: r=0.891, ρ=0.891
964 rows; 578 families
6E all_random_seqs
Shorkie: r=0.744, ρ=0.758
6E all_random_seqs
Shorkie_Random_Init: r=0.692, ρ=0.701
6E all_random_seqs
DREAM-RNN: r=0.981, ρ=0.983
800 rows; 800 families
6F all_SNVs_seqs
Shorkie: r=0.539, ρ=0.419
6F all_SNVs_seqs
Shorkie_Random_Init: r=0.214, ρ=0.204
6F all_SNVs_seqs
DREAM-RNN: r=0.866, ρ=0.709
1000 rows; 861 families
6G motif_perturbation
Shorkie: r=0.819, ρ=0.745
6G motif_perturbation
Shorkie_Random_Init: r=0.457, ρ=0.381
6G motif_perturbation
DREAM-RNN: r=0.983, ρ=0.970
1000 rows; 4 families
6H motif_tiling_seqs
Shorkie: r=0.561, ρ=0.546
6H motif_tiling_seqs
Shorkie_Random_Init: r=0.296, ρ=0.301
6H motif_tiling_seqs
DREAM-RNN: r=0.943, ρ=0.944
1000 rows; 2 families
Measured expression / effect
Predicted logSED / alternate−reference effect
DREAM-RNN predicted expression / effect

**Figure 6-figure supplement 3.** Shorkie, Shorkie_Random_Init and DREAM-RNN on the DREAM MPRA benchmark. Predicted versus measured expression for the five categories of Figure 6D–H (rows) and three models (columns). Shorkie and Shorkie_Random_Init predictions are log fold-change scores from in silico MPRA construct insertions 180 bp upstream of the TSS across 22 selected genes, averaged over eight cross-validation folds.

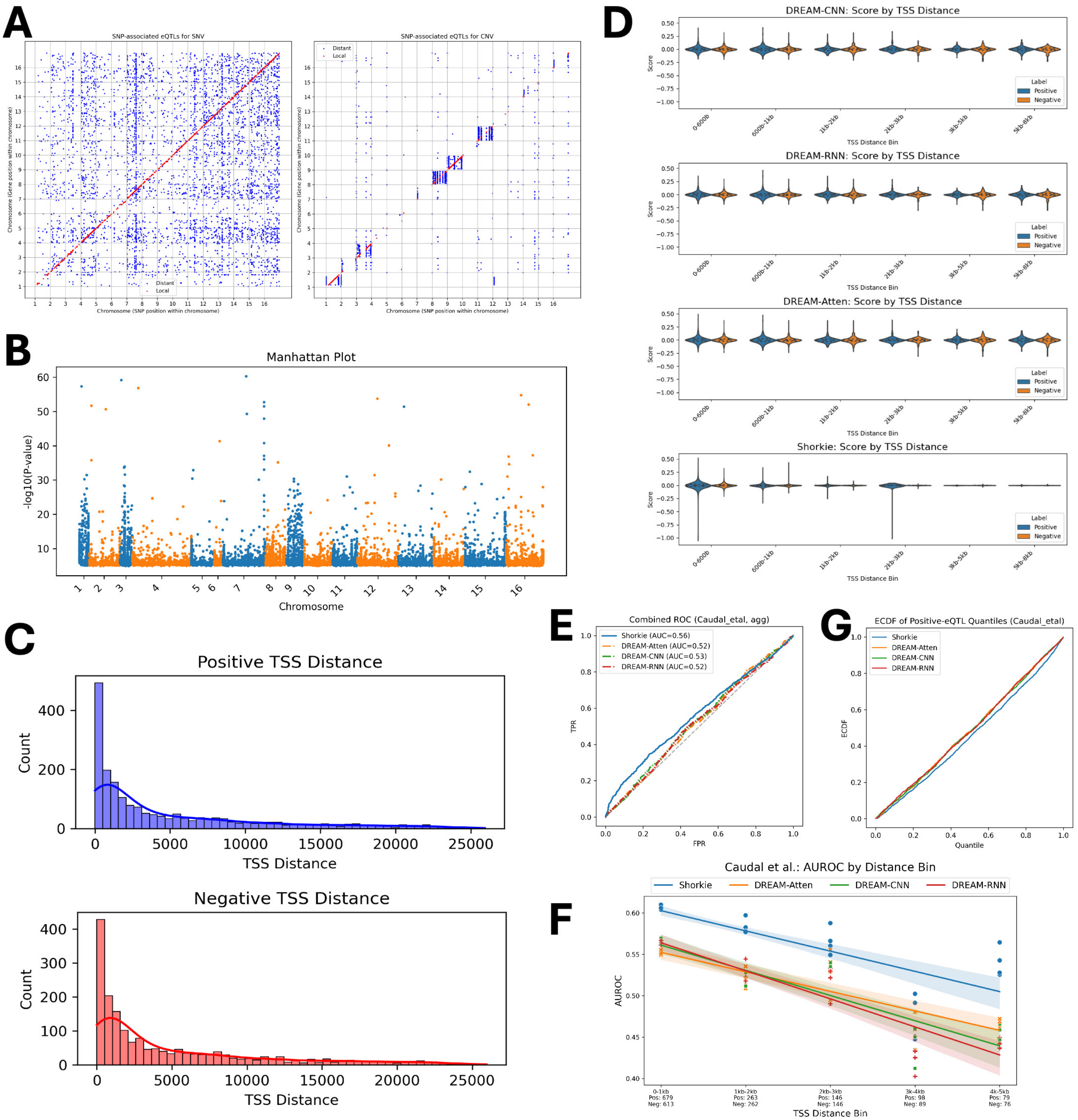


**Figure 7-figure supplement 1.** Predictions of variant effects on the Caudal et al. eQTL dataset by Shorkie and the DREAM Challenge models. **(A)** Genomic locations of SNP-associated eQTLs (left) and copy-number variant (CNV)-associated eQTLs (right), differentiated by local (red) and distant (blue) effects, adapted from Caudal et al. (2024). **(B)** Manhattan plot displaying all SNP-associated eQTLs across the Saccharomyces cerevisiae genome. **(C)** Distribution of distances from eQTL variants (both positive and negative) to their target gene transcription start sites (TSS). **(D)** Score distributions for positive and negative eQTLs as predicted by Shorkie and the top three models from the DREAM Challenge (DREAM-CNN, DREAM-RNN, DREAM-Atten; Rafi et al., 2024). **(E)** Receiver operating characteristic (ROC) curves comparing the ability of each model to classify positive versus negative eQTLs. **(F)** Area under the

ROC curve (AUROC) stratified by TSS distance bins, highlighting how classification performance varies with genomic distance. **(G)** Empirical cumulative distribution functions (ECDFs) of positive-eQTL quantile scores for Shorkie and the DREAM Challenge models, where a steeper rise at lower quantiles indicates stronger enrichment of high-effect predictions among true positives.

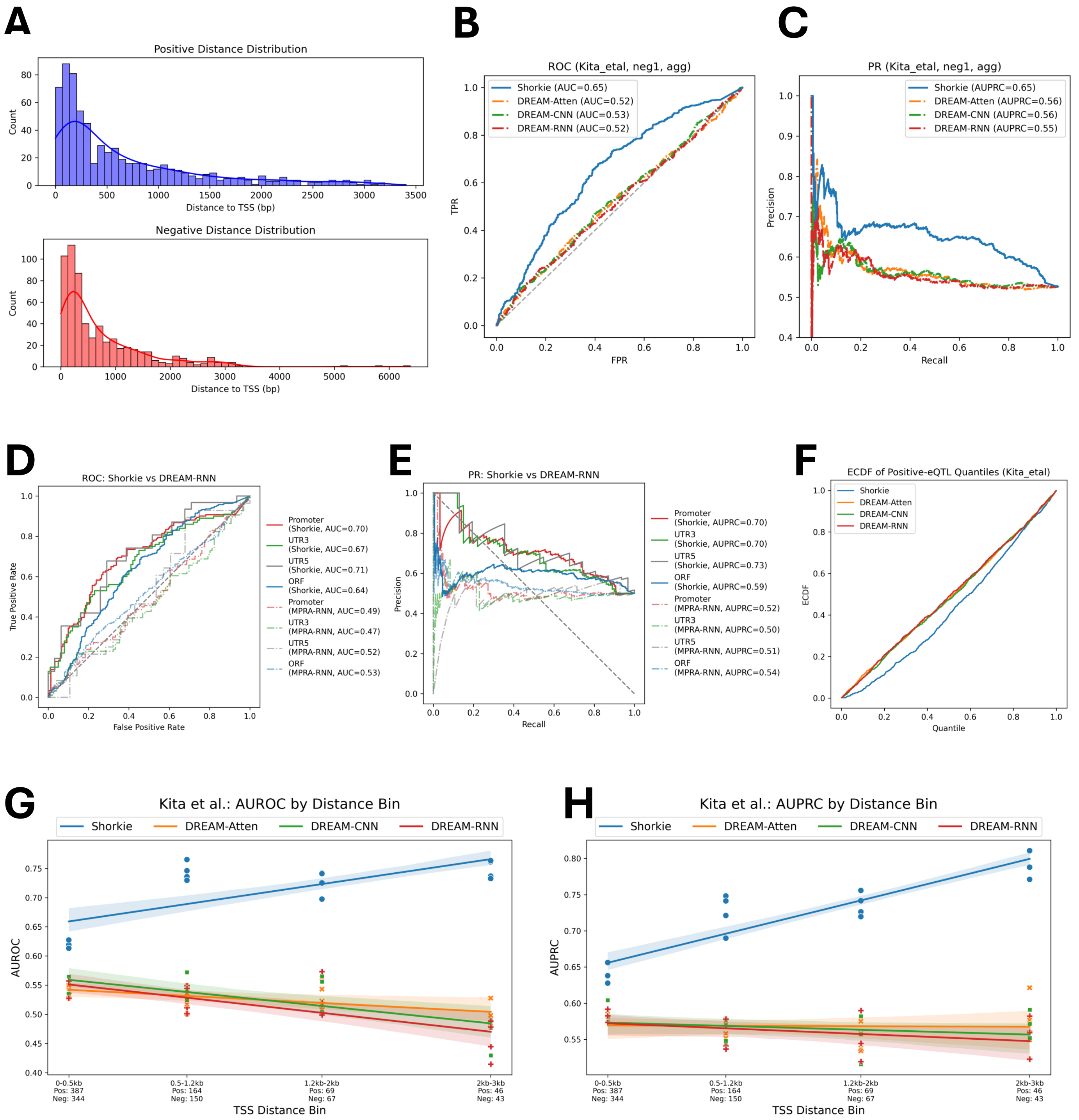


**Figure 7-figure supplement 2.** Evaluation of variant effect predictions on the Kita et al. eQTL dataset using Shorkie and DREAM Challenge models. **(A)** Distribution of distances from positive and negative eQTL variants to the transcription start site (TSS) of their target genes. **(B)** Receiver Operating Characteristic (ROC) curves and **(C)** Precision-Recall (PR) curves comparing the performance of Shorkie and DREAM Challenge models in distinguishing positive from negative eQTLs. **(D)** ROC and **(E)** PR curves for eQTL SNP locations: (1) promoter, (2) 3' UTR, (3) 5' UTR, and (4) open reading frame (ORF), comparing Shorkie with DREAM-RNN. **(F)** Empirical Cumulative Distribution Functions (ECDFs) of positive eQTL quantile scores for Shorkie and DREAM Challenge models. A steeper rise at lower quantiles indicates a higher enrichment of high-effect predictions among true positives. **(G)** Area under the ROC curve (AUROC) and **(H)** Area under the PR curve (AUPRC) stratified by TSS-distance bins, illustrating the variation in model classification performance relative to genomic distance.

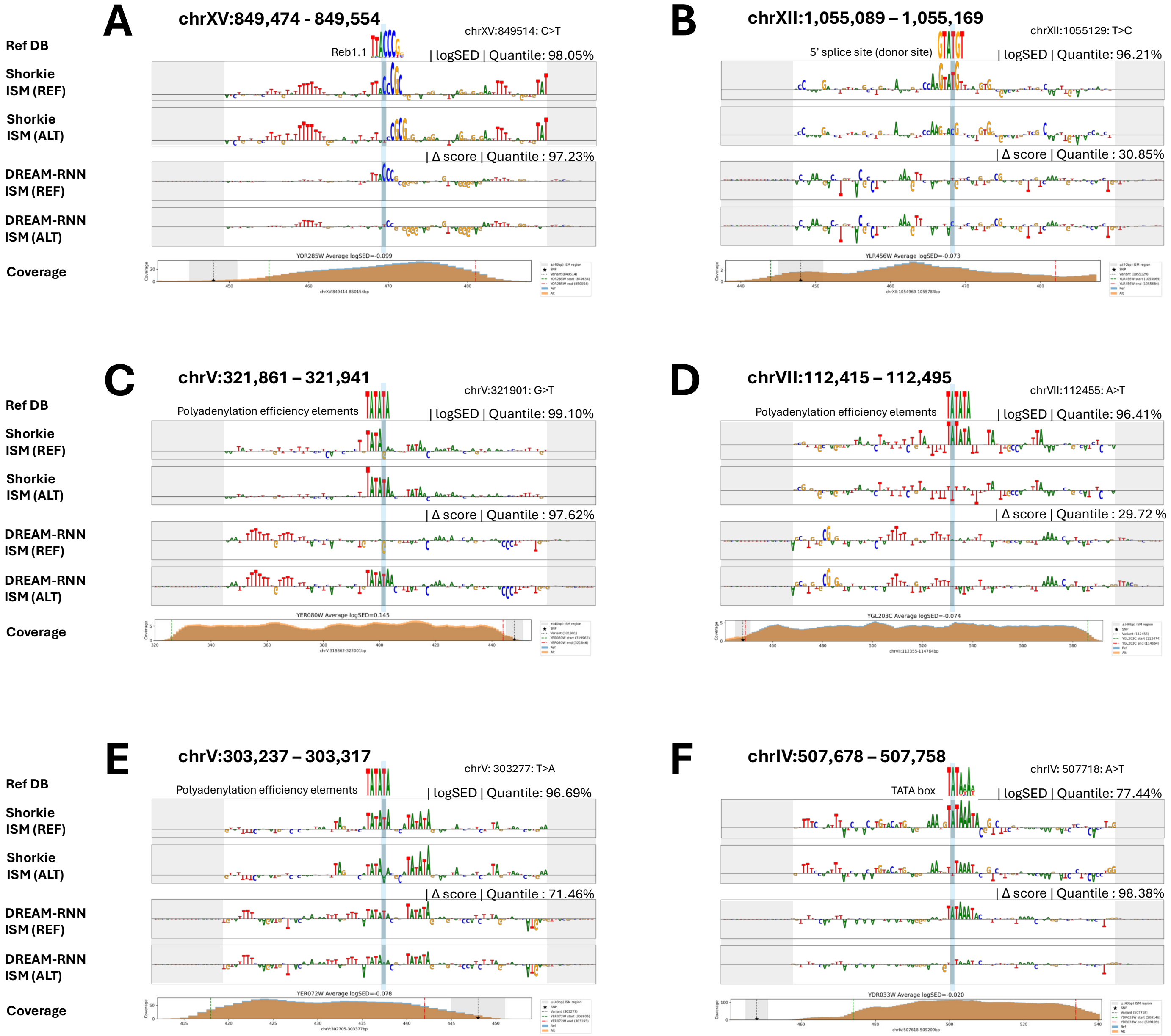


**Figure 7-figure supplement 3.** Shorkie *in silico* mutagenesis (ISM) analysis of predicted variant effects at eight eQTL loci. Shown are Shorkie ISM maps alongside DREAM-RNN Δ-score profiles for the reference (REF) and alternative (ALT) alleles, each plotted over an 80 bp window centered on the variant. Negative scores indicate a predicted decrease in expression for the ALT allele relative to REF, while positive scores indicate an increase. Below each panel, the exact genomic span (chromosome:start–end) and coverage (read depth) are provided.
